# Pervasive GxE interactions shape adaptive trajectories and the exploration of the phenotypic space in artificial selection experiments

**DOI:** 10.1101/2023.01.13.523786

**Authors:** Arnaud Desbiez-Piat, Adrienne Ressayre, Elodie Marchadier, Alicia Noly, Carine Remoué, Clémentine Vitte, Harry Belcram, Aurélie Bourgais, Nathalie Galic, Martine Le Guilloux, Maud I. Tenaillon, Christine Dillmann

**Author notes:** Corresponding authors: Maud.Tenaillon[at]inrae.fr and Christine.Dillmann[at]inrae.fr, Université Paris-Saclay, INRAE, CNRS, AgroParisTech, GQE - Le Moulon, 91190, Gif-sur-Yvette, France. These authors contributed equally to this work (Maud I. Tenaillon, Christine Dillmann).

## Abstract

Quantitative genetics models have shown that long-term selection responses depend on initial variance and mutational influx. Understanding limits of selection requires quantifying the role of mutational variance. However, correlative responses to selection on non-focal traits can perturb the selection response on the focal trait; and generations are often confounded with selection environments so that genotype by environment (GxE) interactions are ignored. The Saclay Divergent Selection Experiments (DSE) on maize flowering time were used to track the fate of individual mutations combining genotyping data and phenotyping data from yearly measurements (DSEYM) and common garden experiments (DSECG) with four objectives (1) to quantify the relative contribution of standing and mutational variance to the selection response, (2) to estimate genotypic mutation effects, (3) to study the impact of GxE interactions in the selection response, (4) to analyze how trait correlations modulate the exploration of the phenotypic space. We validated experimentally the expected enrichment of fixed beneficial mutations with an average effect of +0.278 and +0.299 days to flowering, depending on the genetic background. Fixation of unfavorable mutations reached up to 25% of incoming mutations, a genetic load possibly due to antagonistic pleiotropy, whereby mutations fixed in the selection environment (DSEYM) turned to be unfavorable in the evaluation environment (DSECG). Global patterns of trait correlations were conserved across genetic backgrounds but exhibited temporal patterns. Traits weakly or uncorrelated with flowering time triggered stochastic exploration of the phenotypic space, owing to microenvironment-specific fixation of standing variants and pleiotropic mutational input.

---

Empirical description of phenotypic shifts have nourished quantitative genetics models aiming at deciphering the underlying evolutionary processes (Hill and Caballero 1992; Walsh and Lynch 2018). The persistence through time of heritable variation (Odhiambo and Compton 1987; Moose *et al*. 2004; Weber and Diggins 1990; Caballero *et al*. 1991; Mackay 2010; Lillie *et al*. 2019; Wisser *et al*. 2019) fit well with Fisher’s infinitesimal model (Fisher Ronald Aylmer 1930) and the derivatives of the breeder equation (Lush 1943; Lande 1979; Lande and Arnold 1983). These models indeed predict a continuous and linear response with no finite limits, especially under truncation selection known to be the most effective form of directional selection (Crow and Kimura 1979). However, under the alternative assumption of finite genetic architecture the long-term selection response is expected to plateau, reached all the sooner as selection-induced linkage disequilibrium diminishes the standing genetic diversity, the so-called Bulmer Effect (Bulmer 1971; Hospital and Chevalet 1996). The plateau value, that is the expected maximum shift, depends on both, the initial standing variance and the effective population size (Robertson 1960; Roberts 1966;Falconer 1971). The latter determines the flux of incoming *de novo* mutations. Interestingly, when this flux is large enough, the long-term response to selection can be infinite even with finite genetic architectures (Hill 1982a,b; Lynch and Hill 1986; Weber and Diggins 1990; Wei *et al*. 1996; Walsh and Lynch 2018). Hence quantifying the proportion of genetic variance due to the input of new mutations (*V*_*m*_), the so-called mutational variance, is central to understand the seeming absence of selection limits (Turelli 1984).

Mutational variance is determined by the mutation rate per locus as well as the architecture of the trait — *i*.*e*. the size of the mutational target and the distribution of fitness effects (DFE) of incoming mutations (Bürger 1993; Bürger and Lande 1994). One popular strategy to approach the incoming mutational variance is to rely on divergent selection experiments (DSEs) to estimate the mutational heritability. Hence, starting from a genetically-homogeneous material such as inbred strains, the mutational heritability can be estimated from DSEs, as the slope of the selection response across generations. The mutational heritability denotes the ratio of the mutational variance over the residual variance, 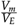 (Hill 1982b; Houle *et al*. 1996; Keightley 2010). Previously estimated values fell within the range 4.10^*−*4^ to 3.10^*−*2^ (Houle *et al*. 1996; Keightley 2010). However, because DSEs offer control over the pedigree of individuals, a classical animal model (Kruuk 2004) can also be applied to decompose phenotypic variance and directly estimate mutational variance considered as a random effect. An interesting aspect of the animal model lies in its ability to account for the effects of newly occurring mutations as they increase similarity among individuals and therefore modify the additive genetic variance-covariance matrix (Wray 1990) (used to compute the Best Linear Unbiased Predictor of Mutational effects, subsequently called BLUPM). This is particularly true in populations characterized by a small effective population size, where the effect of increased similarity is magnified.

There are however limitations to the use of selection experiments. The first limitation is that time-series (generations of selection) introduce confounding effects between generational environments and genetic changes because of ubiquitous genotype-by-environment (GxE) interactions (Li *et al*. 2017), *i*.*e*. the genetic variability in plastic responses. Replicated common garden experiments can however provide effective settings to disentangle these two effects. They have been successfully employed to test for local adaptation for life-history traits, phenology and allometric relationships (reviewed in de Villemereuil *et al*. (2016)). The second limitation is that the trait under selection does not evolve independently from other traits, and correlative responses to selection at other traits may perturb selection responses. Such correlative responses may result from linkage disequilibrium between quantitative trait loci (QTL) evolving under drift/selection. They may also result from correlated physiological functions where a first trait governs a second one, e.g., flowering time in maize is partly determined by floral transition time. Finally, genetic correlations may result from true pleiotropy when a single QTL affects several traits (Chen and Lübberstedt 2010; Reinert 2022).

The evolution of a population in the multivariate phenotypic space is governed, in the short-term, by the extension of the breeder’s equation 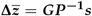 (Lande 1979; Lande and Arnold 1983) linking the vector of mean response **Δ*z*** to the ***G***-matrix. The ***G***-matrix describes the standing additive genetic variances and covariances for a suite of traits, ***s*** being the vector of selection differentials and ***P*** being the observed phenotypic matrix of measured traits. However, in the long-term, the effects of new mutations on genetic variances and covariances –represented through the ***M***-matrix–, define the evolvability of the population (*e*.*g*. see Houle *et al*. (2017)). In the context of an adaptive walk towards a fitness optimum, Fisher’s geometric model predicts that the size of adaptive mutations tends to decrease with the number of influenced traits (reviewed in (Tenaillon 2014)). Consequently the expected DFE is exponential in the case of an infinite number of correlated traits influencing fitness, and follows a beta distribution when the number of correlated traits is small to moderate (Martin and Lenormand 2008). Correlative response to stabilizing selection arises because of an interplay of pleiotropic effects of mutations and correlational selection, favoring certain combinations of trait values. This interplay is however modulated by the intensity of genetic drift: in infinite populations, genetic correlations are unaffected by the strength of selection while in small populations the strength of selection does affect genetic correlations (Chantepie and Chevin 2020; Doroszuk *et al*. 2008).

Here we used the Saclay DSEs on maize flowering time that display significant selection responses over 18 generations -despite a dearth of initially standing variation and a high drift-high selection regime (Durand *et al*. 2010, 2012, 2015; Desbiez-Piat *et al*. 2021) -with four main objectives: (1) to quantify the relative contribution of standing variation and *de novo* mutational variance to the selection response using the BLUPM model (Wray 1990); (2) to estimate the DFE of selected *de novo* mutations; 3) to study the impact of GxE interactions in the observed selection response; (4) to describe correlations between traits and how they modulate the exploration of the phenotypic space. To fulfill these objectives, we combined yearly measurements of flowering time, common garden experiments on a subset of 308 genotypes measured during two consecutive years for eleven traits, as well as genotyping data to follow the fate of individuals throughout pedigrees.

## Materials and methods

### Saclay DSEs

Two independent Divergent Selection Experiments (DSEs) for flowering time, have been conducted on Saclay’s plateau since 1993 (corresponding to 18 generations), from two independent genetic backgrounds: an early-flowering American Dent, F252 registered in 1979 by the company Agri-Obtention, and a late Io-dent dent, MBS847 (thereafter called MBS), registered by the company Mike Brayton Seeds in 1982. Estimates from the genome-wide genotyping of parent lines F252 and MBS with the 50K maize SNP microarray (Bouchet *et al*. 2013; Durand *et al*. 2015) revealed a residual heterozygosity of about 1.9 % for F252, and 0.19 % for MBS over 29,000 SNPs mainly concentrated in genic regions. The selection scheme has previously been described in Durand *et al*. (2010, 2015); Desbiez-Piat *et al*. (2021). Briefly, within each DSE, the ten earliest (resp. ten latest) flowering individuals were selfed at each generation, producing each 100 offspring used for the next generation of selection within the Early (resp. Late) populations (Fig. S1). Within each population, we evaluated 100 offspring of a given genotype in four rows of 25 plants randomly distributed in a four-block design, so that each block contained 10 rows. We applied a truncation selection of 10/1000=1%. Following Durand *et al*. (2015); Desbiez-Piat *et al*. (2021), we designated as *progenitor*, a selected plant represented by its progenies produced by selfing and evaluated in the experimental design at the next generation. We conditioned selection on the maintenance of two families, *i*.*e*. two sub-pedigrees derived from two separate *G*_0_ ancestors. Thus, each family was composed of three to seven progenitor at each generation with the additional condition that at least two different *G*_*n−*1_ progenitors were represented. Furthermore we applied a two-steps selection procedure. First, among the 100 offspring per progenitor, we recorded the flowering time of the 12 most extreme (most early/late) individuals per (early/late) family, *i*.*e*. 12*×*5 = 60 individuals per family or 120 per population. Records constituted the Saclay DSEs Yearly Measurements (thereafter DSEYM). Second, to ensure the maintenance through time of minimal fitness, we selected based on an index among the 12*×*5 earliest (resp. latest) individuals, the 5 earliest (resp. latest) individuals with the highest kernel weight per family. Seeds from selected genotypes at all generations were stored in cold chambers.

F252 and MBS genealogies can be traced back from generation 18 to the start of the divergent selection experiments, G0 (Fig. S2 and Fig. S3). The initial MBS pedigrees encompassed four families: ME1, ME2 composing the MBS Early (ME) popula-tion, and ML1, ML2 composing the MBS Late (ML) population. Likewise F252 Early (FE) population was composed of FE1 and FE2 families. F252 Late populations genealogies were more complex: FVL families (F252 Very Late in Durand *et al*. (2015)) ended at generation 14 with the fixation of a strong effect allele at the *eIF-4A* gene (Durand *et al*. 2015). To maintain two families in F252 Late population, two families FL2.1 and FL2.2 were further derived from the initial FL2. These two families genealogies are rooted in FL2 from a G3 individual.

### Common garden evaluation in 2018 and 2019 -DSECG

Saclay’s DSEs common garden experiments (DSECG) were conducted in 2018 and 2019 on Saclay’s Plateau. We followed a sampling strategy that allowed for the best estimation of progen-itors breeding values (David *et al*. 2022). Sampled progenitors, encompassing 155 MBS (Fig. S2) and 153 F252 (Fig. S3), were evaluated in 2018 and 2019 except for FVL which was not reevaluated fully in 2019, because the latest generation flowered much later than other progenitors.

All seeds were produced in a nursery in 2016 from 25 S1 progenies of each selected progenitor of the genealogy. S2 seeds, obtained by selfing S1 individuals harvested in bulk, constituted the seed lots used in the evaluation trials. The evaluation trials encompassed six randomized-blocks for F252 S2 seeds and six randomized-blocks for the MBS S2 seeds. Half of the blocks were dedicated to early and the other half to late progenitors. Inside a block, we considered the coordinates described on the one hand by the plots (groups of 14 rows of 25 plants sharing the same X coordinate), and on the other hand by the columns (Y coordinate). Early (resp. Late) flowering progenitors were randomly distributed in 13 (resp. 12) plots of 14 rows of 25 plants. Overall, each progenitor was replicated at least twice in two rows of 25 plants randomly distributed among blocks. Note, that some progenitors were replicated more often (up to four times). One row of 25 plants from each of the two inbred lines F252 or MBS were used as control in each block. Controls were produced from 3 generations of multiplication (selfing) of a single individual from the Generation 1 of the early populations FE1 and ME2 (FE1 G1 and ME2 G1).

A total of 10 traits were recorded during both evaluation years. “Flowering time” (FT) was expressed in Days To Flowering equivalent to a growth at 20°C (DTF therefater) following Parent *et al*. (2010). We defined flowering time of progenitor *Y*_*i*_, represented by a row of 25 plants, as the average flowering date. We recorded flowering time (*t*_*i*_) as the number of plant (*Z*_*i*_) for which silks were visible and pollen covered two third of the tassel. Hence, we actually observed the number of plants that effectively flowered between *t*_*i−*1_ and *t*_*i*_. We therefore estimated 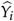 as:

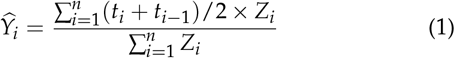

Note that in DSEYM, flowering times were recorded for the 12 most extreme plants only, while in DSECG flowering time of a progenitor was defined as the mean flowering time per row. Hence we did not measure exactly the same trait in the two experiments. However, Durand *et al*. (2015) showed that there was an extremely high correlation between the two measures (r = 0.83 for F252 and r = 0.88 for MBS, p-values *<* 2.2*×*10^*−*16^, in both cases). We therefore used DSEYM to analyze the response to selection over the 18 generations, and compared it to that observed in DSECG.

Moreover, we recorded for 10 plants per row: the plant height (HT) (measured as the distance in cm between ground level and tassel basis), the rank of the upper ear (UE) and of the lower ear (LE), the length in cm of the leaf (LL) blade for the leaf located just below the upper ear, the rank of the upper node with brace roots (BR), the total number of visible leaves (#L). We considered for the subsequent analyses the average over these 10 plants as the phenotypic value. In addition, we recorded the number of visible leafs for these 10 plants every 5 days during the linear part of the vegetative growth. We then computed the phyllochron (PHY) -the time period (in DTF Eq. 20°C) between the sequential emergence of leaves -as the regression slope of the number of leaves over the thermal time during the linear developmental time response. Finally, the thousand kernel fresh weight (TKW in g) and the weight of the upper ear (EW in g) were recorded and averaged across 7 plants per row at complete maturity. Data from F252 and MBS were analysed separately.

### Phenotypic data analyses of Yearly Measurements and Common Garden Evaluation

#### Saclay DSEs yearly measurement (DSEYM) analysis

Following Durand *et al*. (2010, 2015), we corrected the flowering time raw data from DSEYM for experimental design effects (blocks), and used F252 and MBS controls to correct for year effects. We used these estimated genetic values to further decompose the selection responses into two components, one due to initial standing variation and the second due to the incoming mutational variance (Durand *et al*. 2010; Desbiez-Piat *et al*. 2021).

To this end, we used Wray (1990) mixed model approach allowing for both estimation of the initial additive genetic vari-ance 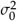 and additive mutational variance 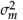. The model can be written:

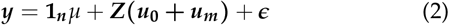

Where, ***y*** is the vector of estimated genetic values after year and block effect correction, **1**_***n***_ is the vector 1 of size *n*, ***u***_**0**_ is a vector of N genetic values that originate from standing variance such that 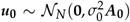, ***u***_***m***_ is a vector of N genetic values that originate from incoming mutational variance with, 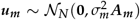, and the residuals are i.i.d and is an incidence matrix. Noteworthy, in ***ϵ*** *∼ 𝒩*_***n***_ (**0**, *σ*^2^ ***I***_***n***_) and ***Z*** this model, there are two kinship matrices, ***A***_**0**_ that describes the kinship among individuals based on the pedigrees and that concerns the standing genetic variation, and ***A***_***m***_ that describes the kinship caused by new mutations arising during the course of the experiment which increases the relatedness among individuals that carry them.

The kinship matrix ***A***_**0**_, or additive genetic relationship matrix (neglecting dominance terms), is computed from pedigree information so that each individual element 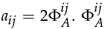 is the kinship coefficient, the probability that an allele randomly drawn for a given gene in individual *i* is identical by descent to a randomly drawn allele from individual *j* at this gene. If *i* = *j*, we can draw twice the same allele (probability 1/2) or the two alleles of *i* (probability 1/2) that are identical by descent in probability *f*_*i*_, the inbreeding coefficient of *i*. Hence, we have: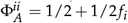 or *a*_*ii*_ = 1 + *f*_*i*_. Recalling that the inbreeding coefficient of an individual is equal to the kinship of its parents, and that under complete selfing, the inbreeding coefficient *f* (*g*) is identical between all individual at generation *g*, we have all progenitors:

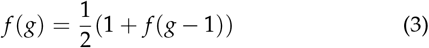

with *f* (0) = 0, i.e. an hypothesis of the BLUPM model is to consider no initial inbreeding from the standing variation point of view, and *a*_*ii*_ (0) = 1, which can be rewritten:

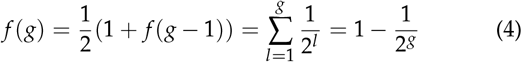

Hence,

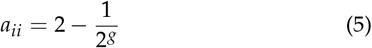

In the same way, for all pairs of individuals (*i, j*), we can easily compute the kinship coefficient from the genealogy. We made the assumptions that:

- when the two focal individuals *i* and *j* did not share a common ancestor, *a*_*ij*_ = 0.
- when the two focal individuals *i* and *j* were from different families (*e*.*g*. FE1 and FE2) and therefore unrelated from a standing variation perspective, *a*_*ij*_ = 0
- when the two focal individuals *i* and *j* within a family had at least one common ancestor,*k* being the most recent, *a*_*ij*_ = *a*_*kk*_ .

In order to account for mutational effects, we made the assumption that mutations occurring at generation *g* are transmitted to their offspring and induce an additional correlation between breeding values at the subsequent generation (Wray 1990), albeit independent from the breeding values at the previous generation. This additional correlation is computed in the same way as for ***A***_**0**_, but there is one matrix per generation, considering that all individuals between generations 0 to *g* are unrelated (because mutations occur independently from each other). Let ***A***_***g***_ be the kinship matrix between *N* progenitors at generation *g* due to mutational effects. We can write:

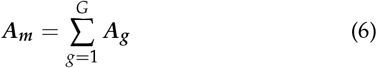

Diagonal elements of ***A***_***g***_ (for *g≥*0) are 1 + *f*_*i*_, where *f*_*i*_ is the inbreeding coefficient of individual i ignoring common ancestors from generations 0 to *g−*1 (Wray 1990). Then, we defined 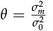, the mutational variance in units of the additive genetic variance in the initial population. Mutational heritability was defined as: 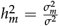, *i*.*e*. the mutational variance in units of residual variance.

This allows us to write the total genetic variance-covariance matrix as:

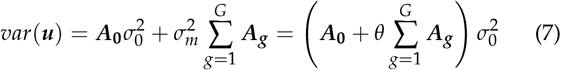

The kinship matrices, ***A***_**0**_ and ***A***_***g***_ were computed using **makeA** function from R package **pedigree** (Coster 2013). The model parameters 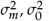 and *σ*were estimated using R package **sommer** (Covarrubias-Pazaran 2016, 2018).

#### Saclay DSEs common garden experiments analysis

All 10 traits measured in 2018 and 2019 (DSECG) were analysed separately using the same BLUPM decomposition as for DSEYM. However, to account for GxE interactions, the variance-covariance for the random effect due to standing genetic variation ***Var*(*G***^**0**^) (respectively incoming mutational variation ***Var*(*G***^***m***^ )) was modeled as:

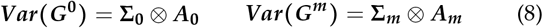

where ***A***_**0**_ (resp. ***A***_***m***_) is the kinship matrix defined previously and **Σ** is the covariance structure for progenitors among environments. We supposed an unstructured variance-covariance defined as:

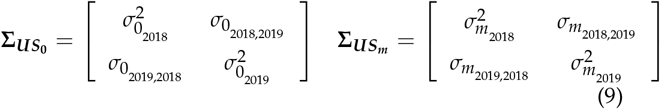

The unstructured model assumes *G×E* interactions and an environment-specific variance in addition to the resulting co-variances.

Furthermore, spatial variation was modeled by adding field coordinates (plots and columns) of each row as random effects. The model can be written as :

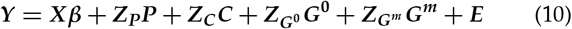

where :

- ***β*** includes *μ* the general mean, and a fixed year effects.
- ***P*** is a random plot effect per evaluation year such that ***P****∼* ***𝒩***_***p***_ **(0, Π**_***CS***_ ), where Π_*CS*_ is a compound symmetry structure, with plot effects nested within block.
- ***C*** is a random column effect per evaluation year such that ***C*** *∼* ***𝒩*** _***C***_ **(0, *X***_***DIAG***_ ) with:

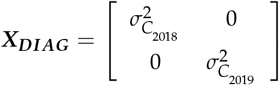
- ***G***^**0**^ is a random initial genetic effect per progenitor and evaluation year so that ***Var*(*G***^**0**^) = **Σ**_***US***_*⊗* ***A***_**0**_
- ***G***^***m***^ is a random mutational effect per progenitor and evaluation year so that ***Var*(*G***^***m***^ ) = **Σ**_***US***_ *⊗* ***A***_***m***_
- ***X***,***Z***_***P***_,***Z***_***C***_,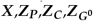 and 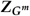 are the corresponding incidence ma-trices
- ***E*** the residuals so that ***E*** *∼* ***𝒩*** _***p***_ **(0, *E***_***DIAG***_ ) with :

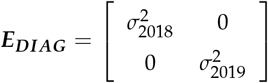

The environment specific BLUPs 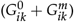 adjusted for co-variances were used in the subsequent analysis.

The validity of this model (Eq. 10) was tested first by computing the distributions of residual values which allowed to verify homogeneity of variances and dispersion around 0, and second using a likelihood ratio test to compare it with a model without *G × E* interactions where : 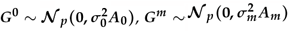 and ***E*** *∼* ***𝒩*** _***p***_ **(0, *σ***^**2**^)

##### Selection response analysis

We estimated the mean annual genetic gain for each trait as the slope of the linear regression between the predicted environment specific genetic values and generations. Each trait, evaluation condition, and family were analysed independently.

##### Distribution of mutational effects

To approximate the DFE of incoming mutations, we used the individual-predicted breeding values due to incoming mutations *u*_*m*_ in Eq. 2 and *G*^*m*^ Eq. 10 that we centered around the mean value of each corresponding generation for each family independently. This procedure allowed us to remove the effect of selection response. To approximate the DFE of selected mutations, we computed the distribution of mutational effects from mutational breeding values by conditioning on whether one of the progenies of the focal progenitor were selected or not in the next generation. To increase analysis power, we pooled the distributions across families of the same population (early or late flowering populations) accounting for the direction of selection (negative sign for unfavorable mutational effect and positive sign for advantageous ones). Hence we obtained a distribution based on 20 estimates per generation (progenitors) per population.

#### Genotyping data

We generated genotypic information for all progenitors along the pedigrees at potentially selected site and *de novo* mutations. More specifically, for a subset of polymorphic SNPs between Late and Early population at G13 (46 SNPs in MBS and 480 in F252), we produced partial genotyping data for 187 over 366 F252 progenitors and 190 over 354 MBS progenitors. We used pedigrees to infer missing genotypes at these loci.

##### SNP detection

We used an RNAseq dataset from five Early and Late progenitors of generation G13 of Saclay’s DSEs, originally produced to study the genes differentially expressed in the apical meristem during floral transition by Tenaillon *et al*. (2019). The use of an RNAseq dataset conditioned the detection of SNPs in genes expressed in the apical meristem, representing about 55% of all annotated maize genes.

Briefly, bulk of S2 seeds from each early/late of the five progenitors (G13) were sown in the field in 2012 and 2013. Total RNAs were extracted from 25 pools of meristems (each constituted by 16 to 31 meristems from the same progenitor and developmental stage) collected during the two years. The 25 corresponding RNAseq libraries were sequenced using 51 bp Illumina single reads. Because, we aimed at detecting SNPs between Early and Late genotypes within each DSE (F252 and MBS), we pooled RNAseq data from all libraries corresponding to the same progenitor, albeit different developmental stages, in a unique fastq file. More specifically, we pooled 6, 7, 4, 5, 3 libraries for each unique G13 progenitors in FE1, FL2.1, FVL, ME2 and ML1 families, called FEE, FLL, FVL, MEE, MLL, respectively in Tenaillon *et al*. (2019) (Fig. S2 and Fig. S3). Reads were trimmed and filtered for low quality and rRNAs were filtered out, following the procedure described in Tenaillon *et al*. (2019). Total number of reads per progenitor ranged between 59 399 604 and 96 708 953 (Table S1).

We further performed variant discovery within each inbred line independently. To do so, we followed the best practices of the Genome Analysis ToolKit (GATK, Auwera *et al*. (2013)) considering each of the five genotypes independently. We first mapped all reads on the B73 maize reference genome V4 (Jiao *et al*. 2017) using STAR (2.6.1) aligner (Dobin *et al*. 2013). We used the two-pass mode, in which a first mapping step detected spliced junctions subsequently employed as additional information to map the reads. We used the default parameters, except for the maximum number of duplicated match a read is allowed, that we set equal to one, hence keeping reads that strictly mapped to one locus. As for the post-mapping step, we used the Picard set of tools (3.8) to (1) index the reference genome, add read group information and sort the reads; (2) retrieve duplicates using MarkDuplicates; process reads incorrectly-mapped to intronic regions with SplitNCigarReads before using ReassignOneMappingQuality; recalibrate the quality scores thanks to baseRecalibrator, using the known variants (Chia *et al*. 2012) (downloaded from Ensembl database (ensembl;version=90;url=http://e90.ensembl.org/Zea_mays.)). We further called SNPs using the GATK’s tool HaplotypeCaller with a minimum PHRED score of 30.0. Next, we ran variant filtering steps with the VariantFiltration tool. Following GATK guidelines for RNA-seq data: we discarded clusters of at least 3 SNPs that were within a window of 35 bases by adding -window 35 -cluster 3 to the command; we filtered on Fisher Strand values (*FS* > 30.0) and Quality By Depth values (*QD* > 2.0). SNPs falling into non-assembled remaining scaffolds, chloroplastic or mitochondrial DNAs were eliminated. Because we expected very few heterozygous SNPs after 13 generation of selfing, we discarded them. Finally, we applied stringent filters for raw QUAL scores (*QUAL* > 40) and genotype quality (*GQ* > 20) and required that at least 5 reads covered a given SNP (*DP* > 5). Altogether, the variant discovery pipeline retained 46 SNPs in the MBS line and 7,030 SNPs in the F252 line. It is important to keep in mind that newly arising mutations appear in a heterozygous state. Therefore, by filtering out heterozygous mutations, we missed the opportunity to discover these mutations at their point of origin. Instead, we identified the subset of mutations that were homozygous for distinct alleles among any of the three progenitors FEE/FLL/FVL or between MEE/MLL.

##### SNP subsampling for KASPar™ genotyping

All 46 SNPs detected between MEE and MLL were retained and used subsequently to genotype MBS progenitors. Overall, most SNPs detected for MBS were located on chromosome 10 in 5 annotated genes (Table S2 and Table S3).

For F252, we subsampled 480 SNPs among the 7,030 detected between FVL, FLL and FEE. To do so, we defined several categories to maximize genome coverage of potentially selected site, while trying to identify *de novo* mutations. First, we decided to preferentially retain SNPs that fell within genes possibly involved in the genetic determinism of flowering time. We considered known flowering time genes in maize (**F_Candidate** in Tenaillon *et al*. (2019)), genes detected as differently expressed between Early and Late G13 progenitors in the RNA-Seq experiment (**Sel** genes in Tenaillon *et al*. (2019)) and genes associated to flowering time variation in a panel of 4,471 inbred lines/landraces ((Romero Navarro *et al*. 2017), **GWA_Candidate** in Tenaillon *et al*. (2019), Fig. S4). Because the DSEs are conducted under complete selfing and effective recombination is expected to be low, SNPs tended to form clusters. We defined these clusters without prior information using a k-means approach on the Euclidean distances between SNP positions (Fig. S5 and Fig. S6). Table S4 shows that the distribution of SNPs along the chromosomes was heterogeneous with only 4 SNPs in chromosomes 4 and 7, up to 131 in chromosome 6. Because we were interested in finding *de novo* mutations, we retained all the SNPs found isolated in 20kb regions, thereafter called **ALONE** (Fig. S4). As for clusters of SNPs, we retrieved one SNP every 200bp within candidate genes, and one SNP every 5 SNPs among SNPs separated by at least 200bp in cluster without candidate genes. Selected SNPs retained for F252 fell within 364 genes, 318 of them being represented by a single SNP, and 46 being represented by two to four SNPs. 48 SNPs were located in genes not annotated in B73 AGPv4 (Jiao *et al*. 2017) but present in the RNAseq data.

#### Inferring genotypes from partial KASPar™ genotyping data and pedigree relationships

In addition to the 308 progenitors measured in common garden experiments, all progenitors along the ancestral path of generation 13 (generation at which SNPs were detected) were genotyped so that overall, 187 F252 (+3 duplicated genotypes from different seed lots, *i*.*e*. genotyping replicates) and 190 MBS independent progenitors were effectively genotyped (Fig. S2 and Fig. S3).

DNAs were extracted from bulk of 15 plants (one standardized fresh leaf disk per plant, to ensure equimolar bulk). Extractions were performed from 30 mg of lyophilized adult leaf material following recommendations of DNeasy 96 Plant Kit manufacturer (QIAGEN, Valencia, CA, USA). We verified through simulations of our multiplication scheme, that the probability of not detecting one of the two alleles of an initial heterozygous progenitor was below 5% for a bulk of 15 S1, S2 or S3 plants.

The genotyping of 190 MBS progenitors for 46 SNPs and 190 F252 progenitors for 480 SNPs was conducting using KAS-Par™ technology (KBioscience’s competitive allele-specific PCR amplification of target sequences and endpoint fluorescence genotyping) on the GENTYANE technical center from INRAE (http://gentyane.clermont.inra.fr/contenu/presentation) The genotyping process was conducted using IFC Dynamic Arrays 96*96 / IFC Controller HX / Juno / Biomark™ HD Reader. The genotype attribution was conducted semi-manually, following a blind protocol, using Biomark software from the Fluidigm® company. Then each data point was validated manually by two independent operators using pedigree information to check for non-Mendelian segregation.

We produced a sparse genotyping matrix using KASPar™ genotyping, so that genotypic information was available only for a subset of all progenitors. From this subset we aimed at inferring all genotypes of the genealogy until generation 18^*th*^. In order to do so, we developed a likelihood model (Text S1), and used a parsimony algorithm developed by Durand *et al*. (2015) to infer missing data in the matrix of genotypes with the highest likelihood at every given SNP. In sum, for MBS (resp. F252), we produced an inferred genotyping matrix composed of 342 progenitors and 41 SNPs (resp. 351 progenitors and 456 SNPs). SNPs were oriented so that we followed in the subsequent analyses the fate of the minor allele.

#### Association mapping

Following Durand *et al*. (2015), we computed the additive (*a*) effects associated with each SNP, and for each trait independently, through linear regression:

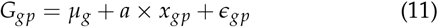

where *G*_*gp*_ is the environment specific (DSEYM or DSECG 2018 or DSECG 2019) predicted breeding value of progenitor *p* at generation *g. μ*_*g*_ is the average trait value calculated over all progenitors at generation *g*, and *x*_*gp*_ is an indicator variable equal to -1 for homozygous progenitor *aa*, 0 for heterozygous progenitor an 1 for homozygous progenitor *AA*. Allelic status *A* corresponded to the less frequent allele in the starting population at *G*_1_. When *G*1 allele frequency was equal to 0.5, we arbitrarily chose A to be the reference allele of B73 AGPv4 genome (Jiao *et al*. 2017).

We tested the significance of the association between trait variation and the segregation of alleles by simulating a null distribution for the additive estimated effect *a* using gene dropping simulations and the same model as Eq. 11. For standing variation, genotypes were simulated by dropping the two possible alleles throughout the pedigrees considering heterozygous ancestors at generation G0. At each generation, the genotype of each progenitor was randomly drawn from the alleles produced by its ancestor assuming Mendelian inheritance (Durand *et al*. 2015). For *de novo* mutations, we proceeded similarly except that simulations started at the genealogical position of the mutated individual. Note that for both standing and *de novo* variants, the initial detection of SNPs from RNASeq data generated at generation 13 introduced an ascertainment bias, *e*.*g*. their detection was conditioned on their differential fixation between early and late populations at G13. Therefore, we only used the subset of simulations that displayed the same genotypes in the 2 (resp. 3) individuals used for SNP detection in MBS (resp. F252). Interestingly, because of the small population sizes in our design (Desbiez-Piat *et al*. 2021), we expected standing alleles to be fixed at *G*_18_ within each family through drift (even without selection). This leads for each inbred line to different possible genealogical structures leading to a wide multi-modal distribution of possible additive values obtained through simulations (Fig. S7, Fig. S8 and Fig. S9). Hence, by conditioning on the observation of the genotype at generation *G*_18_, we were able to accurately control for genetic structure and deconvolute this mixture of distributions. Overall, probabilities of observed *a* additive values among 5,000 simulations per case were determined (p-values) through a symmetrical test. In addition, we controlled for multiple tests by setting a 5% False Discovery Rate (FDR).

Furthermore, the set of simulated allele trajectories through gene dropping was also used as the null hypothesis of fixation time against which observed trajectories were tested.

#### Evolution of residual heterozygosity across time points

In order to quantify the initial amount of residual heterozygosity and to verify that no pollen contamination had occurred during the course of the DSEs, we used previously published sequencing data at generation 13 for both F252 and MBS (Tenaillon *et al*. 2019) that we combined with whole genome sequencing data from generation 1 for both MBS and F252. Additionally, we analyzed unpublished RNASeq data produced at generation 18 for MBS. For F252 and MBS, generation 1 progenitors from which offspring were derived by selfing (see below) for sequencing are indicated Fig.S2 and Fig.S3.

##### G1 Plant material and DNA sequencing

Plants used for whole genome sequencing from F252 and MBS seedlots were obtained from three generations of selfing of one individual: FE1 G1 for F252 and ME2 G1 for MBS (Fig.S2 and Fig.S3). Plants were grown in standard conditions and transferred to dark chamber three days before seedling leaf sampling. Sampling was performed as follows: a leaf punch from each individual plant was sampled and used for genotype check using a set of 30 microsatellites spanning the 10 maize chromosomes. In parallel, 1 g of bulk leaf tissue from 4 or 5 plants was flash frozen and stored at -80° and subsequently used for high molecular weight extractions after microsatellite validation. Extracted DNAs were checked for purity and low fragmentation and subsequently pooled together to reach a pool of 15 (F252) and 12 (MBS) plants. Paired-end 150b reads were generated from 700-800bp insert size using PCR-free Illumina sequencing for a total of 255,411,879 reads for MBS and 251,968,819 reads for F252 (Table S1).

##### G1 mapping and variant detection steps

The resulting reads were assessed for quality using FastQC (Andrews *et al*. 2012), library bar-code adapters were removed, and reads were trimmed according to a quality threshold using TRIMMOMATIC (Bolger *et al*. 2014) invoking the following options (ILLUMINACLIP:adapters.fa-:2:30:10 LEADING:3 TRAILING:3 SLIDINGWINDOW:4:20 MINLEN:75). These filtered reads were used for downstream analysis. We aligned them to the B73V4 reference genome assembly (Jiao *et al*. 2017) using bwa v0.7.17 (Li and Durbin 2009). The resulting alignment files were sorted using SAMtools version 0.1.11 (Li *et al*. 2009) and were cleaned by keeping only primary alignment, properly-paired and unique. Duplicates sequences were removed using Picard Toolkit v2.26.11. SNP variants were called using the Genome Analysis Toolkit (GATK) haplotype caller (Bathke and Lühken 2021) following best practices (https://gatk.broadinstitute.org/hc/enus/sections/360007226651-Best-Practices-Workflows), filtered for mapping quality (MAPQ > 20) and read depth (n > 15). The combined data was again filtered to obtain biallelic sites across the two resequenced genomes using GATK GenotypeGVCFs option.

#### G18 Plant material, RNA sequencing and variant discovery

Selected progenitors from G18 (Fig.S2) were selfed to produce seeds that were sown at Université Paris-Saclay (Gif-sur-Yvette, France) during summer 2017 and 2018. We collected 22 and 26 plants for ME2 and ML1 progenitors respectively for a total of 48 plants (Table S1) and, dissected their shoot apical meristems that were flash frozen and stored at -80°. Total RNAs from individual meristems were extracted with the RNAqueous-Micro Kit (ThermoFisher) following the manufacturer instructions. Extracted RNAs were checked for purity and quantity using the Agilent 2100 bioanalyzer (California, USA). Libraries construction (SMART-Seq v4 Ultralow input, Takara) with barcoded adaptors as well as RNA sequencing on NextSeq500-Illumina (single-end 1×5 bp) was conducted by the POPS transcriptomics platform (IPS2, Université Paris-Saclay). Approximately 34 million reads were produced for each of the 48 samples. In order to perform detection of variants, we combined all reads obtained for a given progenitor: ME2 G18 (resp. ML1 G18) consisted in 22 (resp. 26) RNAseq runs (Table S1). Variant discovery from G18 RNAseq read libraries in MBS background (for ME2 G18 and ML1 G18) followed the exact same pipeline as for G13 progenitors.

#### Heterozygosity levels estimation

From RNAseq generated for the seven progenitors of G13 and G18, we filtered out genomic regions with a depth below 10 reads. We next computed within each genetic background, the subset of genomic regions covered in all progenitors of G13 and G18. We thereby obtained a partial representation of the progenitors genomes covering 32Mb for MBS and 34Mb for F252, distributed along contiguous genomic regions varying from 200bp to 4.5kbp spaced by roughly 900bp on average, and up to a maximum of 35Mb (Fig. S10 and Fig. S11). From this subset of regions, we selected those covered in the corresponding progenitor G1 DNAseq and encompassing SNPs in at least one of the progenitors. We determined observed heterozygosity levels within one progenitor by computing the average number of heterozygous sites weighted by the length of the considered regions. Note that differences of heterozygosity levels among individual was computed on a restricted set of high quality SNPs (QUAL *≥* 40, QG *≥* 20, DP *≥* 10).

## Results

### Two successive adaptive phases drive the selection response in Saclay’s DSEs

From Saclay’s DSE Yearly Measurement, the mean initial flowering time at *G*_0_ for F252, was equal to 66.4 DTF, emphasizing an earlier flowering genetic background than MBS which initially flowered around 79.5 DTF (Fig. 1). A simple linear regression on total predicted breeding values (Eq. 2) over generations summarizes the raw responses to selection. Within the two genetic backgrounds, the strongest selection response was observed in late populations with 0.346 DTF per generation in ML2 and 0.332 in FL2 (Table 1). The response for FVL family was so strong (1.016), that G14 plants were not able to produce enough viable offspring for the next generation (Durand *et al*. 2012), and that FL2 family had to be split into two (FL2.1 and FL2.2) families to recover a second biological replicates. Interestingly, the selection response was asymmetrical, so that early families responded less than late families with genetic gains comprised between -0.138 and -0.243 DTF per generation in accordance with (Durand *et al*. 2015). In addition, the response to selection appeared also less variable across early than across late families in F252 and MBS (Table 1 and Fig. 1).

**Table 1.** Selection response computed from total breeding values.

| Family | Mean genetic gain (in DTF eq. 20°C) | Std. Error for genetic gain | R<sup>2</sup> | G18 Cumulative Response |
| --- | --- | --- | --- | --- |
| FE1 | -0.138 | 0.008 | 0.220 | -4.10 |
| FE2 | -0.180 | 0.008 | 0.391 | -7.34 |
| FVL | 1.016 | 0.030 | 0.610 | 12.13 |
| FL2 | 0.332 | 0.045 | 0.179 | 3.16 |
| FL2.1 | 0.064 | 0.017 | 0.024 | 4.57 |
| FL2.2 | 0.079 | 0.017 | 0.062 | 4.26 |
| ME1 | -0.243 | 0.006 | 0.601 | -5.96 |
| ME2 | -0.197 | 0.007 | 0.443 | -4.95 |
| ML1 | 0.170 | 0.012 | 0.211 | 8.63 |
| ML2 | 0.346 | 0.016 | 0.364 | 15.53 |

**Figure 1.**
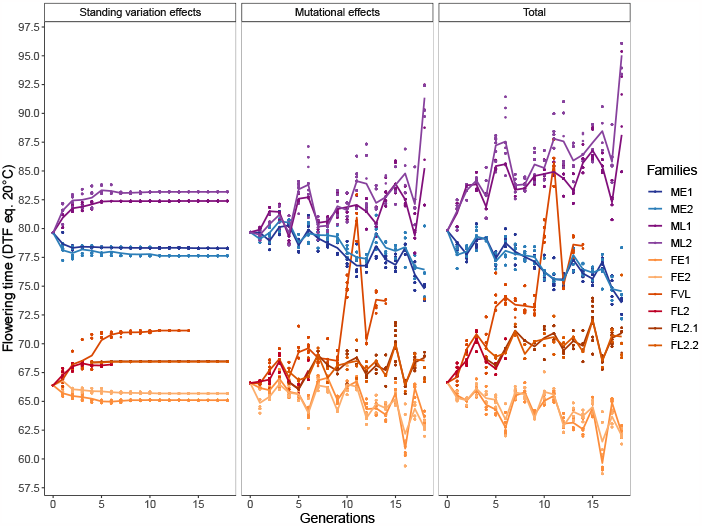
Saclay’s DSEYM Decomposition of the breeding values (right panel) into an initial standing variation component (left panel) and a mutational effect component (middle panel), and their evolution through generations. The right panel indicates the sum of the two predicted values in F252 (orange-red) and MBS (blue-purple). Colors indicate the families, and lines indicate the evolution of the mean value per generation and family through time. Each dot corresponds to a progenitor.

Several factors could contribute to the robust and sustained response to selection. These factors include *de novo* mutations, recalcitrant heterozygosity lasting longer than predicted (3-5 generations) by Desbiez-Piat *et al*. (2021), and involuntary gene flow through pollen contamination. To ensure the reliability of our findings, we took steps to eliminate contamination as a potential confounding factor. To do this, we conducted a thorough analysis of the raw KASPar genotyping data, searching for any abrupt large heterozygous regions that might suggest contamination. We found no evidence of such regions infiltrating the pedigrees, confirming that contamination was not a concern (Fig. S12, Fig. S13). Next, we monitored the level of residual heterozygosity in the progenitors at generation 1 and 13 for F252 and at generation 1, 13, and 18 for MBS. Our observations revealed that there was no global increase in heterozygosity between generation 1 and 13 (and 18 for MBS), further supporting the absence of pollen contamination (Table 2, Fig. S10, Fig. S11).

**Table 2.** Distribution of heterozygosity levels at generation 1, 13 in MBS and F252 and generation 18 in MBS.

| Progenitor | Average (SD) | 90% Quantile | 99% Quantile |
| --- | --- | --- | --- |
| ME2 G1 | 0.1070%<br>(0.679%) | 0% | 3.60% |
| ME2 G13 | 0.0518%<br>(0.329%) | 0% | 1.33% |
| ML1 G13 | 0.0405%<br>(0.305%) | 0% | 1.20% |
| ME2 G18 | 0.0499%<br>(0.378%) | 0% | 1.54% |
| ML1 G18 | 0.0503%<br>(0.380%) | 0% | 1.60% |
| FE1 G1 | 0.1130%<br>(0.697%) | 0% | 4.00% |
| FE1 G13 | 0.0466%<br>(0.306%) | 0% | 1.33% |
| FL2.1 G13 | 0.0457%<br>(0.305%) | 0% | 1.33% |
| FVL G13 | 0.0432%<br>(0.297%) | 0% | 1.33% |

In the initial sequencing of generation 1 progenitors for both MBS and F252, we observed an average residual heterozygosity of 0.1%, indicating a small amount of initial standing variation after just one generation of selfing and selection (Tab 2). Inspection of genomic patterns of residual heterozygosity revealed that the majority of regions (94%) had no heterozygous sites (Tab 2). Interestingly, as we progressed to subsequent generations, we found that the average residual heterozygosity decreased by half, reaching approximately *≈*0.05% at generations 13 and 18 where 99% of regions encompassed less than 1.60% heterozygous sites (Tab 2). The decline of heterozygosity observed in later generations highlighted the persistence of recalcitrant heterozygosity. Noteworthy, we observed pattern similarity between generations, between families but also between genetic backgrounds (Fig. S10, Fig. S11) suggesting shared mechanisms underlying recalcitrant heterozygosity.

Note that simulations of the experimental scheme predicted a mean level of heterozygosity of 0.3 % (SD=0.3%) at generation 1 considering the conventional inbred line production scheme and, a mean level of 0.083% (SD=0.047%) at generation 20 (Desbiez-Piat *et al*. 2021). However, of the high quality heterozygous sites in ME2 G1: 51% remained heterozygous in ME2 G13 and ME2 G18, indicating that they did not contribute to the selection response, 33.4% were fixed between G1 and G13 and, 6.4% were fixed between G13 and G18. Note that 9.2% were heterozygous in ME2 G1 and ME2 G18 but not in ME2 G13, because ME2 G13 was not on the ancestral path of ME2 G18. We revealed a similar pattern in the late F252 family: of the initial heterozygous sites in ME2 G1: 49.1% were still heterozygous in ML1 G13 and ML1 G18, 34.1% were fixed between G1 and G13, and 6.6% were fixed between G13 and G18. 10.2% were heterozygous in ME2 G1 and ML1 G18 but not ML G13. Likewise in F252, of the high quality heterozygous sites in FE1 G1: 55.3% were still heterozygous in FE1 G13 and the rest was fixed between G1 and G13; for FE1 G1: 51.3% were still heterozygous in FL2.1 G13 and FVL G13, 42.8% were heterozgous in FE1 G1 but neither in FVL G13 nor FL2.1 G13, 3.5% being specific to G1 and FVL G13 and 2.4% being specific to G1 and FL2.1 G13. Altogether these observations revealed the existence of two distinct phases that played crucial roles in maintaining the selection response. In the initial phase, residual heterozygosity exerted a strong influence, significantly contributing to the selection response. However, as we progress to subsequent generations, the role of residual heterozygosity diminished gradually. Furthermore, an heterogeneous distribution of heterozygosity was observed along the genomes (Fig. S10, Fig. S11).

To investigate how much of the phenotypic response of Saclay DSEs observed from yearly flowering time measurements could be explained by initial standing variation *versus* incoming mutational variance, we used Wray (1990) BLUPM model. We estimated an initial standing variance of 3.15 (resp. 3.72) in MBS (resp. in F252), and mutational variance estimate of 0.357 (resp. 0.203) (Table 3). These translated into mutational heritabilities of 0.116 (resp. 0.152) in MBS (resp. in F252) (Table 3). The relative importance of mutational variance in the response to selection was higher in MBS with an estimate of *θ* of 0.113 than in F252 (*θ* = 0.0547, Table 3), consistent with the fixation in F252 of a strong effect standing variant (Durand *et al*. 2012).

**Table 3.** Variance decomposition of the selection response for flowering time from yearly measurements. Variance components (Standard Error)[Zratio] were computed by contrasting early and late families. 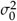corresponds to initial standing variance, 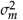 to mutational variance and *σ*^2^ to the residual variance.

|  | MBS | F252 |
| --- | --- | --- |
| $\sigma_0^2$ | 3.15 (1.34)[2.35] | 3.72 (0.189)[19.7] |
| $\sigma_m^2$ | 0.357 (2.72E-02)[13.1] | 0.203 (1.89E-02)[10.7] |
| $\sigma^2$ | 3.06 (6.62E-02)[46.3] | 1.34 (5.85E-02)[22.9] |
| $\theta = \frac{\sigma_m^2}{\sigma_0^2}$ | 0.113 (4.97E-02)[2.28] | 0.0547 (5.93E-03)[9.23] |
| $h_m^2 = \frac{\sigma_m^2}{\sigma^2}$ | 0.116 (9.28E-03)[12.5] | 0.152 (1.57E-02)[9.67] |

More detailed observations of the evolution of standing and mutational variance through generations of selection confirmed these trends (Fig. 1). In MBS, after 18 generations of selection the average total genetic value predicted by the model reached 8.6 for ML1 but 15.5 DTF for ML2 (Fig. 1, right panel). In F252, the very late family FVL quickly responded to selection which led to a drastic shift around 12 DTF and the two derived late families FL2.1 and FL2.2 from FL2, were only shifted by 4 DTF (Fig. 1). The asymmetrical response to selection for early families was observed within each component of the selection, so that for example, ME1 (resp. ME2) total gain of -6.0 (resp. -5.0) could be decomposed into a standing variation effects of -1.2 (resp. -1.9) and a mutational effect of -4.8 (resp. -3.1), these cumulative effect being smaller in absolute values than their late counterparts. For F252, a shift around -1.3 (resp. -0.7) DTF was also reached due to standing variation in FE1 (resp. FE2), but mutations accounted only for -2.8 (resp. -3.7) DTF, so that they plateaued to a maximum shift around -4 DTF after 10 generations of selection. Finally, the decomposition of the selection response into a shift of 2.9 (resp. 3.7) DTF (left panel) for ML1 (resp. ML2) due to standing variation and a shift of 5.7 and 11.9 DTF respectively due to incoming mutational implied that mutations accounted for 67% and 77% for ML1 (resp. ML2) of the total genetic gain. On the other hand for FL2.1 and FL2.2 mutations accounted for 54% and 52% of the total genetic gain respectively (Fig. 1), which confirmed the prominent role of mutational input to the selection response in MBS.

### Selected incoming mutations are biased towards favorable effects in the direction of selection

We approximated the distribution of the mutational effects of incoming mutations and that of the selected mutations. In the following, we designated mutations conferring earliness in the early populations and lateness in the late populations as favourable. Compared with non selected incoming mutational effects that displayed an average effect of *−*0.439 DTF for MBS and *−*0.352 DTF for F252, we observed among the selected mutations, a strong enrichment in favorable mutations with an average mutational effect of +0.299 DTF and +0.278 DTF for MBS and F252, respectively (Table 4 and Fig. 2). The top 5% selected mutational effects were greater than +2.09 DTF and +1.36 for MBS and F252 respectively, while the 95 percentile of unselected mutational effects equaled +1.1 and +0.683 for MBS and F252. However, we selected up to 25% of unfavorable mutations in MBS (resp. 18% in F252) (*i*.*e*. with sign opposite to the direction of selection) so that for example 5% of the selected mutations had effects below -1.01 DTF in MBS and -0.739 DTF in F252 (Table 4 and Fig. 2).

**Table 4.** Mutational effect percentiles (Q) according to evaluation environments for selected and unselected mutations (in DTF eq. 20^*o*^ *C*)

| Inbred | Selected | Env. | Q5 | Q50 | Mean | Q95 |
| --- | --- | --- | --- | --- | --- | --- |
| MBS | No | DSEYM | -1.94 | -0.412 | -0.439 | 1.1 |
| MBS | No | 2018 | -1.42 | 0 | -0.12 | 0.693 |
| MBS | No | 2019 | -1.58 | 0 | -0.128 | 0.826 |
| MBS | Yes | DSEYM | -1.01 | 0.173 | 0.299 | 2.09 |
| MBS | Yes | 2018 | -0.982 | -0.00239 | 0.0818 | 1.47 |
| MBS | Yes | 2019 | -1.07 | 0 | 0.0875 | 1.64 |
| F252 | No | DSEYM | -1.66 | -0.335 | -0.352 | 0.683 |
| F252 | No | 2018 | -0.882 | -0.000976 | -0.106 | 0.507 |
| F252 | No | 2019 | -1.17 | 0 | -0.14 | 0.721 |
| F252 | Yes | DSEYM | -0.739 | 0.28 | 0.278 | 1.36 |
| F252 | Yes | 2018 | -0.857 | 0.0292 | 0.0837 | 1.02 |
| F252 | Yes | 2019 | -1.07 | 0.0175 | 0.11 | 1.3 |

**Figure 2.**
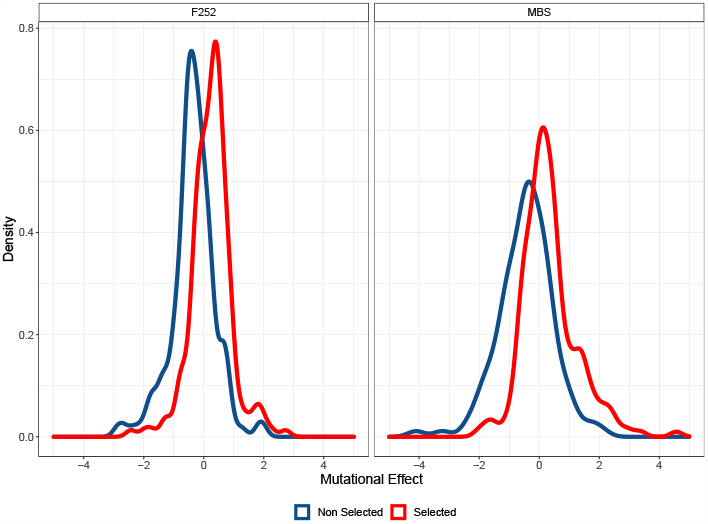
Distribution of effects of non selected (blue distribution) and selected (red distribution) incoming mutation in F252 and MBS. Positive values correspond to mutational effects of the same sign as the direction of selection.

The individual monitoring of mutations along pedigrees and the evolution of their frequency in populations provide essential information on their nature (standing or incoming), their effects and the evolutionary forces presiding over their fate. We followed a subset of standing variants and *de novo* mutations by producing genotypic information at 41 (resp. 456) SNP for MBS (resp. F252) using KASPar genotyping technology. We inferred the genotypes of all individuals along the pedigrees. SNPs were detected from RNAseq data conditioning on their differential fixation between early and late populations at generation 13 (Tenaillon *et al*. 2019). Due to the limitations of RNAseq data, such as incomplete coverage of annotated genes and the emphasis on exonic SNPs, we might not have detected all standing and *de novo* mutations, especially those occurring in regulatory regions known to be functionally relevant. Nevertheless, despite these imperfections, we do not anticipate any bias in our estimation of the standing versus mutational variance because no linkage disequilibrium was expected between these two types of mutations.” Because of the high drift-high selection regime, differential fixation between populations can arise by either selection or drift. To enrich for potentially selected variants in our KASPar assays, we therefore devised a number of filters by targeting flowering time genes as well as flowering time genome wide association (GWA) hits.

We detected two *de novo* mutations per inbred line (Fig. S14). Both mutations detected in MBS appeared in the late flowering ML1 family. The first one appeared at generation 3, on chromosome 2 at position 6,294,005. This mutation was the unique one within a 20kb-window. It appeared in the gene Zm00001d002125, also known as RPD2 which encodes the nuclear RNA polymerase D2/E2, the RNA polymerase IV second largest subunit. RNA polymerase IV plays a role in siRNA-directed DNA methylation (RdDM) and silencing of genes and endogenous repetitive elements; and it has also been shown to regulate flowering-time genes (Pikaard *et al*. 2008). It fixed in 3 generations. The second appeared at generation 10, on chromosome 6 at position 148,046,062. This mutation also appeared to be isolated (i.e. the single one within 20 kb-window). It fell within the Zm00001d038104 gene encoding for a mannosylglycoprotein endo-beta-mannosidase. Interestingly, this gene has been shown to be differentially expressed between the early and the late F252 genotypes at generation 13 (Tenaillon *et al*. 2019). It fixed in 4 generations. The two mutations detected in F252 appeared respectively in early flowering FE1 family and late family FL2.1 (note that FL2, FL2.1 and FL2.2 are represented together Fig. S14). The first one appeared at generation 4, on chromosome 3 at position 141,724,144 and was the unique detected SNP within a 20kb window, but did not lie within any annotated gene. It fixed in 6 generations. The second mutation appeared around generation 5 on chromosome 6 at position 130,321,359 and was the single one detected SNP within a 20kb window. It occurred in gene Zm00001d037579, coding for a serine/threonine-protein phosphatase 4 regulatory subunit 3.

In Fig. 3, we observed that mutation 2 appeared as a homozygous variant in FL2.1 at generation 6, and it was not detected in any other individual before generation 3. Since there was only one progenitor from generation 4 to 6 (Figure S3), the inference algorithm lacked sufficient power to precisely determine the exact time of appearance. To handle this uncertainty, we considered the specific mutation to have emerged in the progenitor of FL2.1 at generation 5 for the null model.

**Figure 3.**
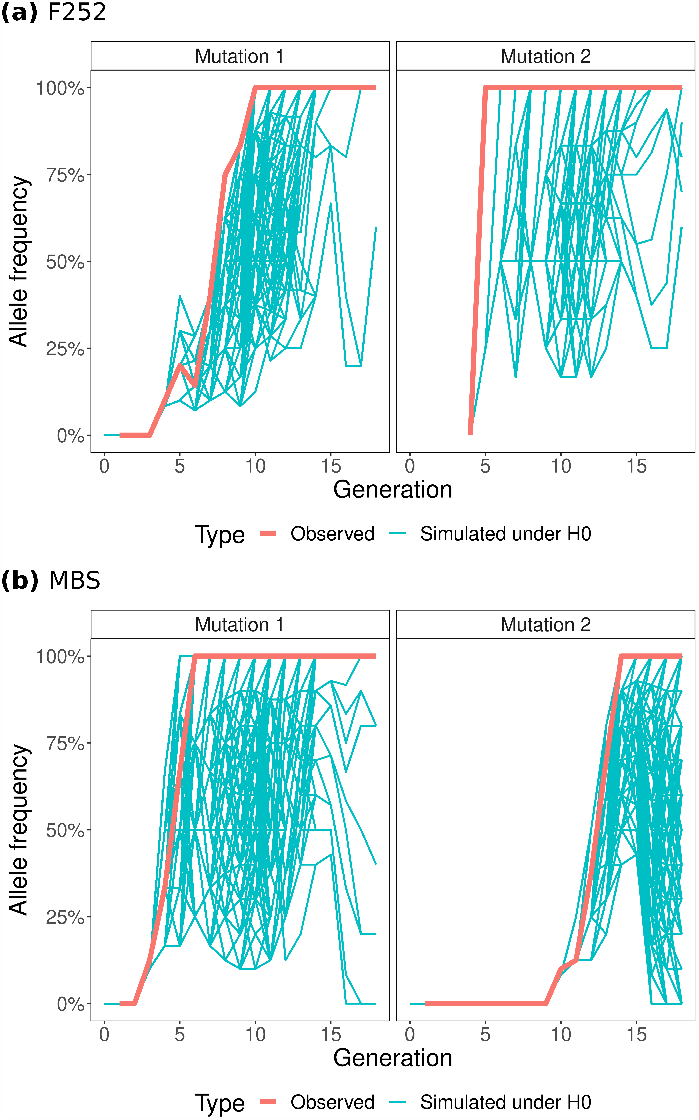
Allele frequency of *de novo* mutations through generations in F252 (a) and MBS (b). Red lines indicate the observed frequency changes while blue lines represent allele trajectories obtained from gene dropping simulations.

We devised a null model for fixation time that explicitly accounted for the generation of occurrence of the mutations (ancestor for standing SNPs and later generations for *de novo* mutations) and their segregation along pedigrees. We performed gene dropping simulations to ask whether their fixation times were consistent with drift (H0). Comparison between observed and simulated allele trajectories for the 4 *de novo* mutations revealed that polymorphic SNPs fixed quicker (*i*.*e*. 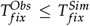) than expected by drift (Fig.3, Fig. S15). The observed fixation times (in 4,5,7 and 1 generation) were shorter than the mean fixation times under H0 (5.25,5.07, 7.2 and 3.3 generations, respectively, Fig.3) with correspondingly significant p-values (<0.05) for 3 out of 4 mutations (P-value=0.11 and <0.001 in all other cases, respectively). Deviations from H0 suggested that selection drove the observed patterns for 3 *de novo* mutations (MBS mutation 2 and the two F252 mutations, Fig.3). As for minor standing alleles, we observed no significant difference between the observed mean fixation time and the expected mean fixation time under H0 in MBS. In contrast, in F252 we observed a mean fixation time of 6.04 (SE:0.09) greater than the expected mean fixation time of 3.64 (SE:0.01) (Fig. S16).

Furthermore, the average number of generations for the minor standing alleles to be lost was smaller for observed SNPs with 6.55 (SE:0.19) generations compared with 8 (SE:0.02) for simulations under H0 in MBS. In F252, we measured an average number of generations for mutation loss of 5.71 (SE:0.11), also smaller than the 6.28 (SE:0.01) generations simulated under H0. Interestingly, in F252, 16% of the standing variants were lost quicker than 95% of the corresponding gene-dropping simulations, but 0.6% had a time-to-loss greater than 95% of the simulations. In MBS, the respective percentages were equal to 11% and 0% (Fig. S16). This suggests that selection might better eliminate minor unfavorable standing alleles, while being less efficient (longer than random fixation time) at fixing minor beneficial alleles.

### Common garden experiments imperfectly mirror past intensity of selection response for flowering time

Historically, Saclay DSEs yearly measurements (DSEYM) of flowering time were used to estimate breeding values for each progenitor. However, because the environment varies from year to year, these estimates are intrinsically limited in their ability to disentangle genetic effects from phenotypic plasticity -the capacity of a genotype to produce multiple phenotypes in response to environmental variations -from *G×E* -when two different genotypes respond to environmental variation in different ways. Plasticity and *G×E* are widely present (de Villemereuil *et al*. 2016) and could explain part of the observed selection response. To test this hypothesis, we conducted two common garden experiments in 2018 and 2019 (DSECG) aiming at evaluating the response to selection at 10 traits -including flowering time -across generations.

We used a likelihood ratio test to show that the model including GxE interactions (Eq 10) was significantly better than model without for all measured trait and the two genetic backgrounds (*p <* 10^*−*3^). Similarly, we validated that model Eq. 10 with an unstructured variance-covariance performed significantly better than other variance-covariance structures using a likelihood ratio test (*p <* 0.05). We visually validated for flowering time the omnipresence of *G×E* interactions by plotting reaction norms using environment specific genetic values per progenitor (Fig. S17). Interestingly, few changes in rank between individuals were observed between 2018 and 2019 but significant changes in scale were observed. To characterize the impact of evaluation environment on the estimation of the selection response in DSECG, we fitted for each family and evaluation year independently, a simple linear regression model on the environment specific genetic values (Table S5). On average, all genotypes of all families flowered around 7.7 DTF eq. 20^*o*^ *C* (resp. 11.7 DTF eq. 20^*o*^ *C*) later in 2019 than in 2018 (Fig. S17) for MBS (resp. F252), exemplifying flowering time plasticity for all genotypes.

Analysis of climatic records revealed that the two evaluation years were rather similar and characterized by an exceptionally hot and dry summer compared to selection years (Fig. S18). Hence, *G×E* interactions could be improperly captured in DSECG because of the lack of environmental contrast between the two evaluation years. We therefore compared the DSECG environment specific breeding values to the ones computed from DSEYM that captured variable climatic conditions over the course of selection years/generations as highlighted by the large ranges of values shown in Fig. S18. In contrast with DSECG, we observed frequent rank changes in flowering time across progenitors (Fig. S17). In addition, we pinpointed important differences in the response to selection between DSECG and DSEYM. For late families, the observed selection response was less pronounced in DSECG compared with DSEYM (as shown Fig. S19). Within MBS, Late flowering ML1 and ML2 presented selection response approximately equal to DSEYM analyses in both evaluation years (around 0.2 and 0.4 DTF eq. 20^*o*^ *C* per generation respectively, Table S5 and Table 1). For late F252 families in contrast, the observed selection responses were greater in 2018 and 2019 than in DSEYM for FVL (1.3 in 2018 and 1.3 in 2019 compared to 1.02 DTF eq. 20^*o*^ *C* per generation in DSEYM), smaller for FL2.1 (0.048 DTF eq. 20^*o*^ *C* per generation in 2018, 0.012 in 2019 and 0.064 in DSEYM) and were not significant for FL2.2 in DSECG (compared to a significant 0.072 DTF eq. 20^*o*^ *C* per generation genetic gain in DSEYM). For FL2 family, the ancestors of both FL2.1 and FL2.2, no significant response was observed in 2018 but a mean gain of 0.31 DTF eq. 20^*o*^ *C* was observed in 2019 and 0.332 DTF in DSEYM. For early families, the mean genetic gain for ME1 around was around -0.1 DTF eq. 20^*o*^ *C* per generation in 2018 and 2019 (-0.24 in DSEYM), it was not significant for ME2 in 2018 and 2019 (-0.20 in DSEYM), and around -0.06 in 2018 and 2019 for FE1 (-0.14 in DSEYM), -0.05 and -0.08 for FE2 (-0.18 in DSEYM). Overall, our results suggest a pattern of selection response consistent across evaluation years (2018, 2019) and, between DSEYM and DSECG, towards lateness, while the pattern towards earliness was less pronounced and more variable between DSEYM and DSECG.

Finally, we asked whether the previous decomposition of the selection response into initial genetic variance and mutational variance was conserved in DSECG (Table 3 *vs* Table S6). Interestingly, the qualitative contribution of initial standing variance and mutational variance to the selection response in DSECG (represented by the predicted genetic values due to mutational input (middle panel) or to initial variance (left panel) Fig. 4) indicates a diversity of behaviours. For example, in ML1 and ML2, the strong observed response was due to the sum of both initial variance and mutational input that increased the time to flowering in 2018 and in 2019. Conversely, the reduced selection response compared to DSEYM for ME2, FE1 and FE2 was due to a reduced contribution of the effect of the mutational input -less expressed -in 2018 and 2019 (Fig. 1 *vs* Fig. 4). This translated into a highly reduced difference between the distribution of selected *vs* unselected mutational effects. The mean selected effects was around 0.08 DTF in 2018 and 2019 compared to 0.3 DTF in DSEYM for MBS and F252, while the average unselected effects were equal to around -0.1 in 2018 and 2019 compared to-0.4 in DSEYM (Table 4 and Fig. S20). On the other hand, the strong response to selection for lateness in FVL was due to a magnified effect of its standing variance contribution (Fig. 4).

**Figure 4.**
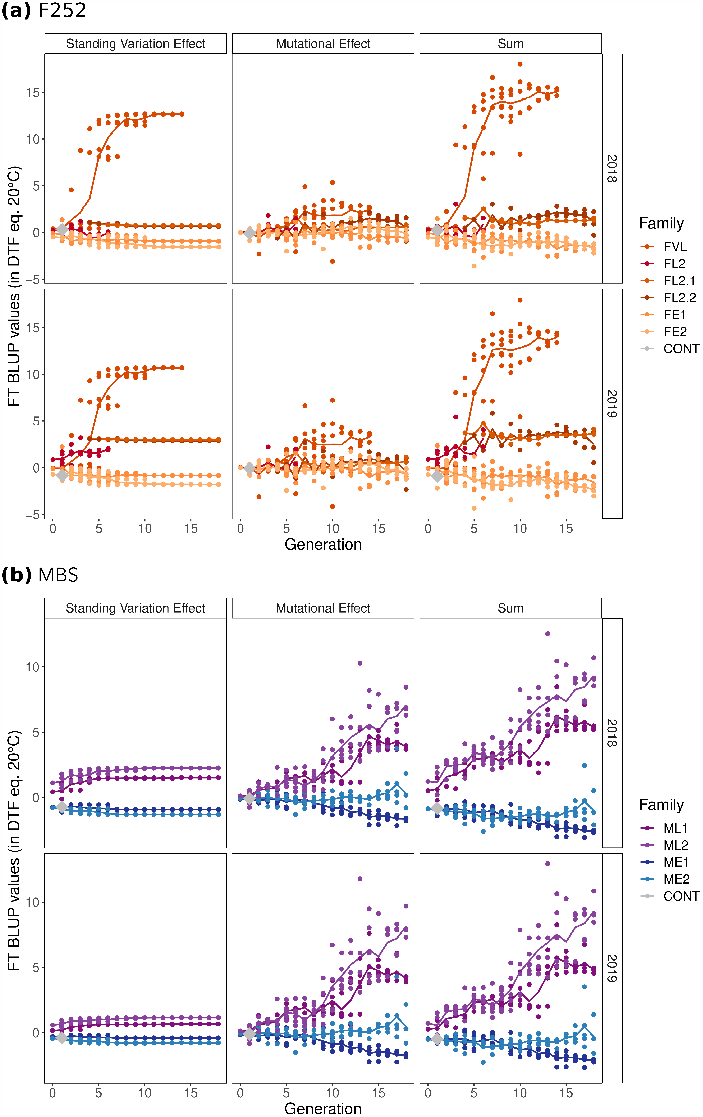
Saclay’s DSECG, Decomposition of the environment specific breeding values (right panels) into an initial standing variation component (left panels) and a mutational effect component (middle panels), and their evolution through generations in (a) F252 and (b) MBS. The right panels indicate the sum of the two predicted values. Colored lines indicate the evolution of the mean value per generation and family through time. Each dot corresponds to a progenitor.

### Selection for flowering time drives the exploration of the multivariate phenotypic and genotypic space

We asked how selection for flowering time had impacted adaptive trajectories in the phenotypic and underlying genotypic space defined at 9 other non-focal traits measured in DSECG. We applied the same model (Eq. 10) on all measured traits and genetic backgrounds independently. Because all genetic variances of the phyllochron were not significantly different from zero (Table S6) for F252 and for MBS (except 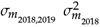) *i*.*e*., total leaf number could not be attributed to changes in leaf emission rate, we excluded this trait from subsequent analyses.

For the nine remaining traits including flowering time, we computed pairwise correlations between traits from breeding values (Fig. S21) and validated that trait correlations patterns were overall conserved across genetic backgrounds (F252 *vs* MBS). For example, our results highlighted a positive correlation between flowering time, leaf number and ear ranks in both genetic backgrounds, early and late populations and evaluation years. These correlation intensities were particularly pronounced in late families for which selection response for flowering time was the strongest (Fig. S21). Noticeably, while this group of trait was on average negatively correlated to yield components (TKW and EW) and to leaf length in late MBS and F252 populations, we observed null or positive correlations in early populations. Differences between early and late populations indicated that selection for flowering time in some instances impacted the structure of the genetic correlation matrix. Likewise, the evaluation environment (2018 and 2019) also impacted genetic correlations but in their intensities rather than in their signs (Fig. S21).

We next focused on the temporal patterns of correlative responses that is the exploration of the genotypic space over time. Families displayed distinct trajectories in the genotypic space under High-Drift High-Selection regime, yet repeatable between evaluation years (Fig. 5 and Fig. S22). This indicated that the correlations among traits were overall conserved. Slight differences in their magnitude across evaluation years (DSECG), however, resulted in translation patterns. Two mechanisms governed adaptive trajectories: first, the observed strong selection response for flowering time drove the evolution of correlated traits; second, the evolution of traits that were less correlated to flowering triggered a stochastic exploration of the genotypic space (Fig. 5 and Fig. S22). Genotypic evolution related to flowering time and correlated traits was captured by the first PCA axis that explained 61% of the total variation ((Fig. 5). This axis differentiated early from late flowering populations. In contrast, the second and third axes that captured 22% and 7% of the variation respectively, highlighted a stochastic space exploration of families within early and late population for traits such as leaf length, plant height, thousand kernel weight (TKW), and ear weight, especially in MBS. In MBS early families, a positive correlation was observed between flowering time and plant height (0.56) while a negative one was observed in MBS late families (-0.82) suggesting variation in internode length between populations (Fig. S23 and Table S5). Interestingly, a global trend of decreasing leaf length was observed in all F252 families (Fig. S24), and in early MBS families, such that ME2 leafs length had decreased by almost 7cm by generation 10 (Fig. S23, Table S5).

**Figure 5.**
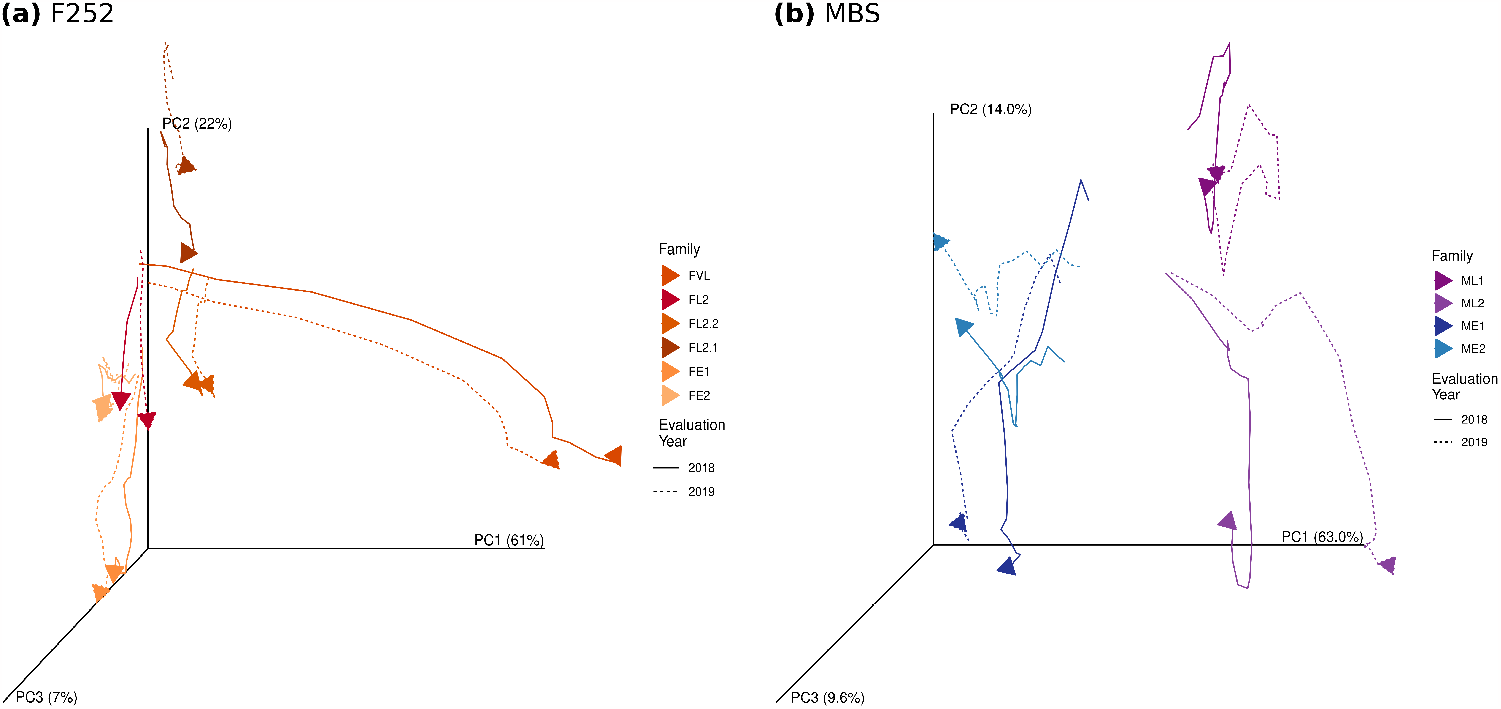
**Evolution through generations of the multitrait breeding values across families in 2018 and 2019** for F252 (a) and MBS (b). The genotypic space defined by 9 traits is summarized by a PCA on total BLUP values. Colored lines indicate the evolution through time of the rolling mean per generation, family and evaluation year on a 3-years window. Colors indicate the families. Solid lines refer to 2018 while dotted lines refer to 2019.

We asked more specifically what was the role of incoming mutations on the exploration of the genotypic space. All mutational variances and covariances for both genetic backgrounds were significantly different from 0 (Table S6) which suggested that incoming mutational variation sustained the exploration of allaxes of the genotypic space. We centered mutational effect within generation and family to remove selection response effect. Pearson’s correlation coefficients for all pair of traits highlighted the high degree of pleiotropy of incoming mutations (Fig. 6). In line with results computed from breeding values, the structure of the correlation matrix was conserved between genetic backgrounds, but not between late and early populations within genetic background. For example, plant height tended to be negatively correlated with total leaf number, ear ranks, brace roots in late MBS and F252 but positively correlated in both early MBS and F252 populations. This was also the case of leaf length, an other growth parameter. This suggested that incoming mutations interacted epistatically with the genetic background. Interestingly, while correlation matrices computed from incoming and standing variants were generally consistent (Fig. S25; Fig. 6), we pinpointed to some differences in sign and intensity. For example in late F252, leaf length and leaf number where negatively correlated when considering incoming mutations but positively correlated when considering standing mutations (Fig. S25, Fig. 6); and in late MBS flowering time was negatively correlated to plant height considering *de novo* mutations but positively correlated considering standing variation. This observation may explain abrupt changes in the direction of the exploration of the phenotypic space after 5-6 generations, for example for ML2 or FL2.2 in Fig. 5.

**Figure 6.**
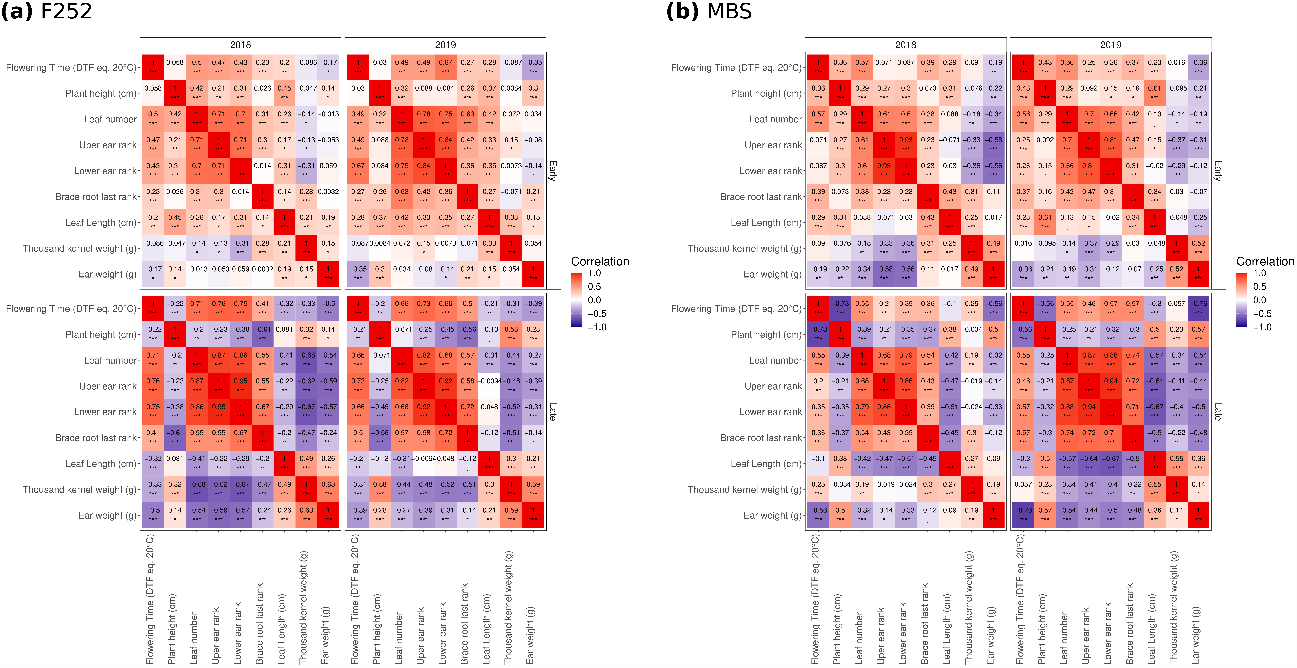
Pairwise correlation matrix between the predicted incoming mutational contribution for all measured traits in Early and Late populations in 2018 and 2019 for (a) F252 and (b) MBS. Pearson’s correlation coefficients were computed after centering values per generation to remove past selection effects. Color corresponds the intensity of the correlations. ‘***’, ‘**’,’*’,’.’ Indicates statistical significance at the 10^*−*3^, 10^*−*2^,5 *×* 10^*−*2^ level, respectively.

In order to provide an experimental validation of standing variant and *de novo* mutation effects as well as to explore their degree of pleiotropy, we conducted an association study on a restricted set of 456 and 41 SNPs for F252 and MBS respectively. Strikingly, 3 of the 4 detected *de novo* mutations were significantly associated with several traits, reinforcing the hypothesis of high degree of pleiotropy of selected *de novo* mutations. In-deed, in MBS, the two detected mutations on chromosome 2 and 6 were significantly associated with at least one of the traits (Table 5, Fig. S27). More specifically, the mutation on chromosome 2 was significantly associated to flowering time measured from DSEYM with an additive effect of 1.43 DTF Eq. 20^*o*^ *C* (Table 5), and to flowering time in 2019 with an effect of 0.73, but not significantly associated in 2018 despite a strong effect of 0.93 DTF Eq. 20^*o*^ *C*. This mutation was also significantly associated to leaf length in 2018 and 2019 with effects of 1.06 and 1.14 cm respectively. The other mutation detected on chromosome 6 in MBS was only associated to plant height in 2019, despite strong estimated effects of 0.90 from Saclay DSEYM, 0.63 in 2018 and 0.52 DTF Eq. 20^*o*^ *C* in 2019. Surprisingly, 37 out of 39 detected standing variants, that clustered together on chromosome 10, were associated to ear weight in 2018 and 2019 (Table S7). For F252, the two detected *de novo* mutations were significantly associated to flowering time in all evaluation environments (Table 5, Fig. S26) and the mutation located on chromosome 3 was significantly associated to 5 out of 8 measured traits in at least one evaluation environment, while the mutation located on chromosome 6 was significantly associated to all measured traits in at least one environment. This high degree of pleiotropy contrasted with what we found for standing variants: out of the 454 detected standing variants, 99 were significantly associated to at least one trait, and only 11.3% were associated to more than 2 traits (Table S8). In F252, 80 genes located in 27 different clusters of SNPs presented at least one SNP significantly associated to at least one trait. A single SNP (located in Zm00001d035439) displayed a significant association with flowering time measured from DSEYM, but 32 others — located in 23 different genes in 8 distinct clusters (Fig. S26) — were significantly associated to flowering time in DSECG.

**Table 5.** Significant trait association (allele effect) of four detected *de novo* mutations in 2018 and 2019. The first columns provide information on the chromosome, position, gene id and and known function.

| Inbred | CHROM | POS | Gene id | description | 2018 | 2019 | DSEYM |
| --- | --- | --- | --- | --- | --- | --- | --- |
| MBS | 2 | 6,294,005 | Zm00001d002125 | nuclear RNA polymerase D2/E2 | LL(1.06) | FT(0.73); LL(1.14) | FT(1.43) |
| MBS | 6 | 148,046,062 | Zm00001d038104 | Mannosylglycoprotein endo-beta-mannosidase | NA | HT(1.05) | NA |
| F252 | 3 | 141,724,144 | NA | NA | FT(-1.18); HT(-3.51); BR(-0.04); LL(-0.47) | FT(-1.27); HT(-2.13); #L(-0.28); BR(-0.07); LL(-0.88) | FT(-1.57) |
| F252 | 6 | 130,321,359 | Zm00001d037579 | binding | FT(0.05); HT(3.08); #L(-0.02); UE(0.05); BR(-0.08); LL(0.64); TKW(5.66) | FT(0.43); HT(2.34); #L(-0.02); UE(0.05); LE(0.1); BR(0.02); LL(1.02); TKW(8.26); EW(0.55) | FT(0.8) |

## Discussion

In this work, we exploited a unique material derived from an artificial selection experiment on plants cultivated in agronomic conditions to explore two essential aspects of observed phenotypic shifts: the effects of mutations with respect to their standing versus *de novo* status, and the multitrait response to selection. We derived Saclay DSEs by applying truncation selection for flowering time in two maize inbred lines for 18 generations. From each of the two inbreds, two late and two early families were derived as independent replicates of the same selection scheme. Saclay DSEs display several peculiarities: effective population sizes are small (*Ne <* 4, (Desbiez-Piat *et al*. 2021)); selection intensity is strong (1% of 500 progenitors selected per family); plants are reproduced through selfing. Despite High Drift-High Selection (HDHS), phenotypic evolution is substantial, continuous (Durand *et al*. 2010, 2015) and sustained by both standing variants and a constant flux of *de novo* mutations (Durand *et al*. 2010; Desbiez-Piat *et al*. 2021). Hence, considering a mutation rate of 30 *×* 10^*−*9^ (Clark *et al*. 2005), and with more than 1000 loci involved in maize flowering time (Romero Navarro *et al*. 2017), along with a median mRNA length of 6000 bp (Jiao *et al*. 2017), we anticipated approximately 0.36 mutations per individual per generation that is 0.36*×*500 = 180 incoming *de novo* mutations per generation per family. This is a coarse estimation that does not consider the genomic variation in mutation rates (Monroe *et al*. 2022).

Here, we first quantified the contribution of standing variation and *de novo* mutational variance to the observed selection response and monitored the fate of polymorphisms through generations. Second, we used common garden experiments to evaluate simultaneously plants from the 18 generations of selection addressing how *G×E* interactions had contributed to the response to selection, how selection on flowering time had impacted the evolution of correlated traits, and finally to which extent pleiotropy had shaped the phenotypic space exploration.

### Two successive yet distinct modes of adaptation are at play in Saclay DSEs

We used an extension of the animal model proposed by Wray (1990) that explicitly accounts for the increase relatedness caused by incoming mutational variance (Wray 1990) to analyse the response to selection in Saclay DSEs Yearly Measurement (DSEYM) and Common Garden experiments (DSECG). In line with previous results (Durand *et al*. 2010, 2015; Desbiez-Piat *et al*. 2021), we found that the observed strong selection response in Saclay DSEs can be decomposed in two successive phases with first, short term fixation of standing polymorphisms and second, a response sustained by the constant flux of incoming mutations (Fig. 1, Fig. 4).

The relative importance of these two adaptive phases measured as the proportion of *de novo* mutational variance *vs*. standing variance (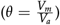) was consistent with a reduced initial standing variation in MBS compared to F252. Consistently, we detected fewer SNPs in MBS (46 SNPs) than in F252 from RNAseq data (7030 SNPs), a result corroborated by previous estimates from 50K SNP data of initial residual heterozygosity roughly 10-fold higher in proportion in F252 than MBS at generation 0 (Bouchet *et al*. 2013). Note that our genome-wide estimates from RNAseq data at generation 1, after a single generation of selfing and selection, revealed estimates around 0.1% in MBS and F252 (Table 2). The flux of incoming mutational variance, estimated through mutational heritabilities for flowering time 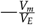 the ratio of mutational variance over residual variance— was unprecedentedly high (Table 3 and Table S6), an order of magnitude higher than the upper bound of previously measured estimates in inbred lines (10^*−*2^ and 10^*−*4^ (Keightley 2010)).

We investigated the allele fixation dynamic, considering both standing and *de novo* variants. The initial phase of the selection response was characterized by the fixation of standing genetic variants. Because standing genetic variation was not entirely fixed over the course of 18 generations, as demonstrated in MBS (Fig. S11), it is possible that the mutational variance estimates might include some of the fixation of recalcitrant heterozygosity. However, our heterozygosity estimates indicate that this effect may be minor, as only 6.4% of the heterozygous markers at G0 were fixed between generation 13 and generation 18, while 33.4% were fixed in the initial adaptive phase between generation 1 and 13.

Interestingly, the preservation of recalcitrant heterozygosity regions across different genetic backgrounds, families, and generations suggests that these regions were maintained in a heterozygous state due to selective mechanisms as previously described in Brandenburg *et al*. (2017). This might explain why these regions do not have a significant impact later in the selection response to flowering time. Additionally, this effect appears to be transient, as the fixation times were relatively short, with 3-4 generations in MBS and 5-6 generations in F252 (Fig. S15).

According to the selection limit theory for small effective population sizes (Robertson 1960), these fixation times aligned with the point at which a plateau was reached for the breeding values in both MBS and F252 backgrounds from the BLUPM model (Fig. 1). Fixation times were consistent with the modeling results calibrated on MBS data (Desbiez-Piat *et al*. 2021). In F252, the times to fixation were actually longer than expected under pure genetic drift, suggesting that clonal interference impaired the selection of beneficial variants in our selfing regime. Clonal interference is known to slow down the fixation of beneficial mutations in bacteria (Gerrish and Lenski 1998; Desai and Fisher 2007; Park and Krug 2007), consistent with theoretical work indicating that the loss of weaker adaptive alleles during a first selective sweep prevents the fixation of multiple other mutations (Hartfield and Glémin 2016). The absence of clonal interference in the MBS background was likely explained by reduced standing variation. Contrasting with the first adaptive phase, we expected no clonal interference in the second phase because fixation times are smaller than waiting times for new beneficial mutations to arise in the population (Gerrish and Lenski 1998). As predicted, the four *de novo* mutations fixed within the expected times (Desbiez-Piat *et al*. 2021).

### Variance estimates and limits of the model

Caution must be exercised when interpreting estimates of mutational heritability in our experimental setting due to the influence of both standing and mutational variances on the total phenotypic variance. As a result, the residual variance may not serve as a wholly accurate proxy for the total phenotypic variance Keightley (2010). Because standing variance decreased over time, we faced the challenge of addressing this issue, which might also have affected previous estimates. However, by considering the residual variance as a stable denominator, we aimed to mitigate the effects of this issue. In a related study, Durand *et al*. (2010) discovered that mutational heritabilities calculated for the first seven generations were notably lower (approximately *≈*2*×*10^*−*2^) than our estimates, an outcome most likely due to the predominance of standing variation fixation in the first generations. On the other hand, mutational heritabilities calculated in the present study for all other traits were two orders of magnitude lower than for flowering time, hence falling within the range of common observations of 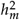 (Keightley 2010).

One assumption of the BLUPM model, which could potentially inflate the estimates of the mutational variance is the assumption of constant mutational input through generations under the Gaussian incremental model (Clayton and Robertson 1955; Kimura 1965; Walsh and Lynch 2018). This assumption has been found to be particularly relevant when effective recombination is limited as observed in Saclay DSE’s (Charlesworth 1993; Walsh and Lynch 2018). Nevertheless, because beneficial mutations fixed randomly across generations, resulting in mutational gaps across successive generations and bursts of response to selection (See Fig. S3 in Desbiez-Piat *et al*. (2021)), the model smoothed incoming mutational variation over generations. The BLUPM model also assumes that all mutations are transmitted to offspring, overlooking the significance of segregational variance within families which is known to account for half of the total additive variance in a panmictic population. While this bias certainly exists, its extent remains uncertain, as we expect the segregational variance to be approximately equal to 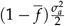 under selfing, with 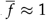 being the average inbreeding coefficient (Walsh and Lynch 2018). Additionally, due to the rapid fixation of adaptive mutations in Saclay DSEs, this bias is likely transient – but see Barton *et al*. (2017) for a possible inclusion of mutations and their segregational variance in the infinitesimal model based on House of Cards approximation. Finally, the Saclay DSEs do not take into account interactions between alleles, such as dominance or epistasis, which are known to play a crucial role in the response to selection (Durand *et al*. (2012) and discussion below). This omission may add complexity to the adaptive process by increasing the stochasticity through the effect of drift (Dillmann and Foulley 1998) that might be partly captured in the additive mutational variance.

### Distribution of fixed mutational effects

From DSEYM, we approximated both the distribution of effects of all incoming mutations and the one of selected mutations. We found an enrichment in beneficial mutations in the selected offspring of a progenitor. Consistently, previous simulations showed that high stochasticity promoted the fixation of small effect beneficial mutations in this High Drift-High Selection regime (Desbiez-Piat *et al*. 2021). Simulations also predicted the fixation of unfavorable mutations (6% only), but to a lesser extent than what we observed (25% in MBS and 17% in F252). Several non-exclusive hypotheses can be formulated here. First, our model considered only additive variance terms. But, epistasis might be a key player of the selection response in our DSEs as shown by earlier work (Durand *et al*. 2012) and the multivariate distribution of incoming mutations varying between early and late populations Fig 6. In other words, epistasis can promote the fixation of favorable mutations in their context of appearance, which later appear as unfavorable in a modified genetic context. In the same vein, directional epistasis and allele specificity makes the order of mutations critical as has been shown in *E. coli* where the fixation of the first mutations determines the effect and fixation of the following ones through the modification of their genetic context of appearance (Plucain *et al*. 2014). Furthermore, in the case of negative epistasis —when the mean mutational effect increases as fitness decreases— selection for low trait values can be impaired (Silander *et al*. 2007). In Saclay DSEs, a possible explanation for selection response asymmetry could reside in a fitness decrease in early families that in turn, would respond less to selection than the late ones in DSEYM (Durand *et al*. (2015) and Fig. 1). The second explanation for the excess of fixation of unfavorable mutations is the environmental fluctuations through generations of selection (Fig. S18) where a favorable mutation occurring in one environment might appear in the later generations as unfavorable (Chen and Zhang 2020). For example, mutations that are unfavorable on average across environments but that confer a transient beneficial advantage might be more likely to fix than mutations that are always neutral or beneficial (Cvijović *et al*. 2015). Such effects of antagonistic pleiotropy on fitness have been shown to be key players in local adaptation (Savolainen *et al*. 2013; Scarcelli *et al*. 2007; Chen and Zhang 2020).

### Evaluation environments do not mirror past selection environments

Previous simulation results pinpointed the importance of micro-environmental effects on the stochastic fixation dynamics of *de novo* mutations under HDHS (Desbiez-Piat *et al*. 2021). However the model did not take into account *G×E* interactions, nor did DSYEM allow for the quantification of its impact on phenotypic shifts. Here, the implementation of a two-years common garden experiment revealed the preponderance of GxE interactions, which impacted the selection response for flowering time in several respects.

A comparison between DSEYM and DSECG revealed the fixation of environment-dependent unfavorable mutations during the selection response. The differences between selected and non-selected mutations were eliminated in the evaluation environments (Table 4 and Fig. S20), indicating that some beneficial mutations in DSEYM became unfavorable in DSECG. This suggests that the same process applies between generations within DSEYM, with changes in allelic effects between selection years, as evidenced *e*.*g*. in *Drosophila* (Rudman *et al*. 2021; Kapun *et al*. 2021). Supporting these findings, the identified mutation on chromosome 2 in the ML1 family occurred in Zm00001d002125, also known as RPD2, which has been demonstrated to regulate flowering-time genes in an environment-dependent manner through siRNA-directed DNA methylation (RdDM) (Pikaard *et al*. 2008). Interestingly, when examining the selection response decomposition, we observed some antagonism between initial standing variation and mutational effects, especially in early flowering populations influenced by GxE interactions. This means that either fixed standing variation or *de novo* mutations compensated for the decrease in flowering time over generations in DSECG (Fig. 4), suggesting a certain degree of compensatory epistasis (Rojas Echenique *et al*. 2019) modulated by the evaluation environment. Regarding the total selection response, *G×E* interactions were significant in DSECG but mainly visible through scale effects, which magnified the asymmetry of the selection response against earliness observed in DSEYM. This could be interpreted because the two years of evaluation (2018 and 2019 on the Saclay plateau) were characterized by a hot summer (Fig. S18), causing early flowering populations to flower in a shorter time window, leading to the observed asymmetrical selection response.Additionally, a further comparison between DSEYM and DSECG environment-specific breeding values (Fig. S17) resulted in numerous rank changes. This finding is in line with the results of Choquette *et al*. (2023), which highlighted the significant impact of adaptive plasticity in the total selection response measured in a complete reciprocal transplant experiment conducted in eight environments of selection.

Our results more broadly question the use of the modern environment to understand past selection events that have shaped the current architectures of traits highly sensitive to environmental conditions. Hence trait expression in the environment of past selection may differ drastically from that of the present-day environment. This is well illustrated in teosintes whose phenotypic characteristics in late glacial-like climatic conditions (low temperature and CO2 levels) differed markedly from that of present-day conditions (Piperno *et al*. 2015). Such plasticity could have drastic effects on association studies particularly for climate-sensitive quantitative traits such as flowering time (Bergelson and Roux 2010). In line with these findings, for the gene Zm00001d047269, encoding the EARLY FLOWERING 4 protein, we found association with flowering date neither in 2019 or nor in DSYEM, despite a significant association in 2018 and strong effects observed on flowering time (3.11 DTF in 2018, 3.46 DTF in 2019, 3.61 DTF at DSYEM). It is remarkable that despite the homogeneity of our genetic backgrounds we found such discrepancy across our GWA studies. A previous study on flowering time in*A. thaliana* has pinpointed to a lack of reproducibility of GWA owing to genetic heterogeneity across populations (Lopez-Arboleda *et al*. 2021). Here we argue that the varying effects of alleles across environments are also key to explain the discordance among GWA studies performed across multiple environments in accordance to (Li *et al*. 2010), and the lack of power to detect significant associations when combined in single-data sets. Our results also raise the question of the application of association mapping in populations submitted to High-Drift High-Selection: on one hand many small-effect mutations follow adaptive trajectories comparable to neutral mutations, on the other hand, large effect mutations fix quickly and are detectable within populations only over restricted time windows. Hence, while the identification of variants underlying phenotypic variation in populations submitted to HDHS constitute a major goal for breeders and evolutionary geneticists, the discovery of alleles “that matter” might remain elusive (Rockman 2012). Note that despite these limitations, we managed to associate the four *de novo* mutations detected to a trait correlated to flowering time suggesting that strong pleiotropic effects may actually facilitate the detection of associations.

### Pleiotropic mutational input drives Saclay DSEs correlative selection response

The patterns observed suggest distinct degree of pleiotropy between the standing variants selected during the first phase of adaptation, and the incoming mutations selected during the second phase of adaptation. Among 456 detected standing variants, 99 were significantly associated to at least one trait, but only 11.3% were associated to more than two traits indicating a lower degree of pleiotropy than that observed for four *de novo* mutations which associated with 2, 1, 5, and 8 traits (Table 5). While we may have limited statistical power to formally test this difference, it is worth noting that this pattern resembles what we observed concerning the pairwise-trait correlations. Specifically, the correlations between standing variations effects Fig. S25 were indeed overall less conserved between years and genetic backgrounds – with the exception of the FVL family which fixed a standing variant of major effect in the first few generations (Durand *et al*. 2012)– than those computed on the predicted effects of incoming mutations (Fig. 6).

Correlative responses in Saclay DSEs are best explained by developmental constraints. Hence, correlations for traits such as flowering time, leaf length and plant height are predicted by developmental models of maize architecture (Zhu *et al*. 2014; Vidal and Andrieu 2020). These models also predict a correlation between blade length and plant size by the successive dependence of the blade length on sheath length, which impacts internodes length, and in turn plant height. Because developmental constraints may emerge from fixation of pleiotropic alleles (Hughes and Leips 2017), our results are consistent with: a first phase of adaptation with fixation of small-to mild-effects standing variants characterized by a restricted pleiotropy, allowing for a stochastic exploration of the phenotypic space (Fig. 5 and Fig. S22); and a second phase, where the fixation of strong effect of intermediate-to high-level pleiotropic *de novo* mutations restrict the phenotypic space exploration. In the omnigenic model of adaptation, this second class of mutations corresponds to the ones that are first fixed and display large effects both on the focal trait and other traits (Liu *et al*. 2019; Boyle *et al*. 2017). Corroborating this expectation, Frachon *et al*. (2017) showed that a small number of QTLs with intermediate degrees of pleiotropy drove adaptive evolution in nature. Note that selection itself may also contribute to reinforce genetic correlations during this second phase. Simulations of the evolution of gene regulatory network under directional selection have indeed depicted a trend towards gain of regulatory interactions and a global increase in the genetic correlations among gene expressions (Burban *et al*. 2022). This is because the rewiring of network is easier to achieve by adding connections on existing ones. Our results demonstrate a dynamic change in the patterns of correlations associated with a change in the source of polymorphism transitioning from standing variation to new mutations.

## Data availability

The generation 1 30x DNA-seq Illumina data for this study have been deposited in the European Nucleotide Archive (ENA) at EMBL-EBI under accession number PRJEB64332 for maize inbred line F252 (https://www.ebi.ac.uk/ena/browser/view/PRJEB64332) and PRJEB64333 for maize inbred line MBS847((https://www.ebi.ac.uk/ena/browser/view/PRJEB64333). The generation 13 RNA-seq data have been deposited in the Gene Expression Omnibus database from the National Center for Biotechnology Information under Bio-project: PRJNA531088. The 25 GEO accessions used are SRX5646859 to SRX5646883. The generation 18 RNA-seq data have been deposited in the European Nucleotide Archive (ENA) at EMBL-EBI under accession number PRJEB64524 (https://www.ebi.ac.uk/ena/browser/view/PRJEB64524). Raw phenotypic data, BLUP values, raw and imputed genotyping data, pedigrees and climatic records were deposited on recherche.data.gouv.fr with the DOI: https://doi.org/10.57745/J0ZRRI.

## Acknowledgments

We are extremely grateful to Randall Wisser for the insightful discussions. We also thank Hélène Corti for quality control work of the genotyping data and Johann Joets for help for preparing and uploading the data files to ENA.

## Supplementary material

**Figure S1.**
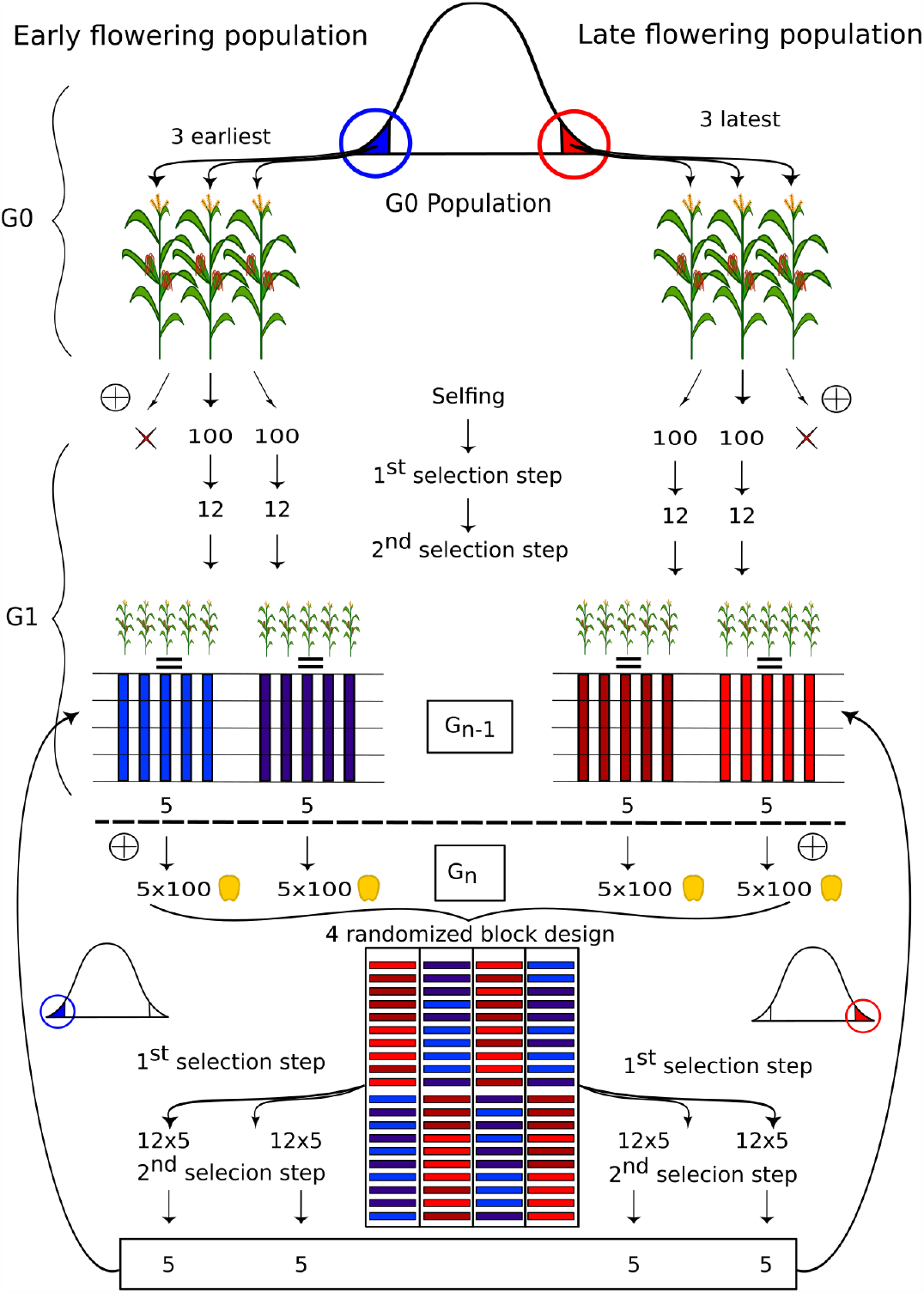
Experimental scheme of Saclay Divergent Selection Experiment for flowering time. For clarity a single scheme is shown but was replicated for the two DSEs. Starting from an inbred G0 population with little standing variation (*<* 1% residual heterozygosity (Durand *et al*. 2015)), the three earliest (resp. latest) flowering individuals represented in blue (resp. red) were chosen based on their offspring phenotypic values as the founders of two families forming the early (resp. late) population. For the subsequent generations, 10 (*≈*5 per family) extreme progenitors were selected in a two step selection scheme among 1000 plants. More specifically, 100 seeds per progenitor were evaluated in a four randomized-block design, *i*.*e*. 25 seeds per block in a single row. In a first selection step, the 3*×*4 = 12 earliest (resp. latest) flowering plants among the 100 plants per progenitor were selected in a first step. Then in a second selection step, 10 (*≈*5 per family) individuals were selected within each population based on both flowering time and kernel weight and the additional condition of preserving two progenitors per family from the previous generation.

**Figure S2.**
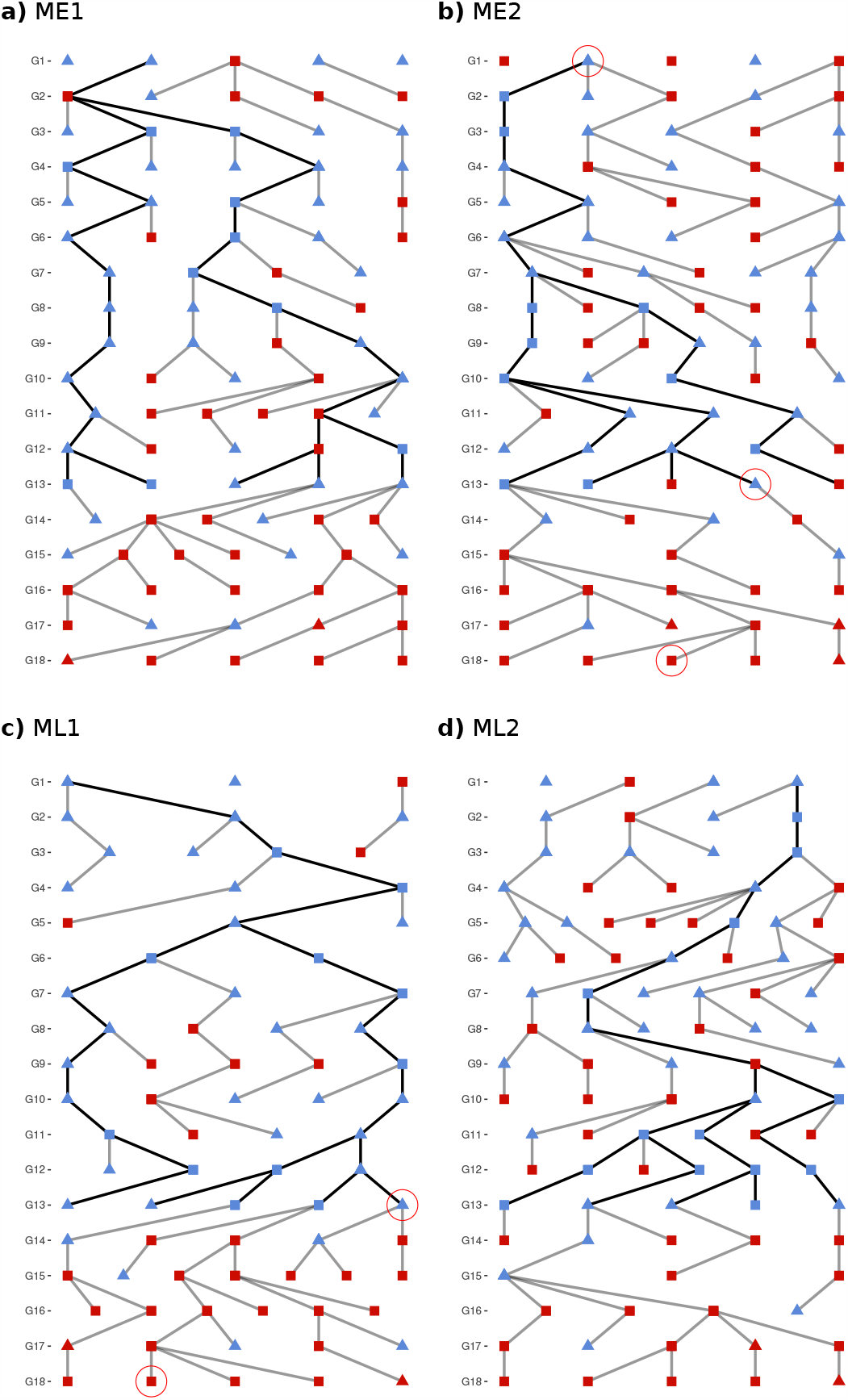
MBS family pedigrees from G1 to G18. The two early families ME1 a) and ME2 b), and the two late families ML1 c) and ML2 d) are presented. Each node corresponds to a progenitor selected at a given generation. Each edge indicates the relationship between a progenitor and its offspring. Individuals represented by a triangle were phenotyped in 2018 and 2019. Squares represent progenitors not included in the common gardens. Individuals represented in blue were genotyped with KASPar. Individual circled in red in ME2 generation 1 was DNA sequenced. The two G13 individuals circled in red in ME2 and ML1 were sequenced through RNAseq and used for SNP detection and the two generation 18 individual circled in red were sequenced through RNAseq. Thick black lines indicates the ancestral path of generation 13.

**Figure S3.**
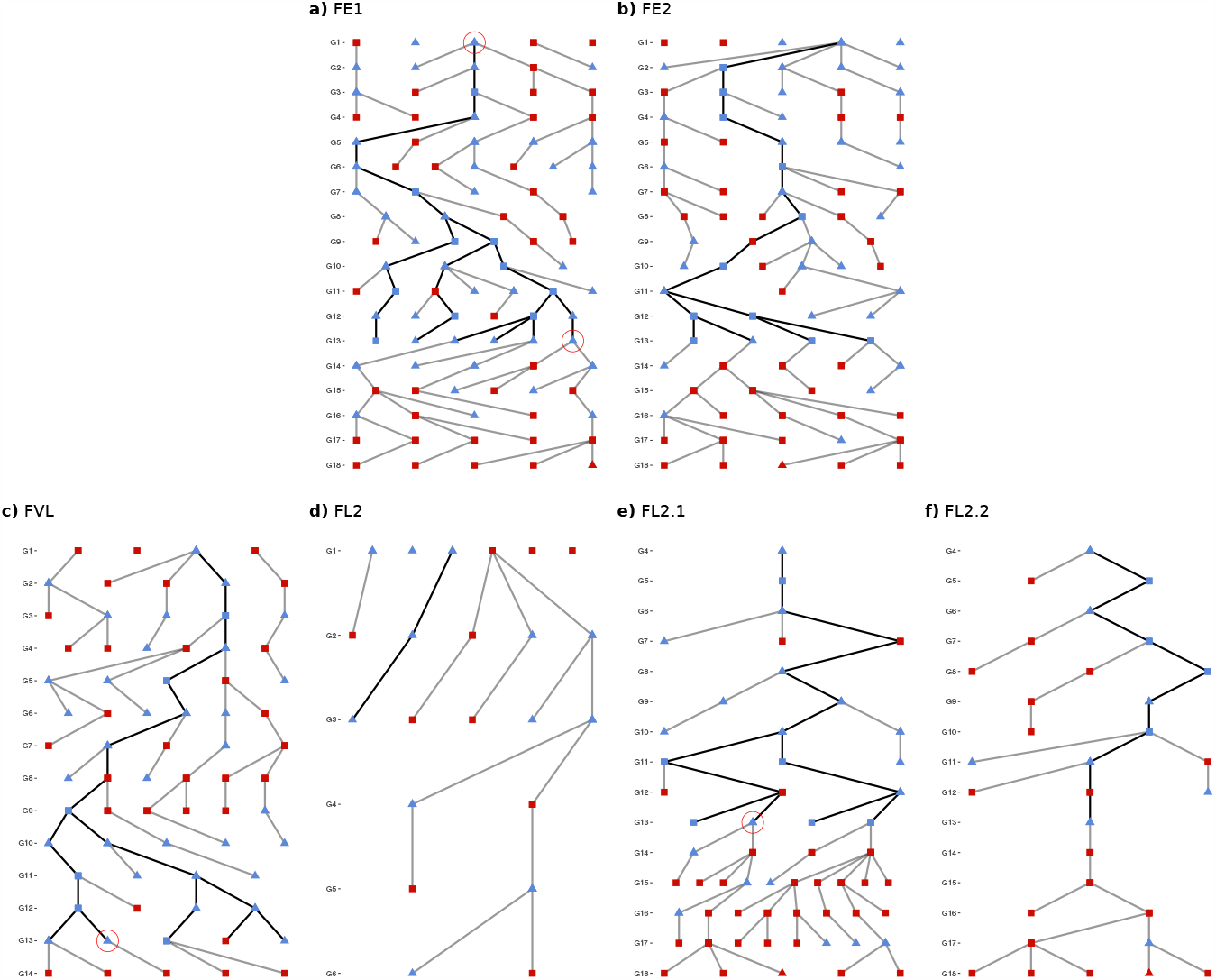
Graphical representation of F252 families pedigree relationship. Two early families FE1 a), FE2 b) together with one late family, FVL c), are represented. FVL c) could not be maintained after G14 (Durand *et al*. 2012). Both FL2.1 d) and FL2.2 e) were derived from a single individual from FL2 f) at G3, after FVL was discarded. Each node corresponds to a progenitor selected at a given generation. Each edge corresponds to a filial relationship between a progenitor and its offspring. Individuals represented by a triangle were phenotyped in 2018 and 2019. Squares represent progenitors not included in the common gardens. Note that FVL individuals were phenotyped in common garden only in 2018. Individuals represented in blue were genotyped with KASPar. Individual circled in red in FE1 generation 1 was DNA sequenced. The three G13 individuals circled in red in FE1, FVL and FL2.1 were used for SNP detection. Thick black lines indicates the ancestral path of generation 13.

**Table S1.** Number of reads per genotype after trimming and filtering for low quality reads and rRNAs removal following the procedure described in Tenaillon et al. (2019) for generation 13.

| Genotype | Number of libraries | Total number of pooled plants | Number of reads |
| --- | --- | --- | --- |
| ME2 G1 | 1 | 4-5 | 255 411 879 |
| FE1 G1 | 1 | 4-5 | 251 968 819 |
| MEE (ME2) G13 | 6 | 150 | 92 504 789 |
| MLL (ML1) G13 | 5 | 178 | 59 399 604 |
| FEE (FE1) G13 | 4 | 100 | 93 216 514 |
| FLL (FL2.1) G13 | 5 | 146 | 96 708 953 |
| FVL G13 | 3 | 56 | 66 890 125 |
| ME2 G18 | 22 | 22 | 754 466 679 |
| ML1 G18 | 26 | 26 | 875 166 361 |

**Table S2.** Distribution of detected SNPs and clusters of SNPs over the maize chromosomes for MBS (46) and F252 (7030)

| Line | Chromosome | 1 | 2 | 3 | 4 | 5 | 6 | 7 | 8 | 9 | 10 |
| --- | --- | --- | --- | --- | --- | --- | --- | --- | --- | --- | --- |
| MBS | #SNPs | 0 | 2 | 0 | 0 | 0 | 2 | 0 | 1 | 0 | 41 |
| F252 <sup>a</sup> | #SNPs | 895 | 37 | 465 | 61 | 179 | 1652 | 73 | 998 | 122 | 804 |
|  | # clusters | 11 | 10 | 12 | 11 | 6 | 12 | 13 | 12 | 7 | 5 |

<sup>a</sup> For F252 clusters of SNPs were defined by k-means algorithm using euclidean distances between SNPs positions.

**Table S3.**
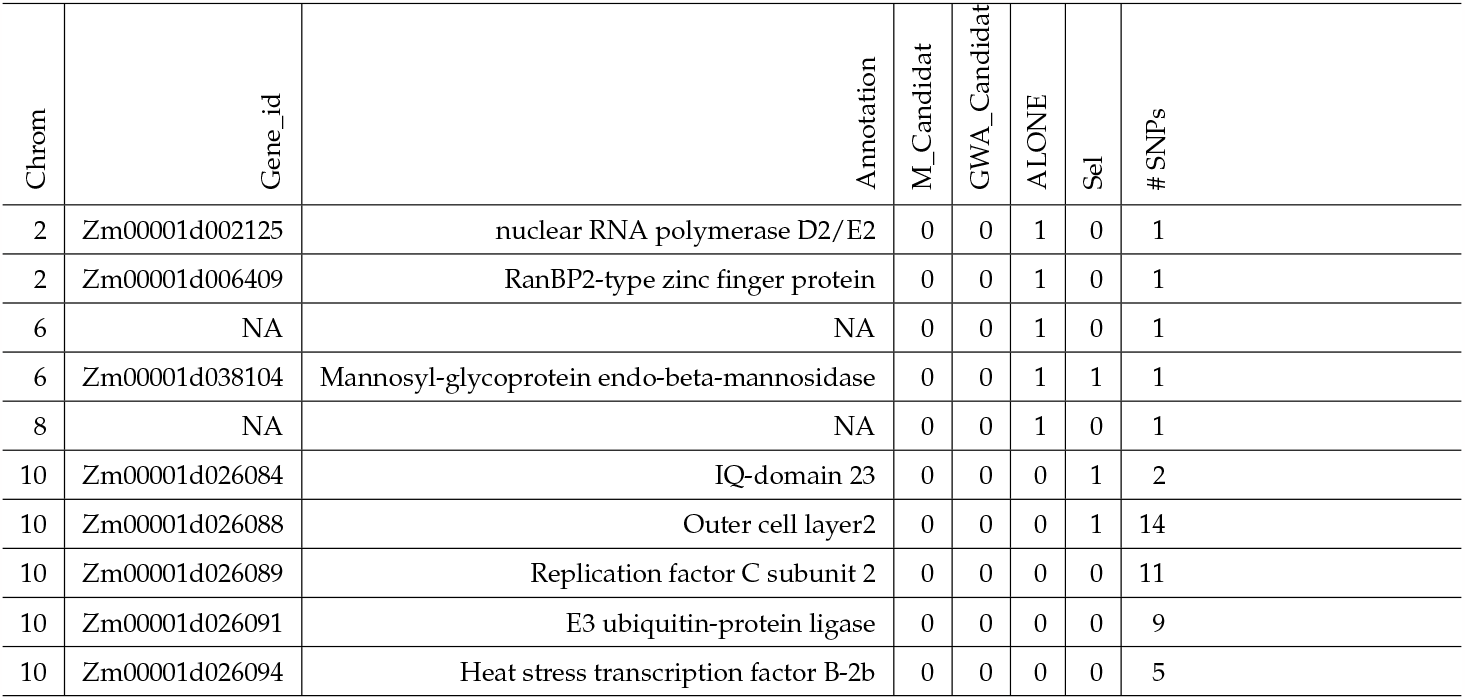
MBS SNP distribution among categories of genes.

**Table S4.** F252 SNPs distribution among chromosomes.

| Chromosome | 1 | 2 | 3 | 4 | 5 | 6 | 7 | 8 | 9 | 10 |
| --- | --- | --- | --- | --- | --- | --- | --- | --- | --- | --- |
| # of SNPs | 91 | 10 | 55 | 4 | 24 | 131 | 4 | 82 | 12 | 67 |

**Figure S4.**
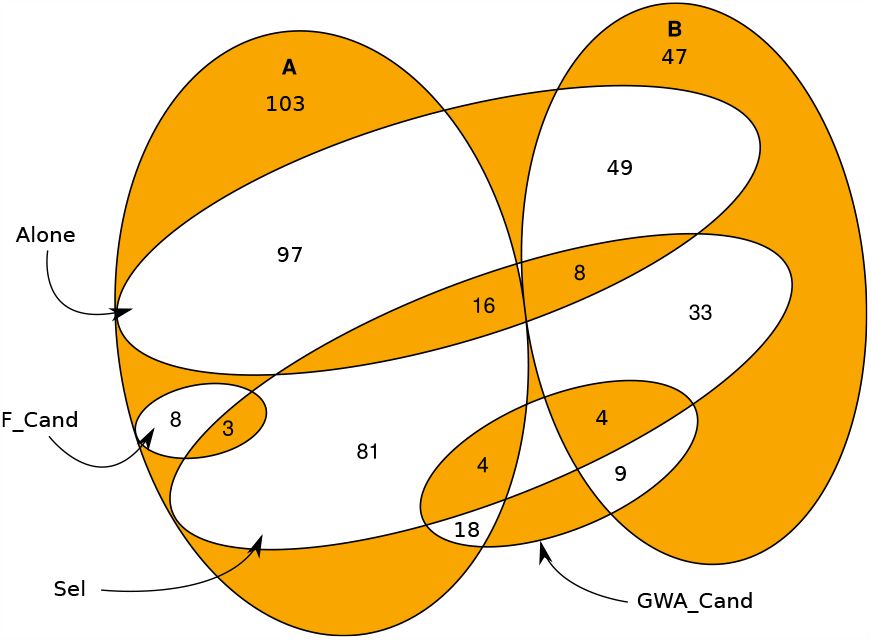
Genotyped SNPs distribution among categories in F252. Categories “F_Candidate”, “GWA_Candidate”, “Sel” correspond to the one defined in Tenaillon *et al*. (2019) such that: “F_Candidate” corresponds to SNPs in known flowering time genes in maize, “GWA_Candidate” corresponds to SNPs in genes associated to flowering time variation in a panel of 4,471 inbred lines/landraces (Romero Navarro *et al*. 2017), “Sel” corresponds to SNPs in genes detected as differently expressed between Early and Late G13 progenitors in the RNA-Seq. “Alone” were SNPs found isolated in 20kb regions, and Group A corresponds to SNPs differing between FE1 and {FVL and FL2.1} while Group B corresponds to SNPs differing between {FE1 and FVL} and FL2.1.

**Figure S5.**
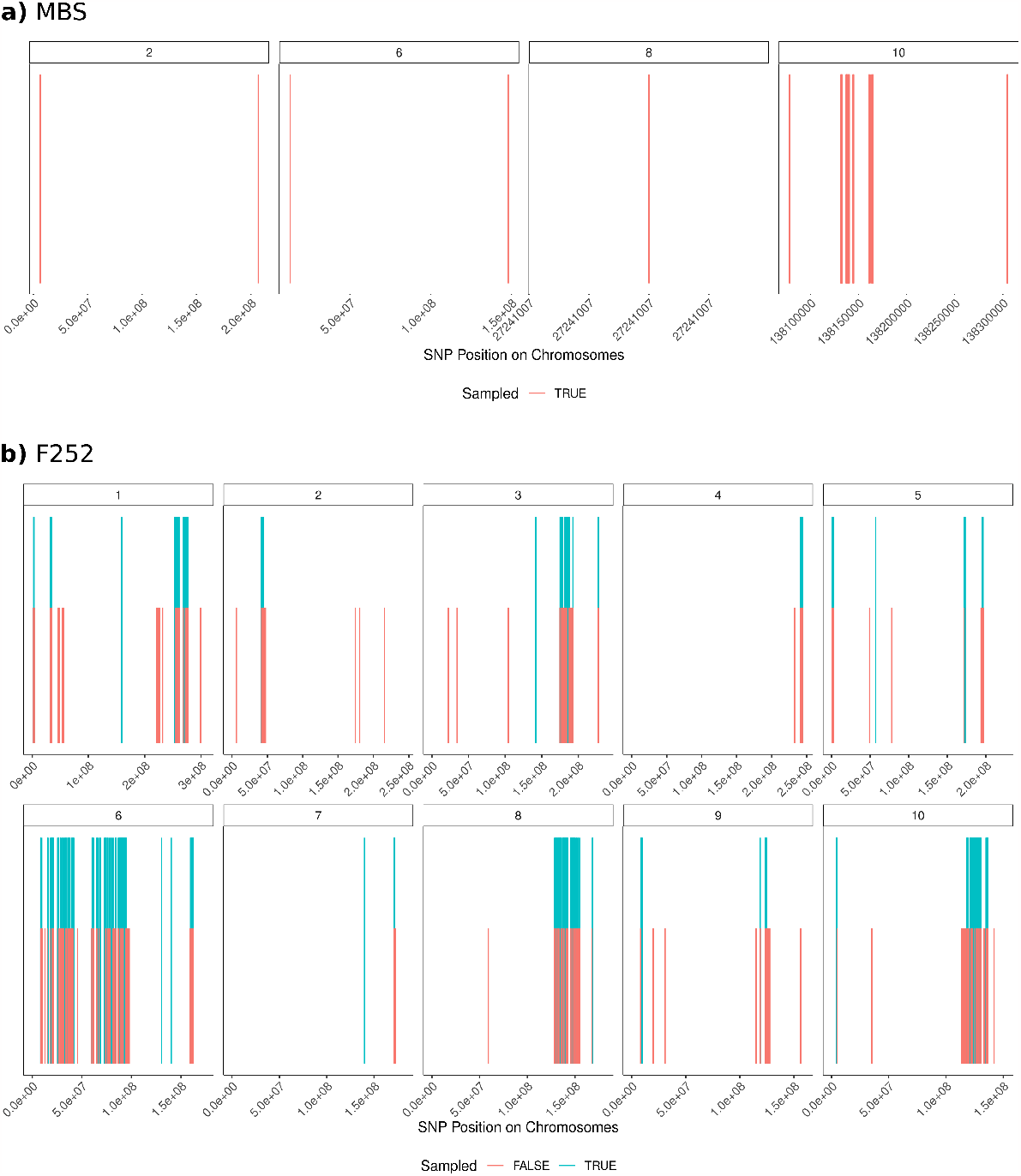
Genomic positions of the detected SNPs a) Each bar represents the position of a SNP. For MBS, all 46 SNPs were sampled for KASPar™ genotyping. **b)** For F252, 480 SNPs (in blue) were sampled among the 7,030 detected (in red).

**Figure S6.**
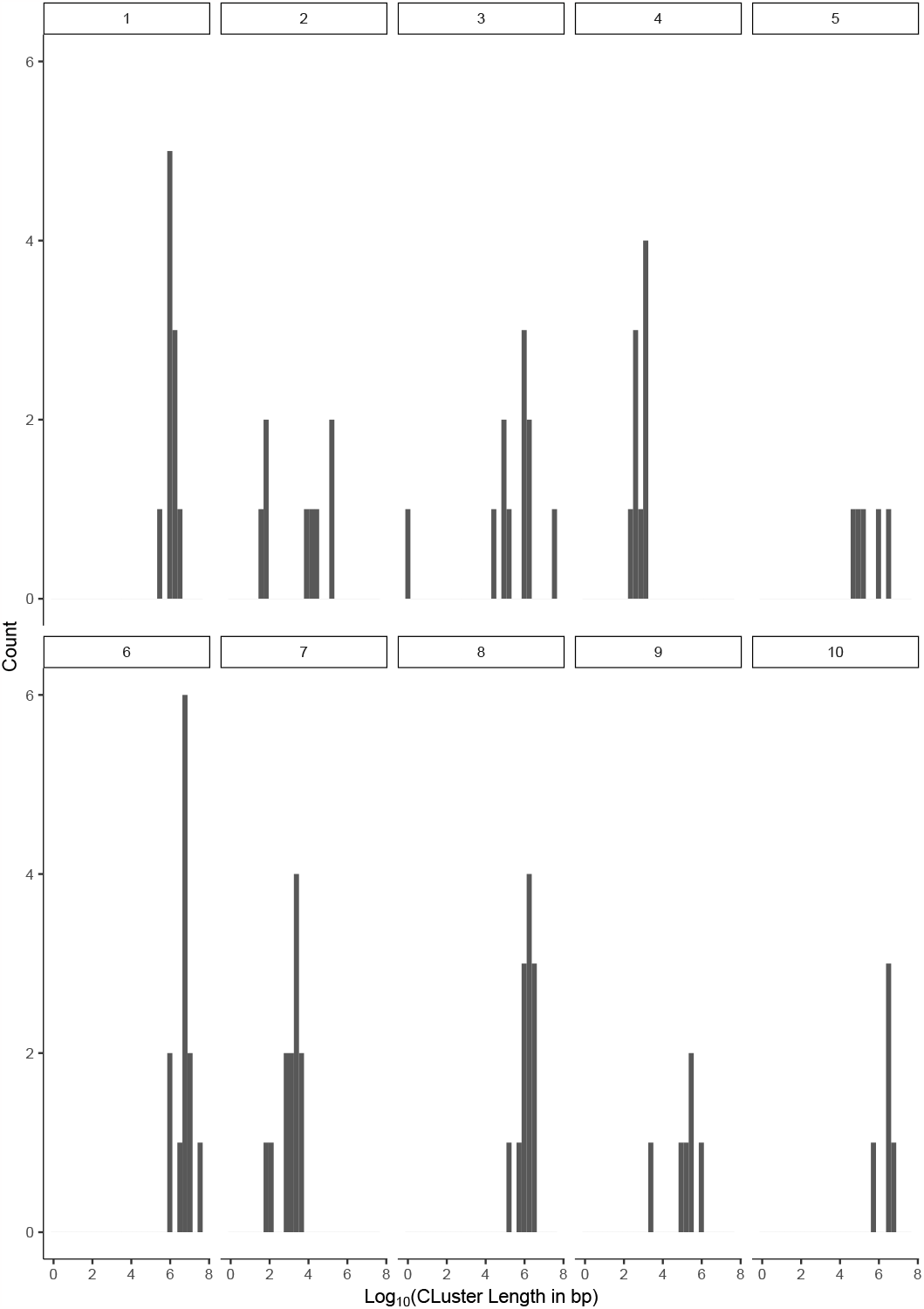
Distribution of the cluster length (*log*_10_(#*bp*)) per Chromosome in F252 after k-means clustering.

## Text S1. Inferring genotypes from sparse KASPar^(tm)^ genotyping data and pedigree relationships

We produced a sparsed genotyping matrix using KASPar™ genotyping, so that genotypic information was available only for a subset of all progenitors. From this subset we aimed at inferring all genotypes of the genealogy until generation 18_*th*_.

In order to do so, we developed a likelihood model, and used a parsimony algorithm developped by Durand *et al*. (2015) to infer missing data in the matrix of genotypes with the highest likelihood at every given SNP. Below we describe the likelihood function that calculates for a given locus, the probability of each genotype given the SNP genotypes of other individuals. This function considers (i) the genealogical information, (ii) the generations of selfing, (iii) the experimental error associated with KASPar genotyping.

### Heredity

Let *X*_*g,i*_ ∈ {0, 1, 2} be the random variable associated to the true genotype of individual *i* at generation *g* at a SNP biallelic locus. *X*_*g,i*_ can be associated to a probability law :

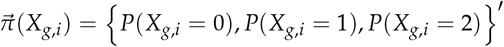

Conditionally to the parent 𝒜_*g,i*_, we have, for any realization *x*_*o*_ ∈ *{*0, 1, 2*}*:

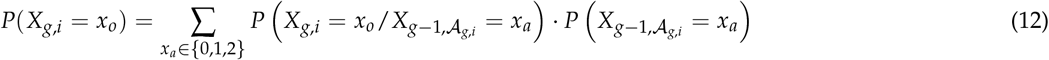

Individuals are reproducing through selfing every generation.

We supposed that mutations occured during meiosis at a rate *µ* per locus, per individual, per generation with equal probability of: *A* → *a* or *a* → *A*.

Selection occurred among the adult progenies, and we considered an additive fitness effect for allele *a*, such that: *w*_0_ = 1, *w*_1_ = 1 + *s* and *w*_2_ = 1 + 2*s*.

Hence, the frequency of selected genotypes *x*_*o*_ among the progenies of parent *x*_*a*_ is:

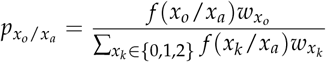

where *f* (*x*_*k*_ /*x*_*a*_) is given by Mendelian inheritance taking into account selfing and mutations. We established the (3 *×* 3) matrix *T* of probabilities of *X*_*g,i*_ given 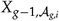 as indicated in the table below:

| Offspring | Parent |  |  |
| --- | --- | --- | --- |
|  | 0 (AA) | 1 (Aa) | 2 (aa) |
| 0 (AA) | $\frac{(1-\mu)^2}{1+2s\mu}$ | $\frac{1}{4+4s}$ | $\frac{\mu^2}{1+2s(1-\mu)}$ |
| 1 (Aa) | $\frac{2\mu(1-\mu)(1+s)}{1+2s\mu}$ | $\frac{1}{2}$ | $\frac{2\mu(1-\mu)(1+s)}{1+2s(1-\mu)}$ |
| 2 (aa) | $\frac{\mu^2(1+2s)}{1+2s\mu}$ | $\frac{1+2s}{4+4s}$ | $\frac{(1-\mu)^2(1+2s)}{1+2s(1-\mu)}$ |

so that,

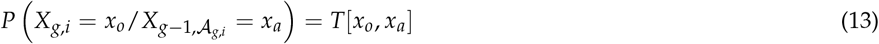

Using (12) and the *T* matrix,we have:

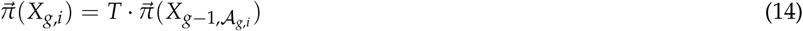

### Modeling SNP genotype

Let 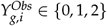 be the random variable associated to the observed SNP genotype of individual *i* at generation *g*. Because of both bulk genotyping and experimental errors, the SNP phenotype does not translate directly into the genotype. Let 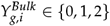 be the random variable associated to the true SNP genotype after bulk sampling, but before KASPar errors.

#### Bulk genotyping

For genotyping, each individual *S*_0_ of the genealogy has been selfed once to produce 25 *S*_1_ individuals, also selfed with seeds (*S*_2_) collected in bulk. We used DNAs from 15 *S*_2_ plants to infer the genotype of the *S*_0_ individual. This procedure may lead to ascertainment bias that occurs with a probability *p*_*b*_. For instance, 15 *S*_2_ plants generated from a *S*_0_ heterozygote may by chance be homozygotes for the *a* allele. This may occur with a probability 0.5 at each generation of selfing (0.25 in *S*_2_). Let

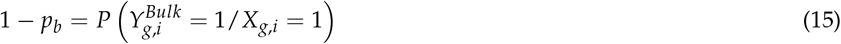

In the bulk of 15 *S*_2_ plants, we have *p*_*b*_ < 0.02. Because we performed many independent bulks, we modeled *p*_*b*_ as a random variable following an exponential distribution of parameter *λ*_*b*_ = 1/0.02 = 50.

We established (3 *×* 3) matrix *T*^*Bulk*^ of probabilities of 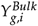 given *X*_*g,i*_ as indicated in the table below:

| $\gamma_{g,i}^{Bulk} =$ | $X_{g,i} =$ | | |
| --- | --- | --- | --- |
|  | 0 (AA) | 1 (Aa) | 2 (aa) |
| 0 (AA) | 1 | $\frac{p_b}{2}$ | 0 |
| 1 (Aa) | 0 | $1 - p_b$ | 0 |
| 2 (aa) | 0 | $\frac{p_b}{2}$ | 1 |

#### Accounting for experimental errors in KASPar genotyping

Experimental biases and misinterpretation of the fluorescence data may occur if:

- homozygous genotypes are widespread on the two-dimensional fluorescence plan. This may occur either because of the differential/variance around the fluorescence signal between the two allele-specific fluorophores,
- or because of the bulk DNAs and variation of allele counts which may translate into a continuum between homozygotes and heterozygotes.

Let *e* be the probability of misinterpretation of one allele of a biallelic SNP genotype. We can establish (3 × 3) matrix *T*^*Kasp*^ of probabilities of 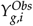 given 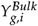 as indicated in the table below:

| $\gamma_{g,i}^{Obs} =$ | $\gamma_{g,i}^{Bulk} =$ | | |
| --- | --- | --- | --- |
|  | 0 (AA) | 1 (Aa) | 2 (aa) |
| 0 (AA) | $(1 - e)^2$ | $e(1 - e)$ | $e^2$ |
| 1 (Aa) | $2e(1 - e)$ | $e^2 + (1 - e)^2$ | $2e(1 - e)$ |
| 2 (aa) | $e^2$ | $e(1 - e)$ | $(1 - e)^2$ |

Hence, we have 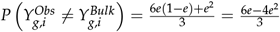.

#### From SNP phenotypes to SNP genotypes

Considering all possible sources of error (bulk genotyping and experimental biases), we have :

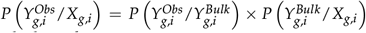, whose corresponding transition matrix is given by the product *T*^*Kasp*^ · *T*^*Bulk*^, which can be written:

| $\gamma_{g,i}^{Obs} =$ | $X_{g,i} =$ | | |
| --- | --- | --- | --- |
|  | 0 (AA) | 1 (Aa) | 2 (aa) |
| 0 (AA) | $(1 - e)^2$ | $2e^2p_b - e^2 - 2ep_b + e + 1/2p_b$ | $e^2$ |
| 1 (Aa) | $2e(1 - e)$ | $-4e^2p_b + 2e^2 + 4ep_b - 2e - p_b + 1$ | $2e(1 - e)$ |
| 2 (aa) | $e^2$ | $2e^2p - e^2 - 2ep_b + e + p_b/2$ | $(1 - e)^2$ |

Hence, we have :

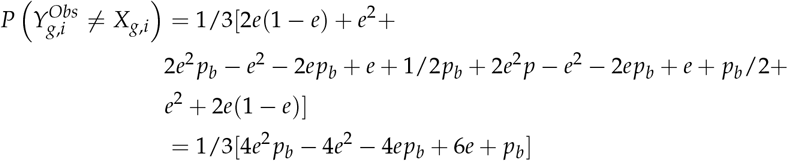

### Likelihood function of genotypes

We have described above how to estimate the probability of a phenotype knowing the genotype of the individual and also how to calculate a vector of probability of the 3 genotypes knowing the genotype of its parent. From there we were able to estimate the likelihood of a genotype of the genealogy knowing its SNP phenotype and its parent.

The parameters of the model are:

- *θ*_*hered*_ = (*µ, s*) that relates to heredity.
- *θ*_*exp*_ = (*p*_*b*_, *e*) that relates to the experiment.

The known information is 𝒜_*g,i*_ the parent of each individual of the genealogy. The random variables are 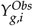 and 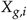. Using conditional probabilities, we have

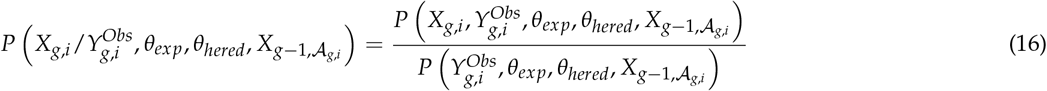

Because the SNP phenotype knowing the genotype does not depend on the parental genotype, and because *θ*_*hered*_ and 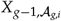 are independent, we have

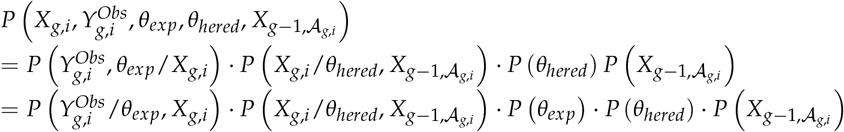

Similarly, the denominator of (16) can be written:

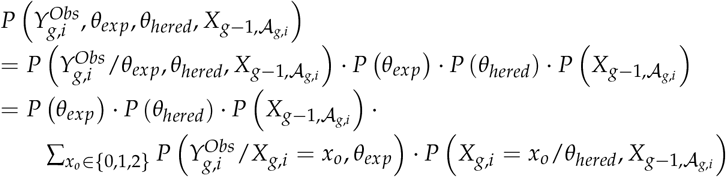

Therefore, we have,

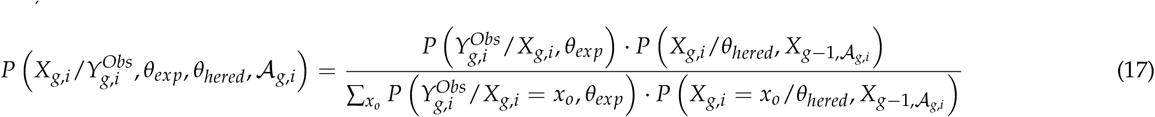

Equation (17) can be computed using the models described above. Furthermore, knowing the parent of each individual of the genealogy, conditional probabilities in (17) do not use any extra information and they are independent. Therefore, we extended this equation to all individuals of a given family in order to capture the genealogical information contained in related individuals. The family is characterized by a combination of genotypes whose likelihood is the product of the likelihood of each individual genotype

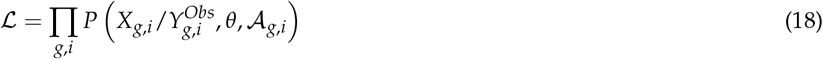

### Inference algorithm description

To perform the inference of missing genotypes we used the algorithm developed by Durand *et al*. (2015).

**Figure S7.**
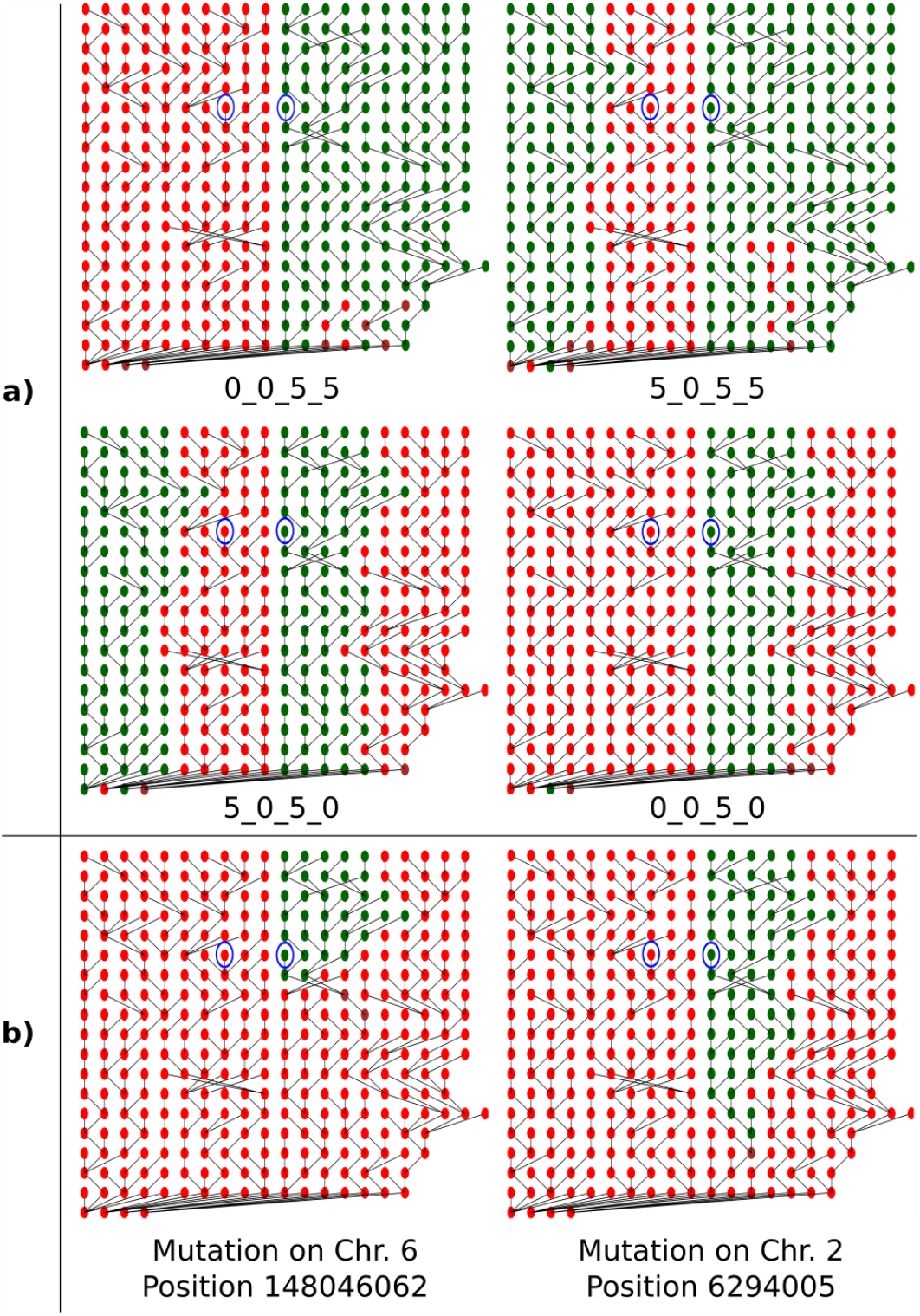
**Example of the main genealogical structure cases in MBS** for a) Standing variation and b) the two detected *de novo* mutations. In MBS, we expected under pure drift 4 main different cases for standing variants after conditioning on the SNP detection at generation 13. If we count the number of genotype *AA* in family ME1, 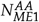, for example, respectively 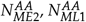 and 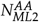, the 4 main cases (neglecting unfixed alleles within family) are:

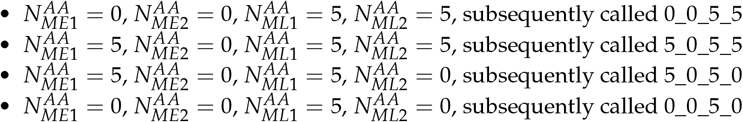

Example of simulated genealogies corresponding to these 4 cases, plus two simulated genealogies corresponding to *de novo* mutations are presented. Note that in MBS these 4 cases encompassed over 99.9% of the simulations for standing variation and the early occurring mutation 2_6294005 (at *G*_3_). *G*_18_ is at the top of each genealogy and *G*_0_ at the bottom, each individual is represented by a dot, whose color indicates the genotype (in red *aa*, in green *AA* and in brown *Aa*). Each line represents a kinship relationship. At *G*_18_ (top), each group of 5 individual belongs to one family, in order: ME1_ME2_ML1_ML2. Blue circles identify the *G*_13_ progenitors used for SNPs detection.

**Figure S8.**
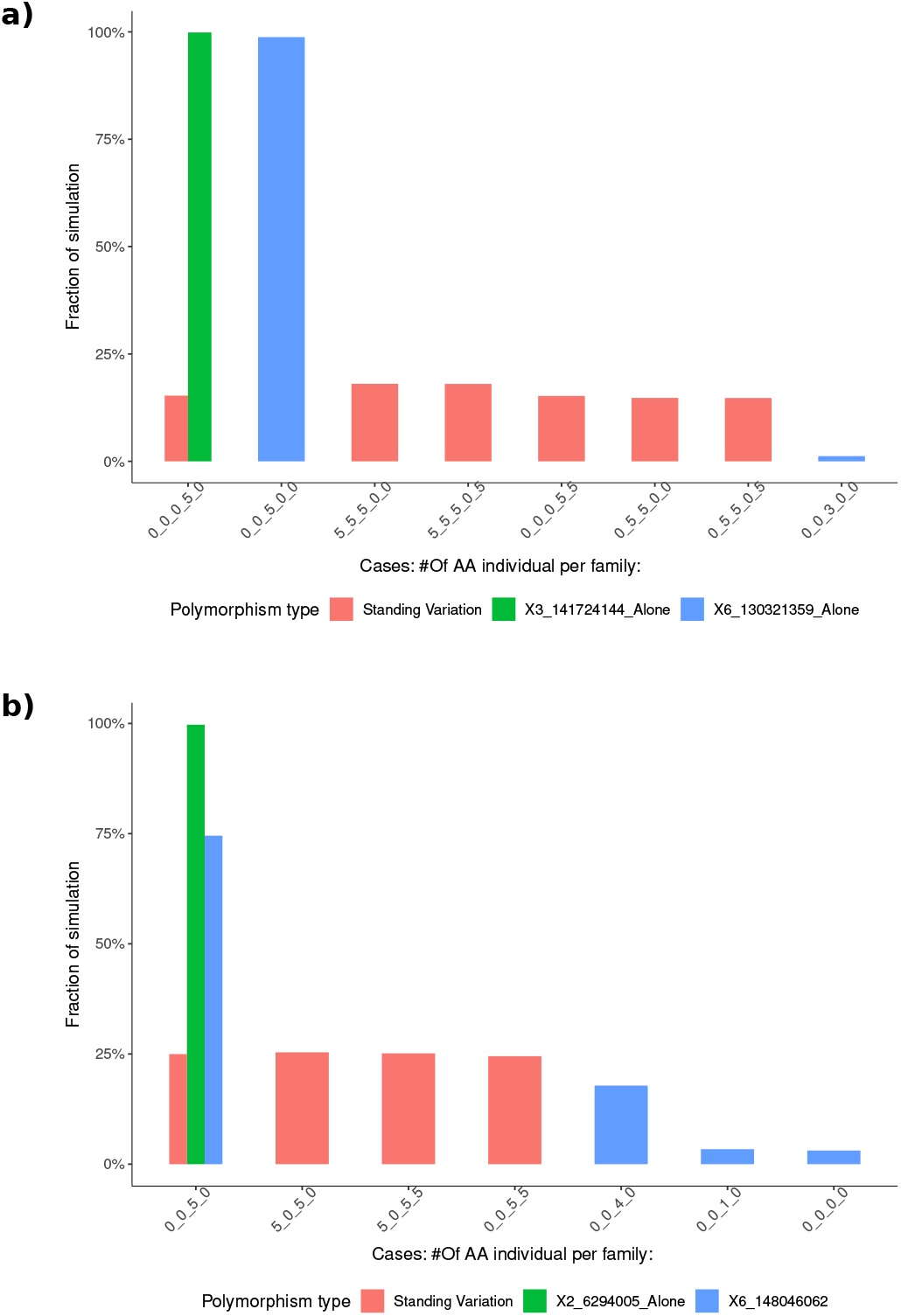
**Frequency distribution of simulated genealogical structures of segregating polymorphisms** in a) F252, and b) MBS. Cases are summarized by AA counts in FVL_FL2.1_FL2.2_FE1_FE2 for F252 and ME1_ME2_ML1_ML2 in MBS. For F252, the typology of possible cases for standing variants is composed of 6 main cases represented in red— each over 8% of the simulations— representing 97% of the possibilities. In MBS, the typology of possible cases for standing variants is composed of 4 main cases represented in red encompassing 99.9% of the simulations. In both cases, unrepresented cases (3% of the simulations in F252 and 0.01% of the simulations in MBS) corresponded to cases were standing variants were not fixed at generation *G*_18_. Green and blue bars corresponds to the two mutations detected in each backgrounds for the corresponding cases.

**Figure S9.**
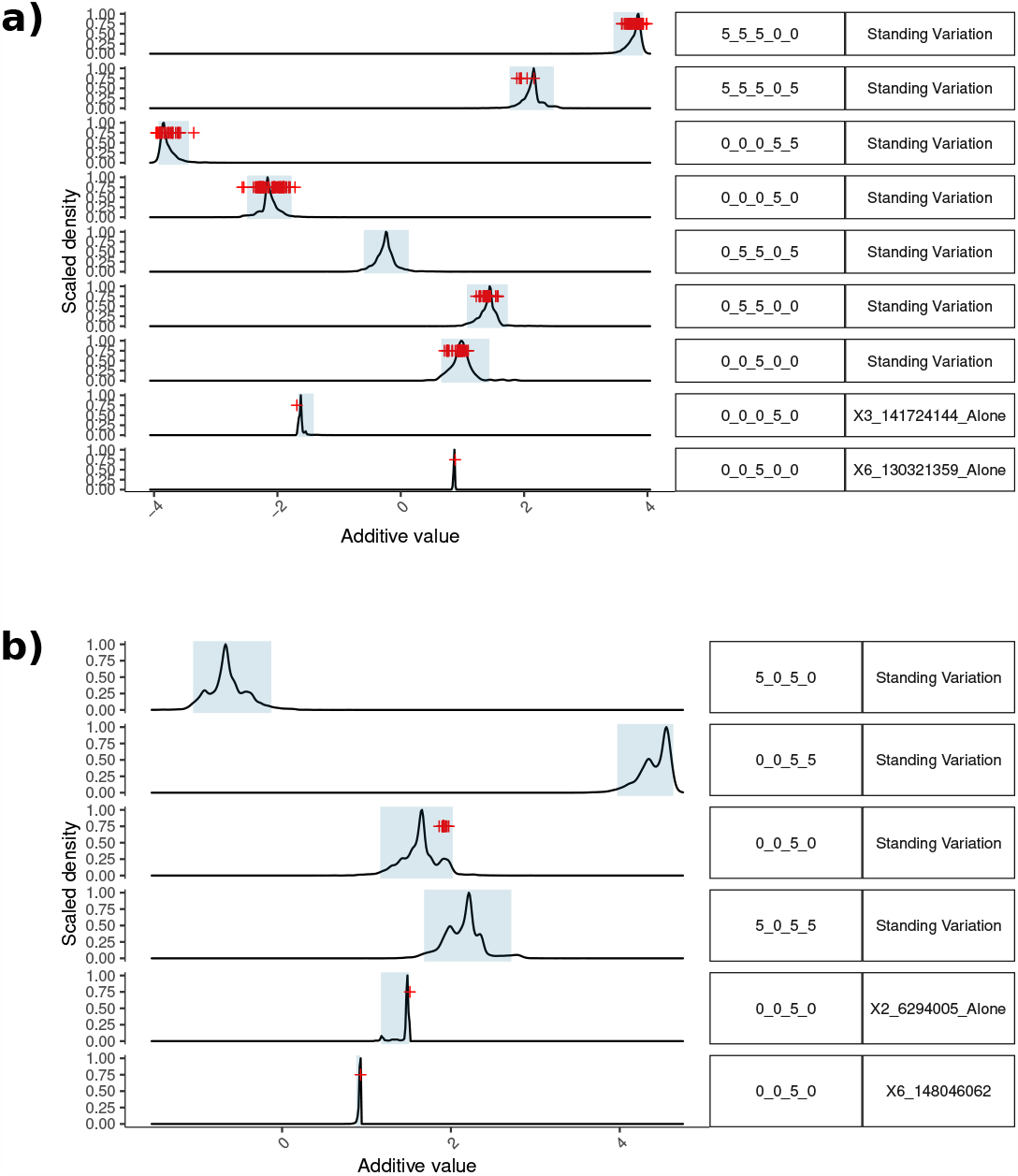
**Distribution of simulated and observed (red crosses)** *a* **values for flowering time (DSEYM) according to the main genealogy at** *G*_18_ **defined Fig. S8** in a) F252, and b) MBS. Shaded areas cover 95% of the corresponding distribution.

**Figure S10.**
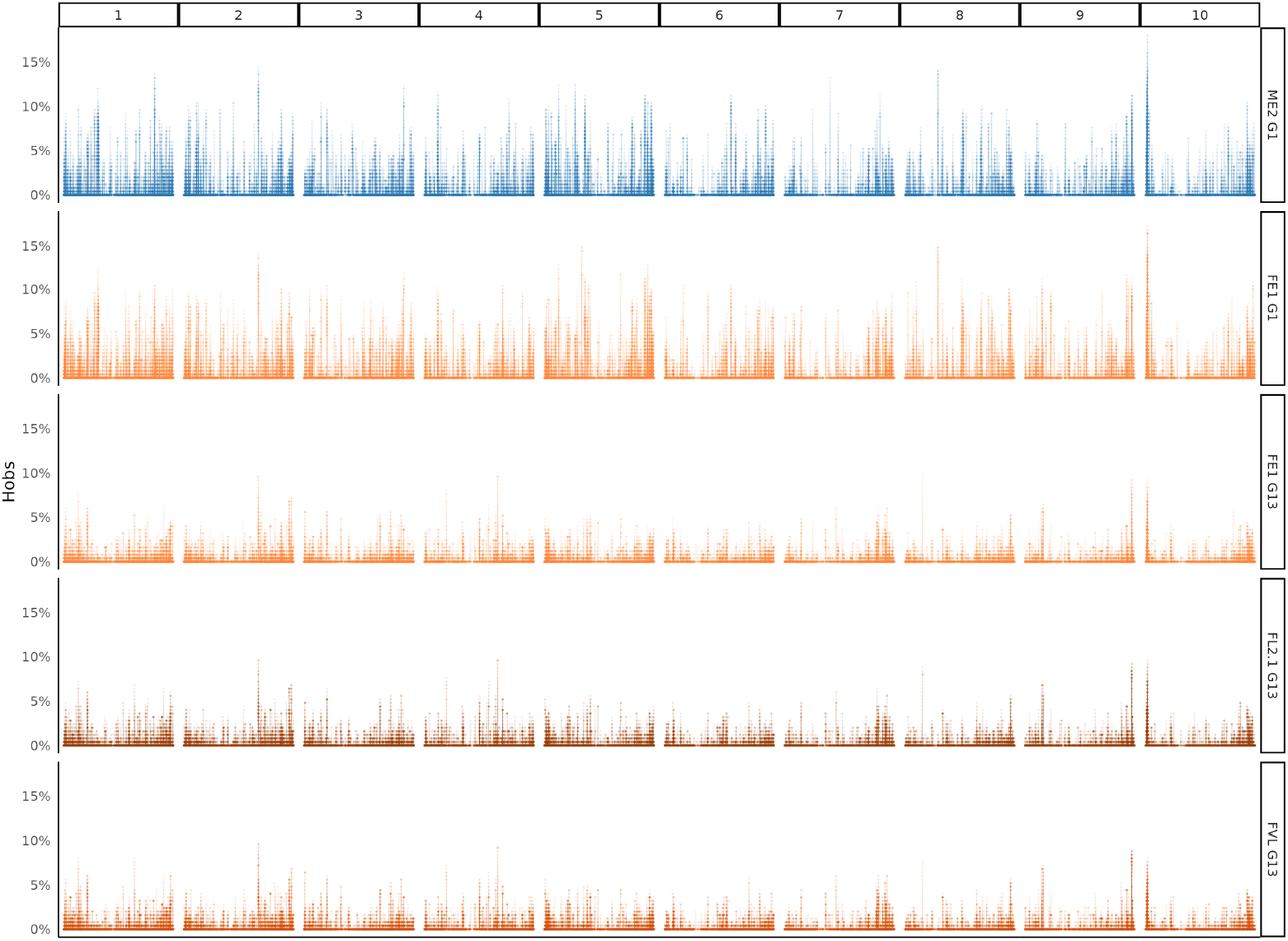
Distribution along whole chromosomes of the observed level of heterozygosity in F252, averaged by sliding window of length 250pb. Each bar corresponds to a region of length 250bp. Each row of graphic corresponds to a unique progenitor of generations 1 and 13. ME2 G1 is provided as an outgroup. Note that more than 95% of the measured regions had no heterozygous sites.

**Figure S11.**
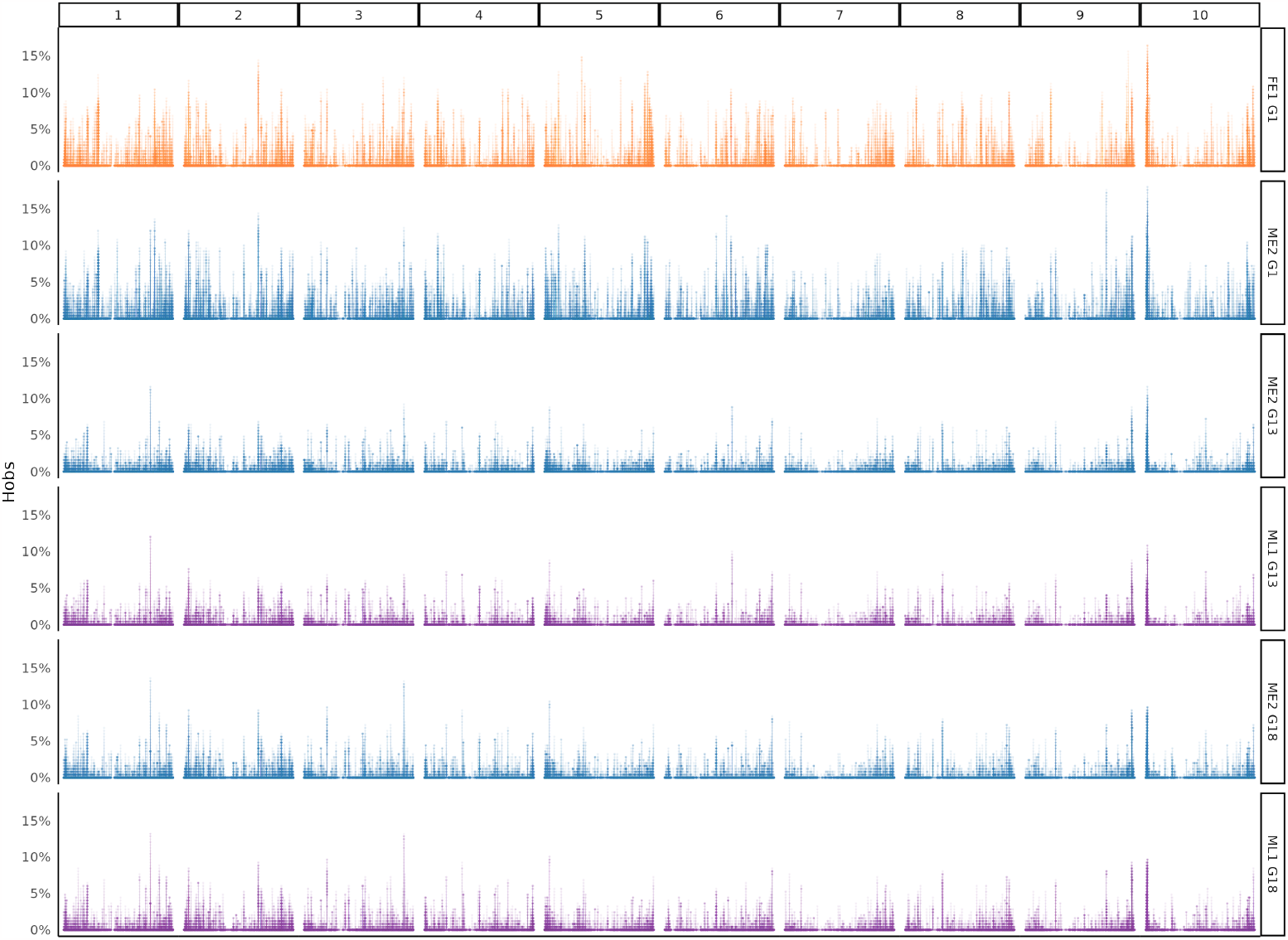
Distribution along whole chromosomes of the observed level of heterozygosity in MBS, averaged by sliding window of length 250pb. Each bar corresponds to a region of length 250bp. Each row of graphic corresponds to a unique progenitor of generations 1, 13 and 18. FE1 G1 is provided as an outgroup. Note that more than 95% of the measured regions had no heterozygous sites.

**Figure S12.**
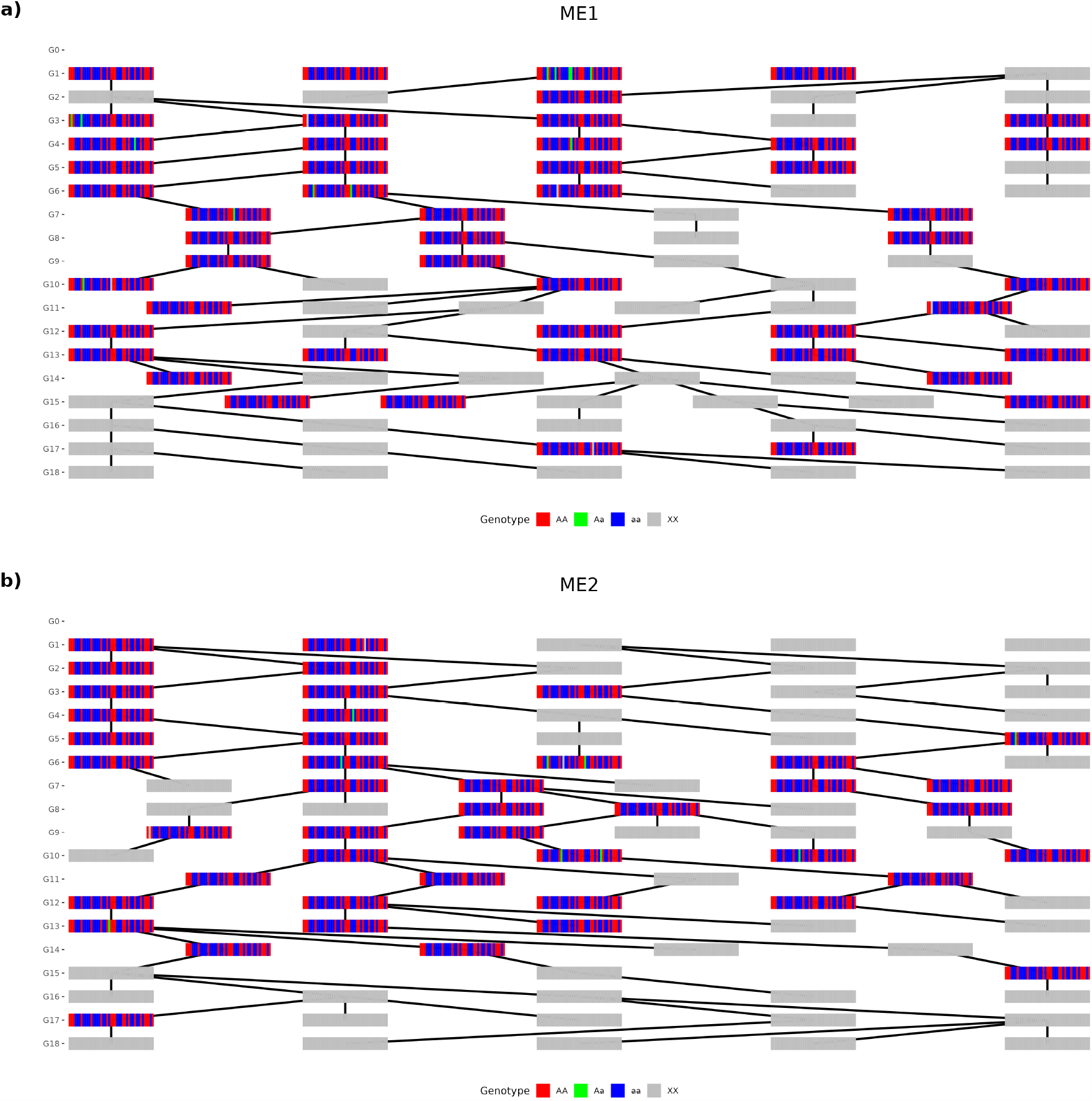
Multilocus representation of KASPar genotypic data along MBS pedigrees from G1 to G18. The two early families ME1 a) and ME2 b) are presented. Each multicolor bar-code represents a genotyped progenitor and grey bars indicate progenitors that were not genotyped. Each edge indicates the relationship between a progenitor and its offspring. Each bars of the bar-codes corresponds to a locus. Each line represents a kinship relationship.

**Figure S12.**
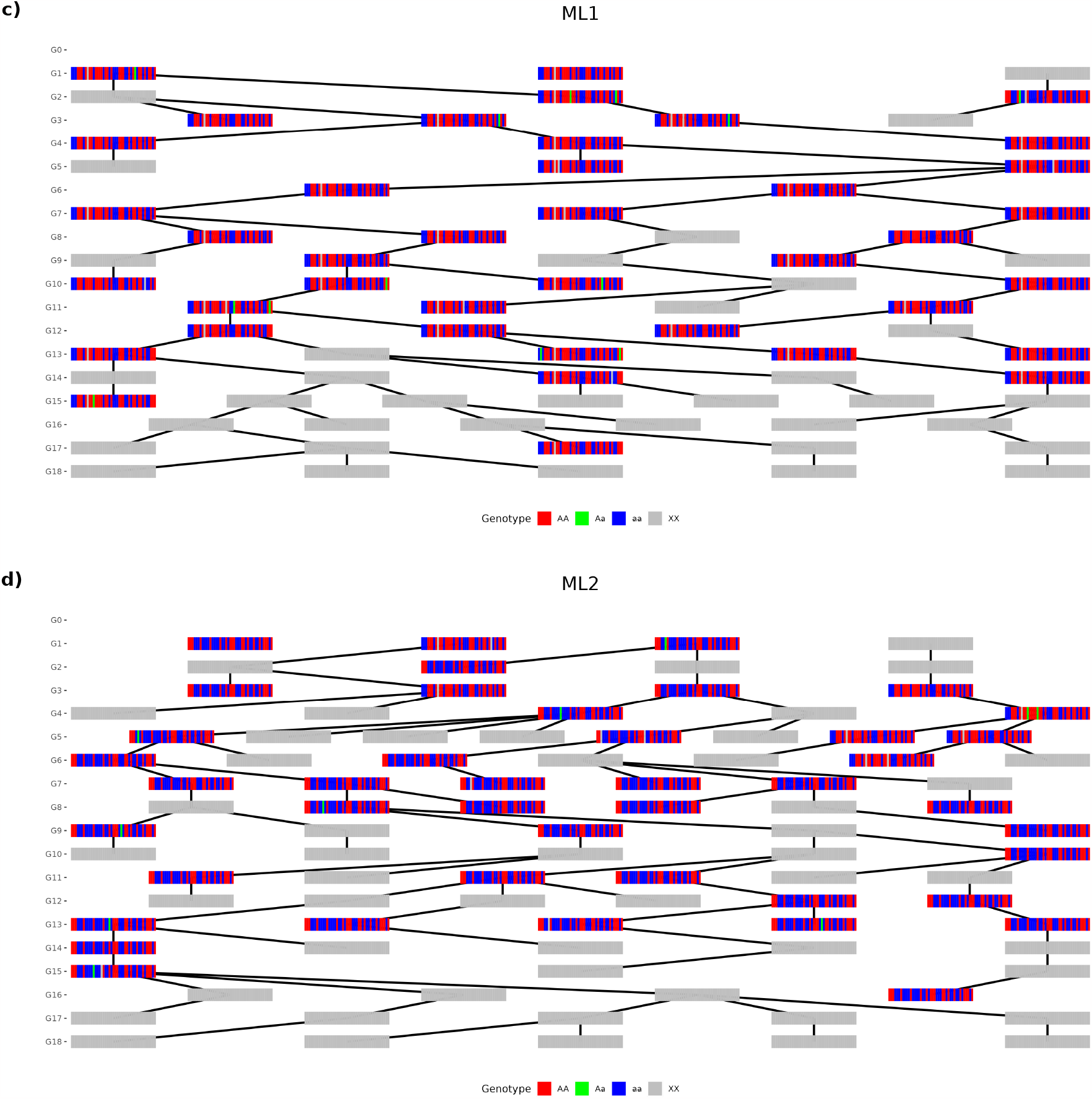
Multilocus representation of KASPar genotypic data along MBS pedigrees from G1 to G18. The two late families ML1 c) and ML2 d) are presented. Each multicolor bar-code represents a genotyped progenitor and grey bars indicate progenitors that were not genotyped. Each edge indicates the relationship between a progenitor and its offspring. Each bars of the bar-codes corresponds to a locus. Each line represents a kinship relationship.

**Figure S13.**
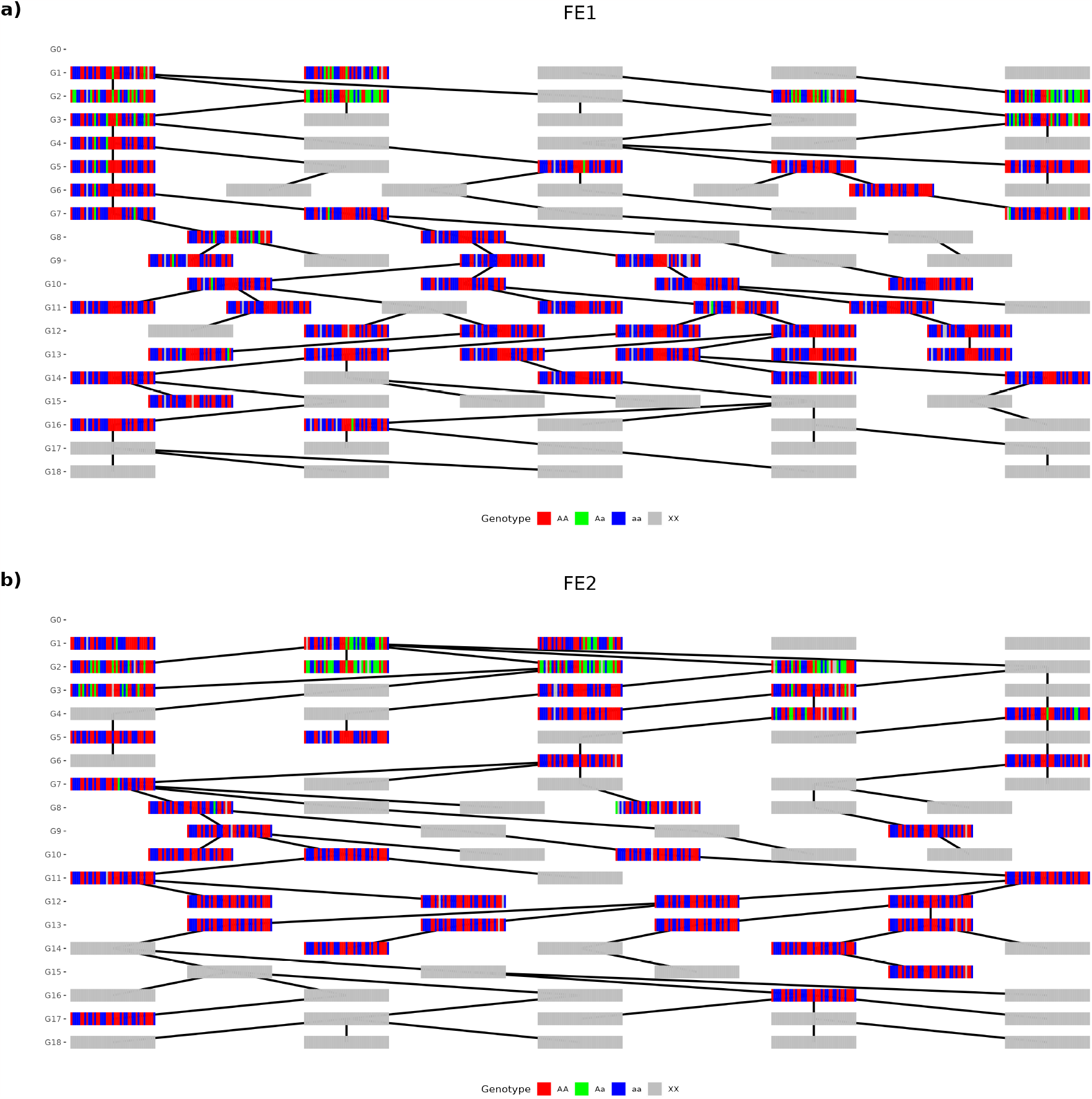
Multilocus representation of KASPar genotypic data along F252 pedigrees from G1 to G18. The two early families FE1 a) and FE2 b) are presented. Each multicolor bar-code represents a genotyped progenitor and grey bars indicate progenitors that were not genotyped. Each edge indicates the relationship between a progenitor and its offspring. Each bars of the bar-codes corresponds to a locus. Each line represents a kinship relationship.

**Figure S13.**
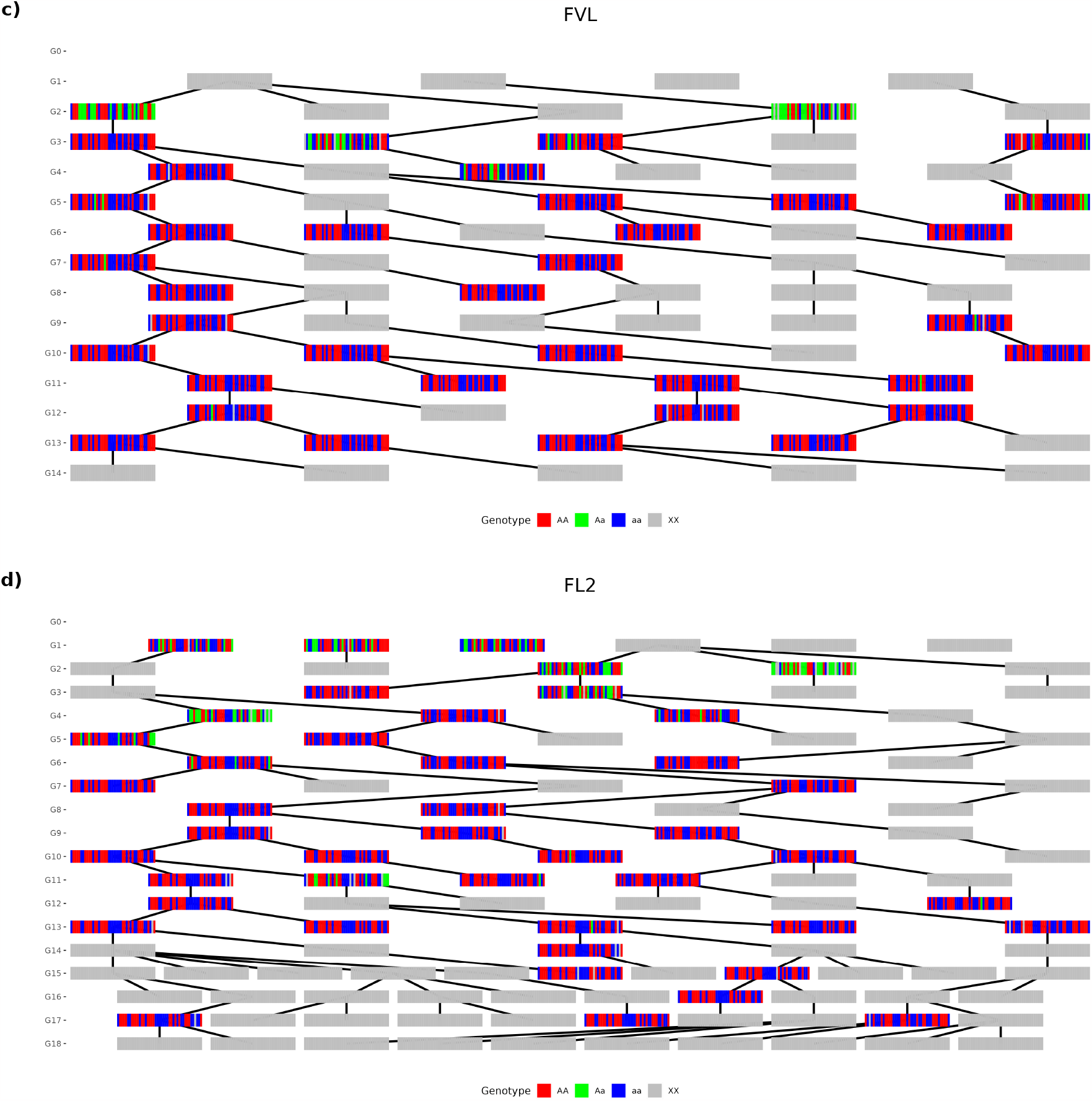
Multilocus representation of KASPar genotypic data along F252 pedigrees from G1 to G18. The two late families FVL c) and FL2 d) are presented. Each multicolor bar-code represents a genotyped progenitor and grey bars indicate progenitors that were not genotyped. Each edge indicates the relationship between a progenitor and its offspring. Each bars of the bar-codes corresponds to a locus. Each line represents a kinship relationship.

**Figure S14.**
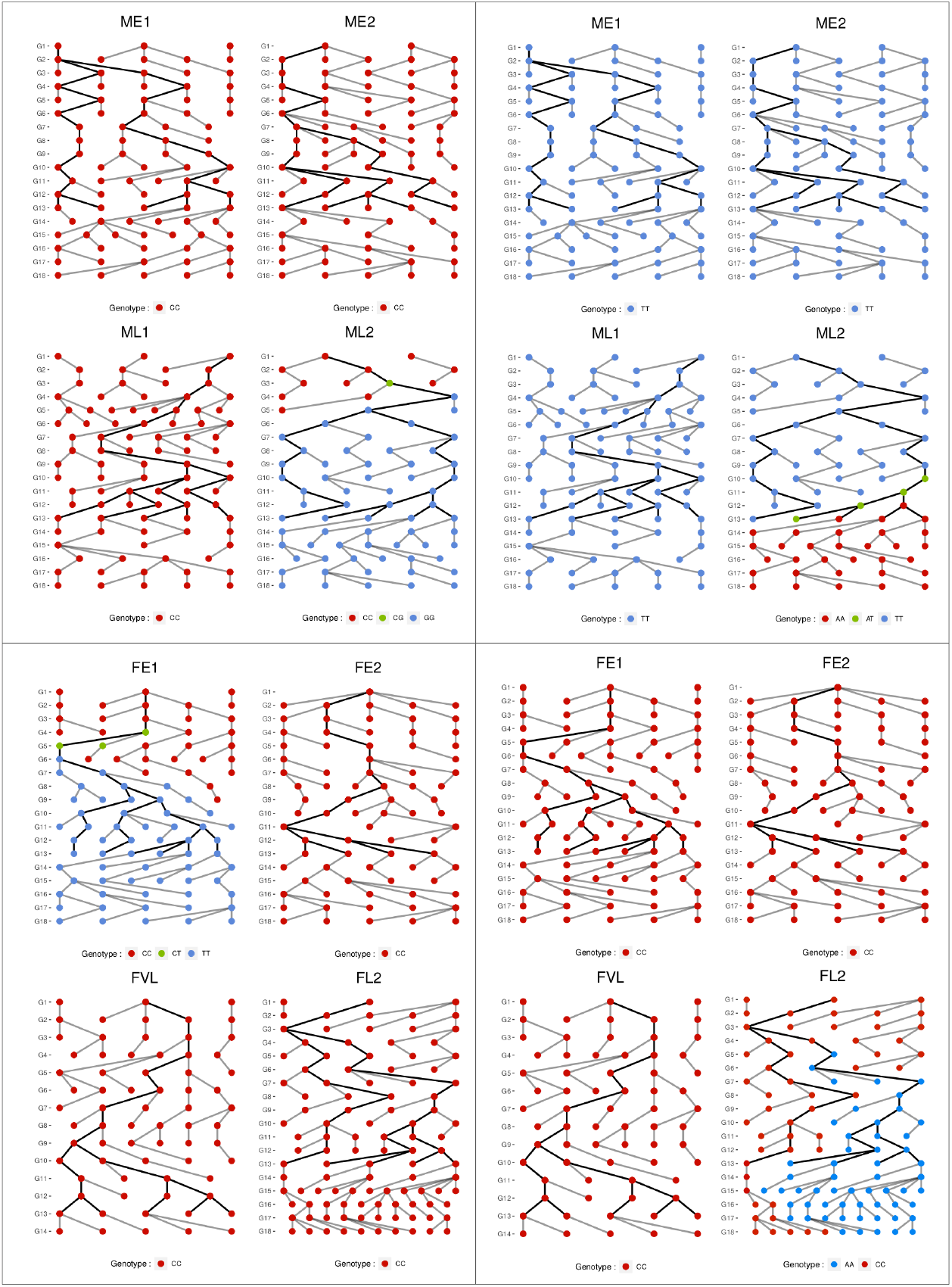
*De novo* mutation dynamics through generations within MBS and F252 families. Colors indicate the genotypes of the 4 detected mutations along the pedigrees. From top-left panels to bottom-right panels, mutations were located on Ch.2, position 6294005 and Ch.6 position 148046062 for MBS, and on Ch.3 position 141724144 and Ch.6 position 130321359 for F252.

**Figure S15.**
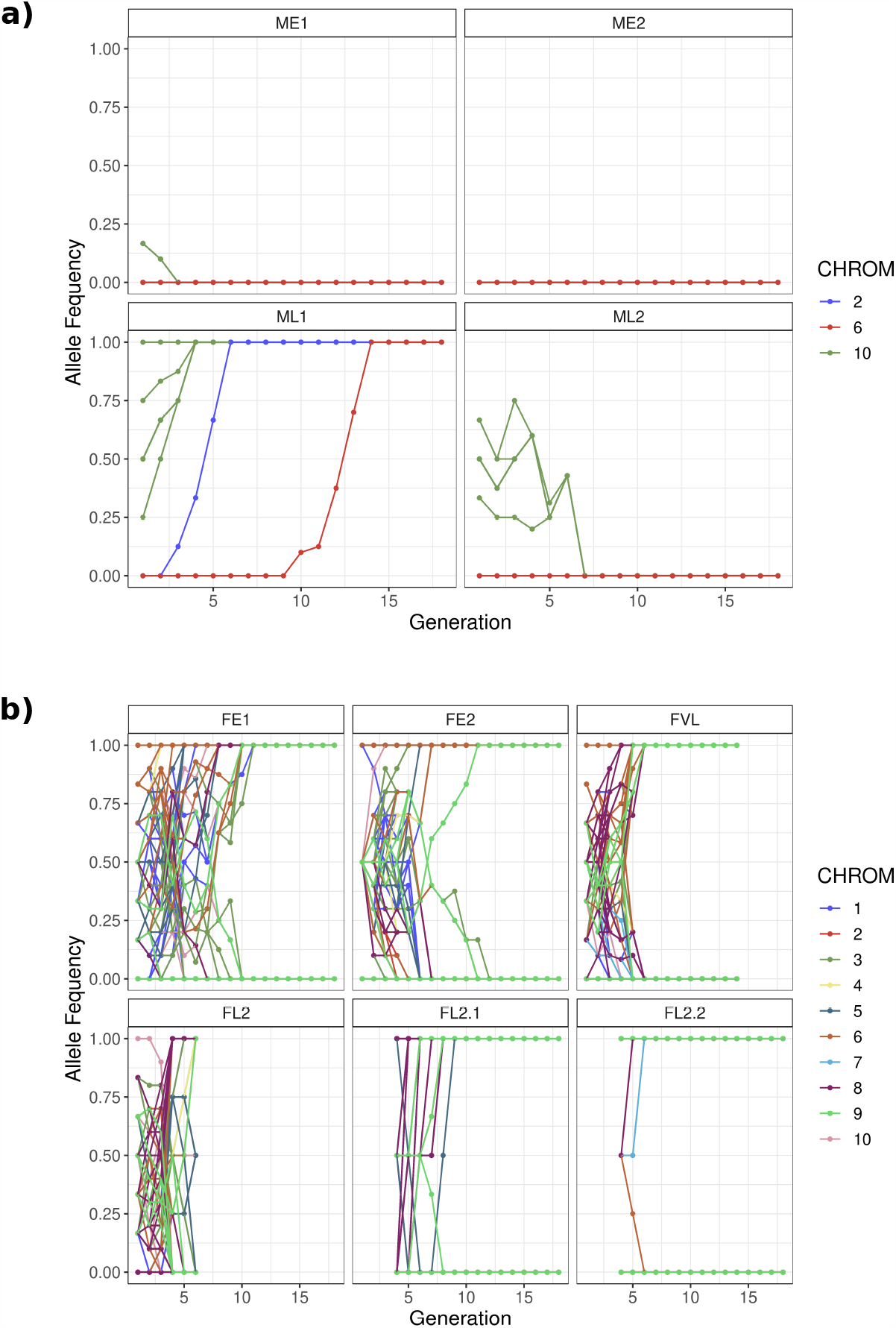
Within-family allele frequency dynamics through generations in MBS a) and F252 b). Evolution of minor allele frequency is presented. SNPs are colored according to chromosomes. Observed noise around allele frequency of 0 or 1 is caused by imperfect inferences of the algorithm. Note that three SNPs in FE2, FL2.1 and FL2.2 respectively, displayed intermediate frequencies. The original KASPar fluorescence data of these particular SNPs were difficult to interpret.

**Figure S16.**
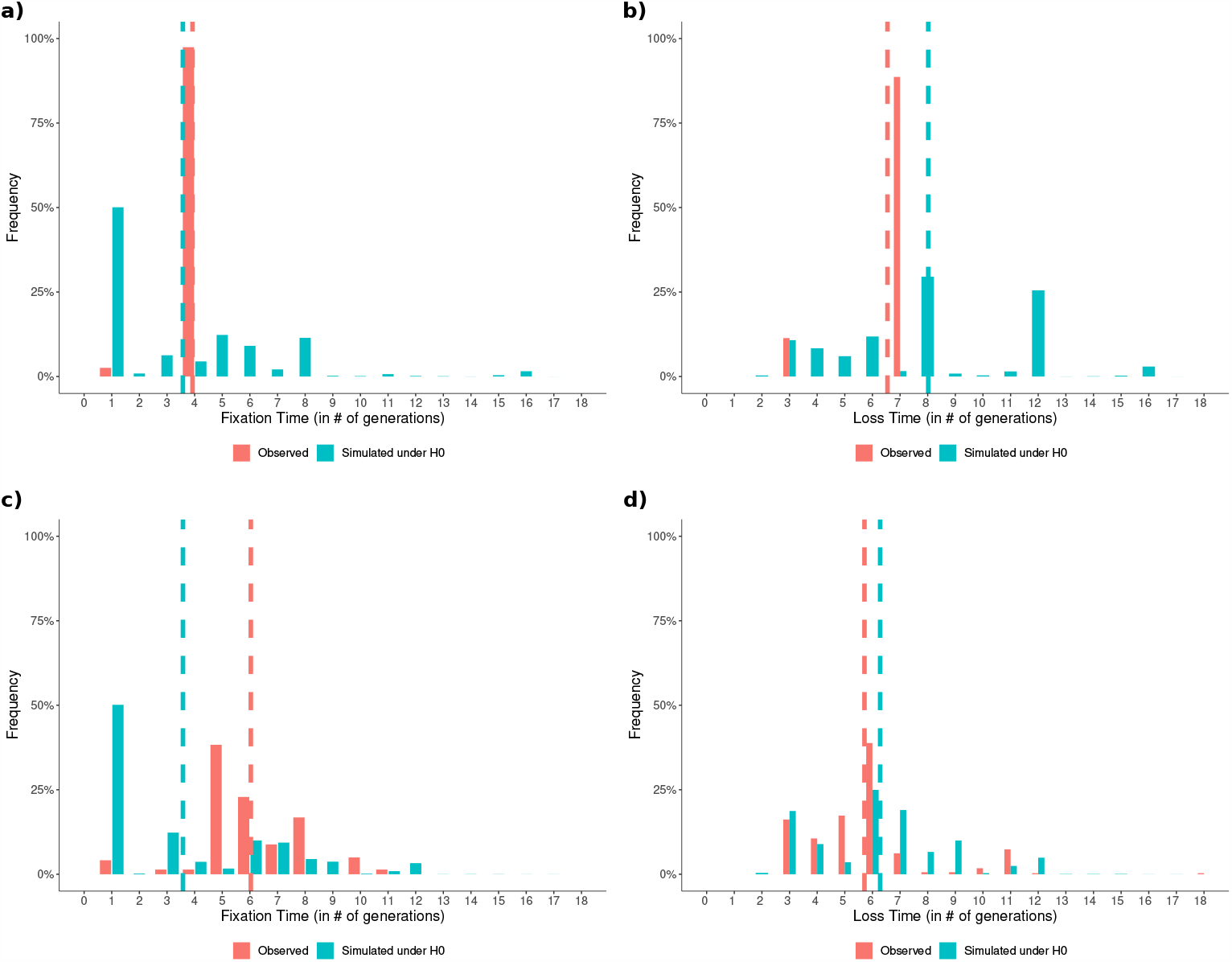
Distribution of standing variants fixation times (left panels) and time-to-loss (right panels) in MBS (upper row) and F252 (lower row) from observed (red) and simulated allele trajectories (blue). Dashed vertical lines highlight average fixation (resp. loss) times.

**Figure S17.**
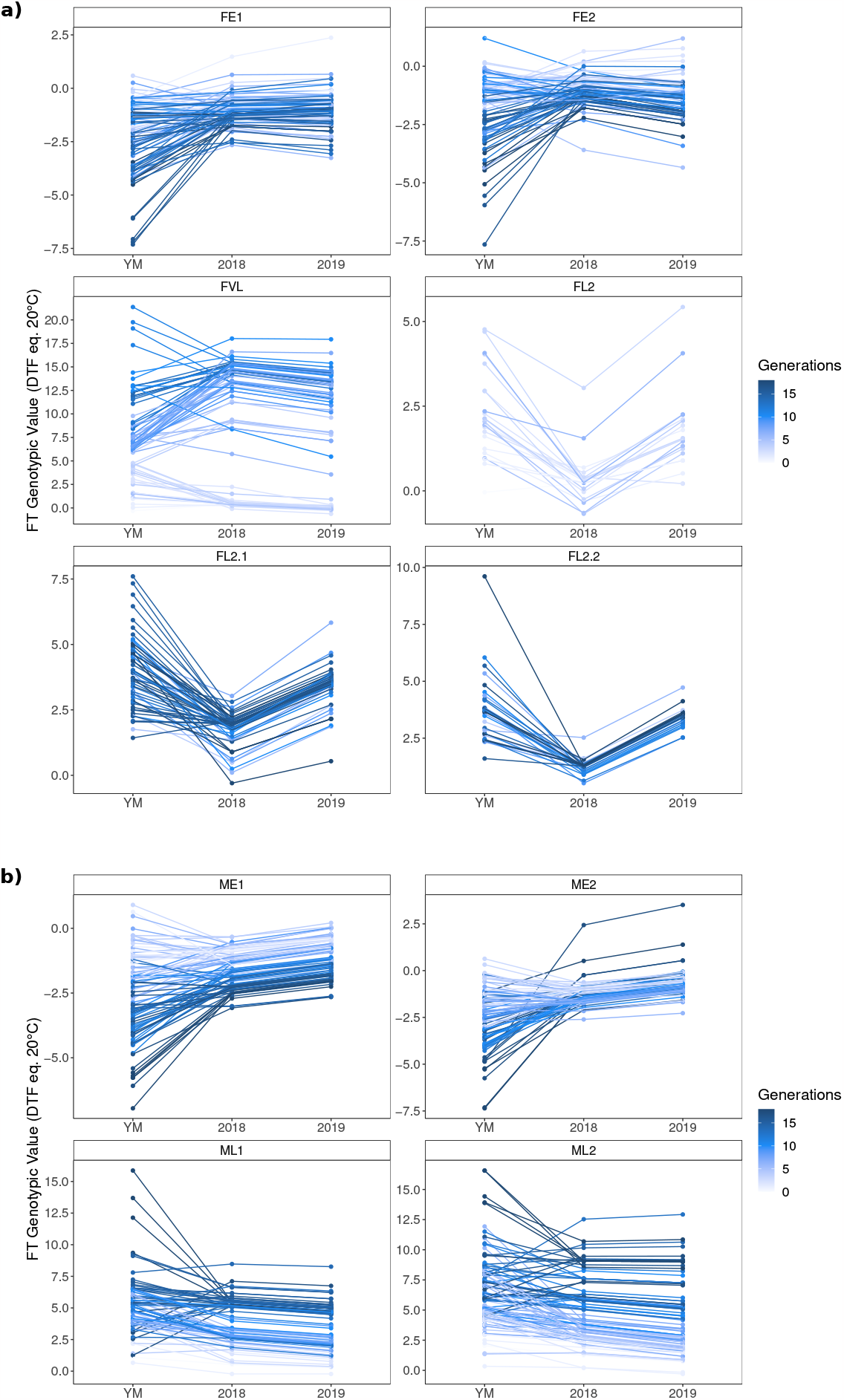
**Flowering time reaction norms per family from predicted genetic values in each environment type** in a) F252 and b) MBS. Each dot per evaluation year represents a progenitor predicted genotypic value. Straight lines link progenitor values from yearly measurement, 2018 and 2019 on a given progenitor.

**Figure S18.**
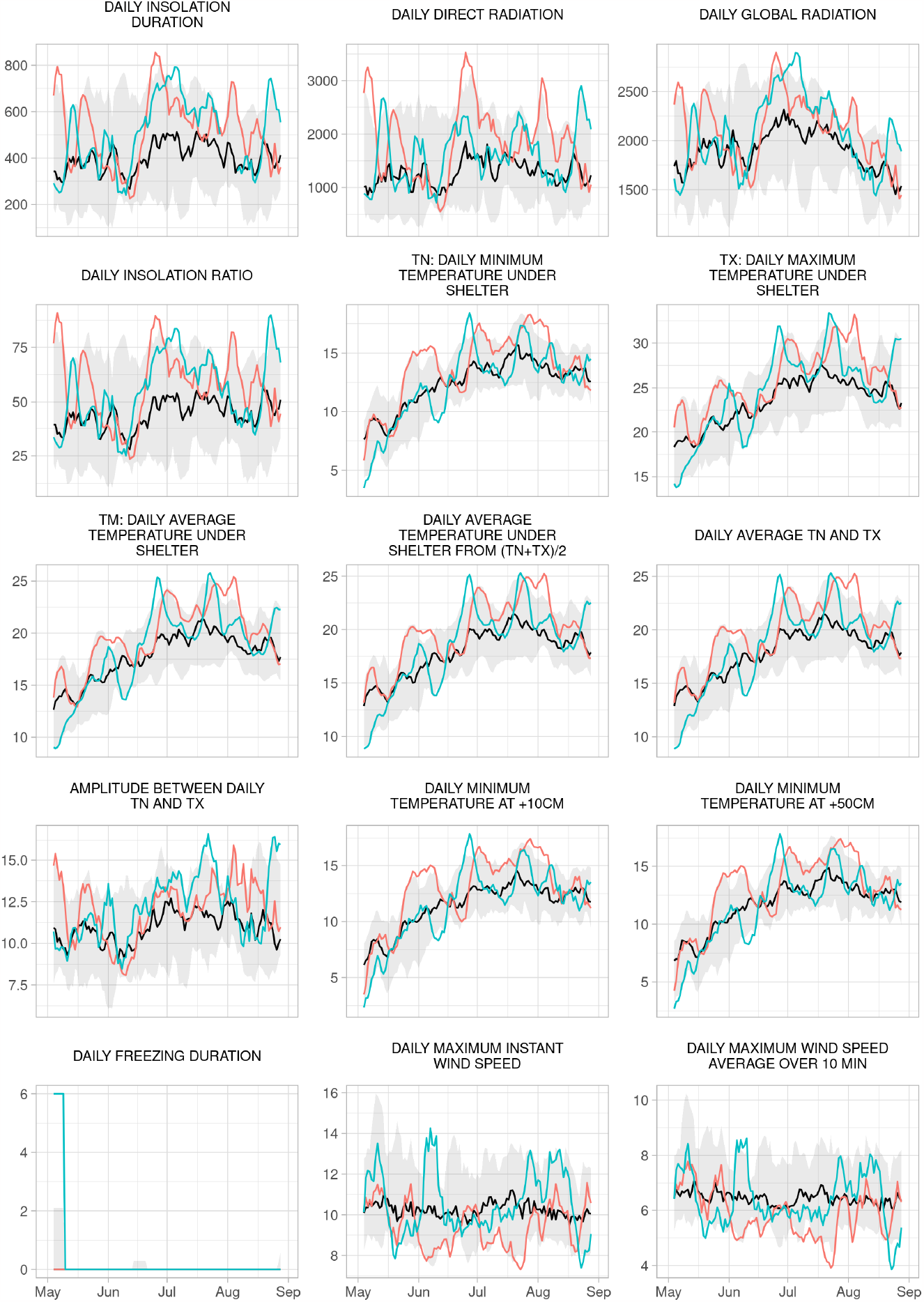
Evolution over the growing season of environmental variables recorded between 2006 and 2019 averaged over a 7-days sliding window. Red (blue) curves correspond to measured values in 2018 (2019). Black curves indicate the evolution along time of the median values over the recorded time period and grey areas include values between 5th and 95th percentiles.

**Figure S18.**
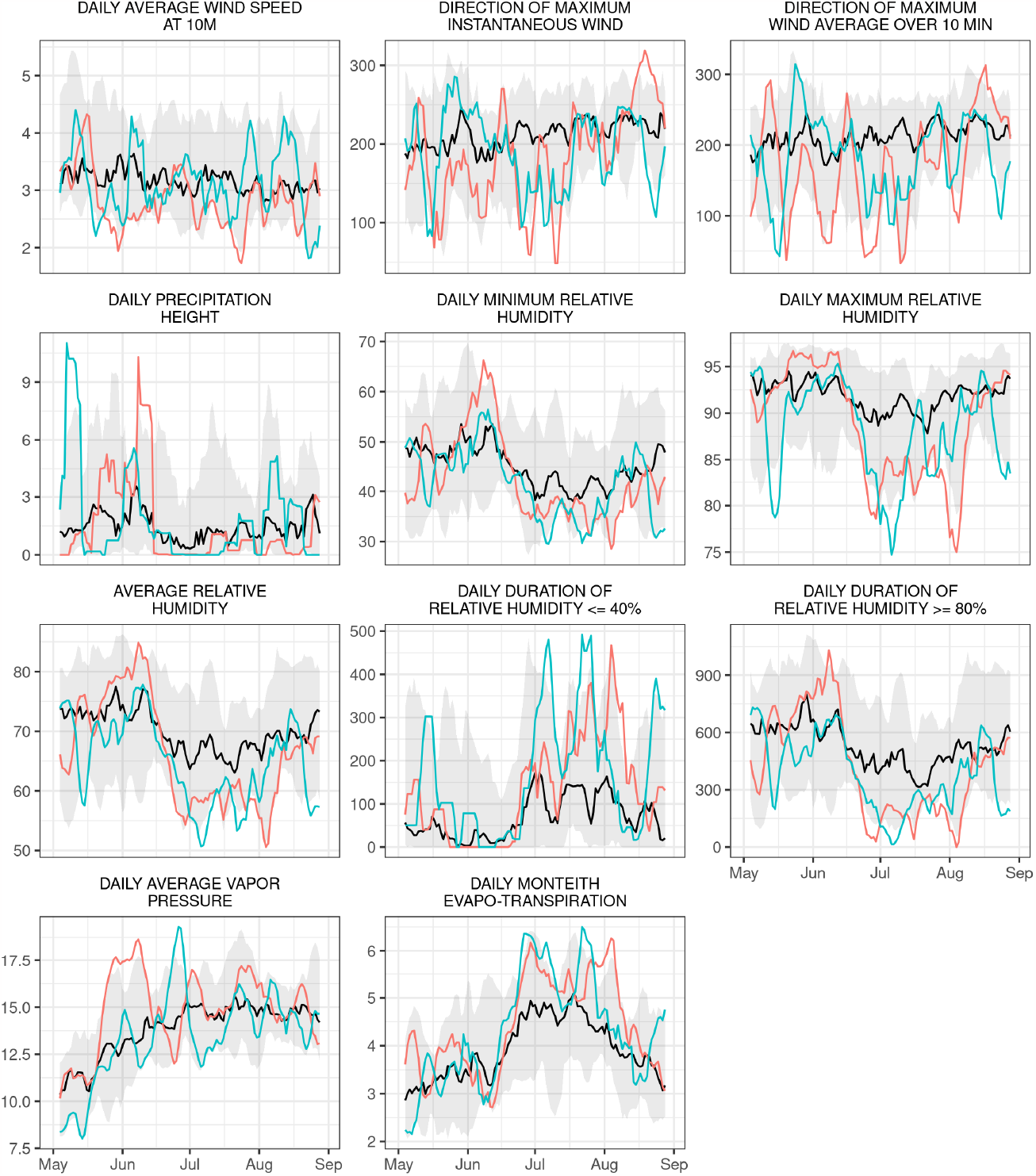
Evolution over the growing season of environmental variables recorded between 2006 and 2019 averaged over a 7-days sliding window. Red (blue) curves correspond to measured values in 2018 (2019). Black curves indicate the evolution along time of the median values over the recorded time period and grey areas include values between 5th and 95th percentiles.

**Figure S19.**
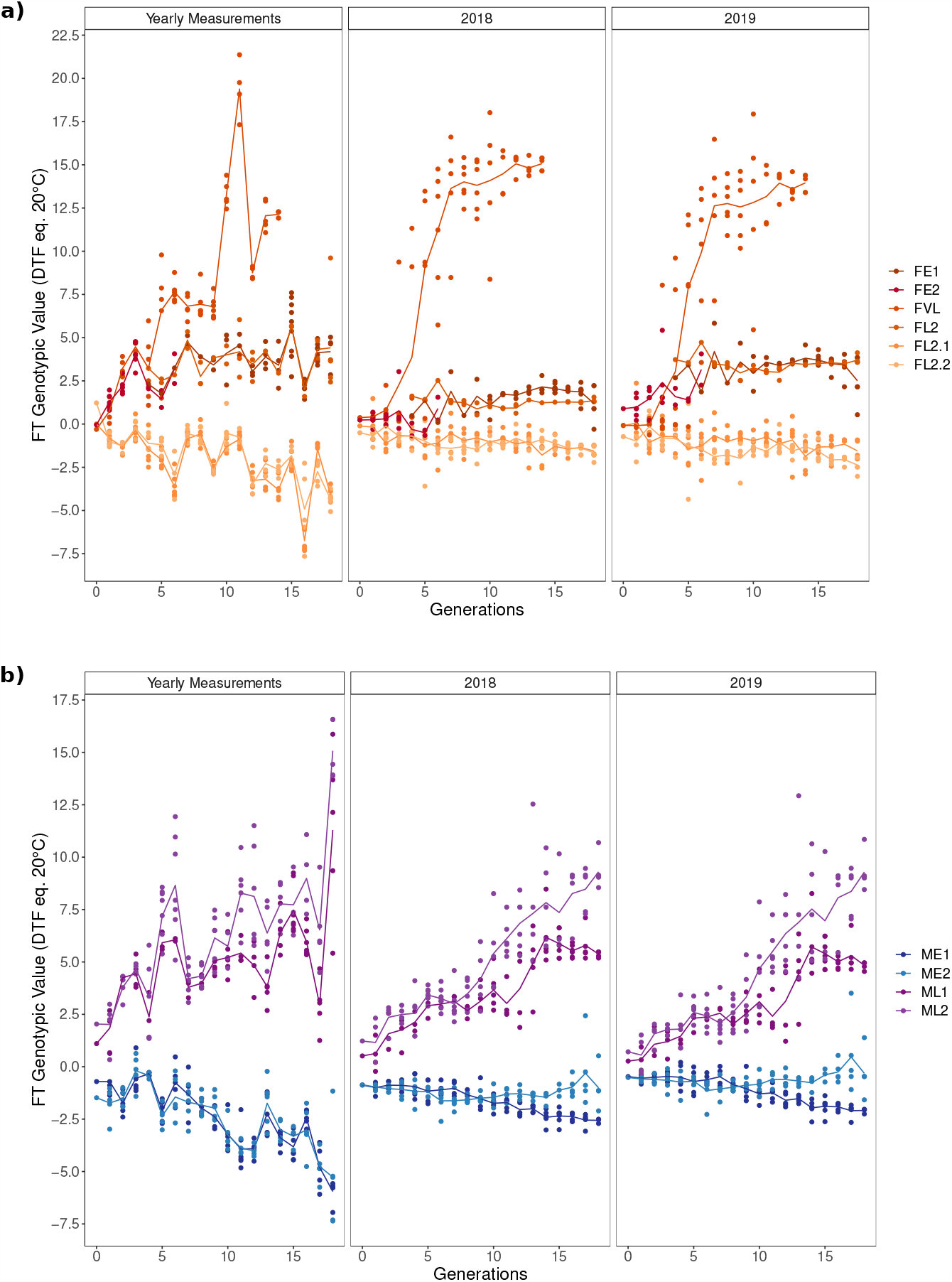
**Comparison between selection response for flowering time in each environment of evaluation** for a) F252 and b) MBS. Left panels corresponds to total genetic values predicted from yearly measurements while middle and right panels corresponds to predicted breeding values from 2018 and 2019 common garden experiments. Colors indicate the different families. Colored lines indicate the evolution of the mean value per generation and per family through time. Each dot corresponds to a progenitor.

**Figure S20.**
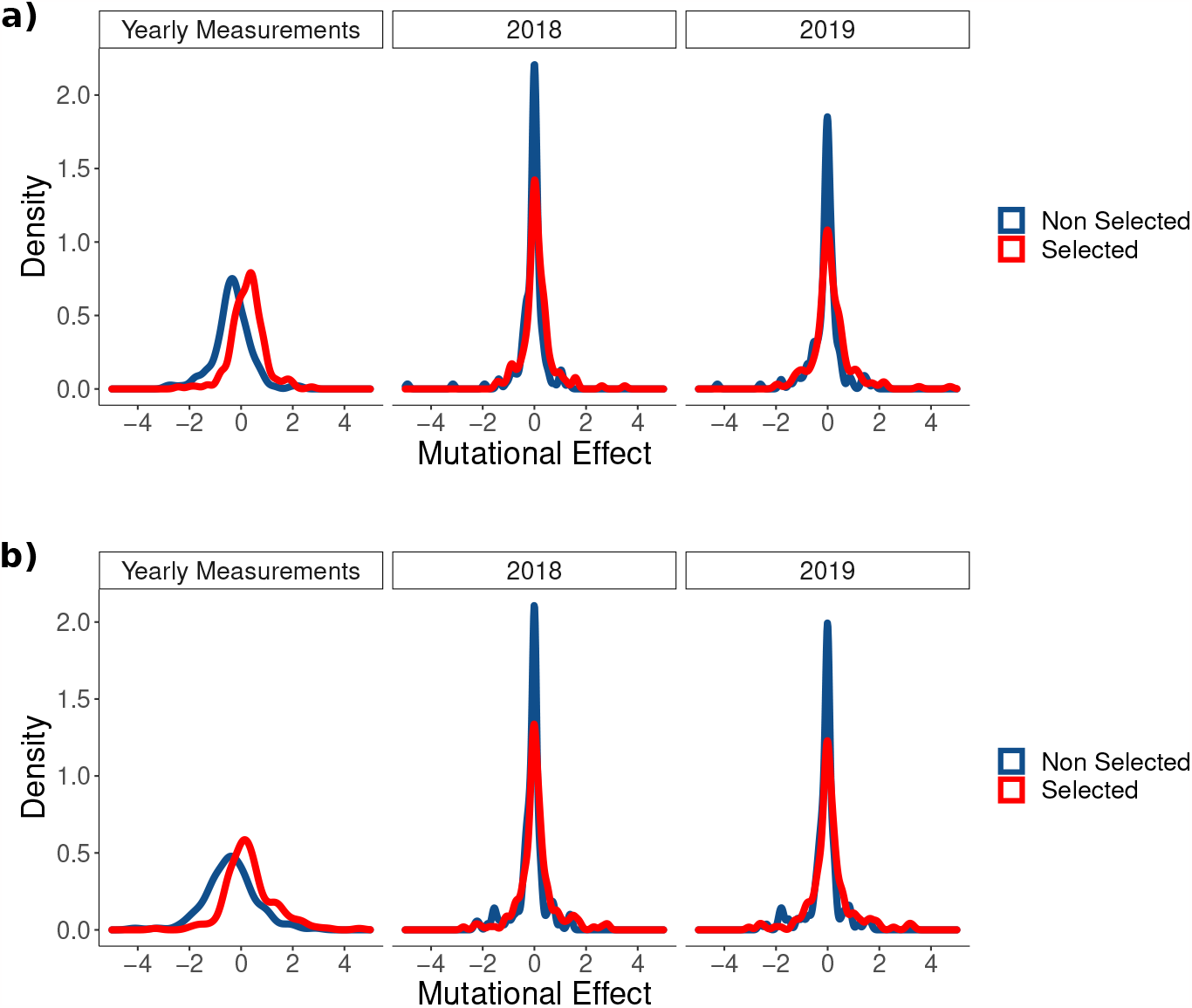
**Comparison between estimated distribution of selected (in blue) and non selected (in red) mutational effects in each environment of evaluation** for a) F252 and b) MBS. Positive values correspond to mutational effects of the same sign as the selection direction.

**Table S5.**
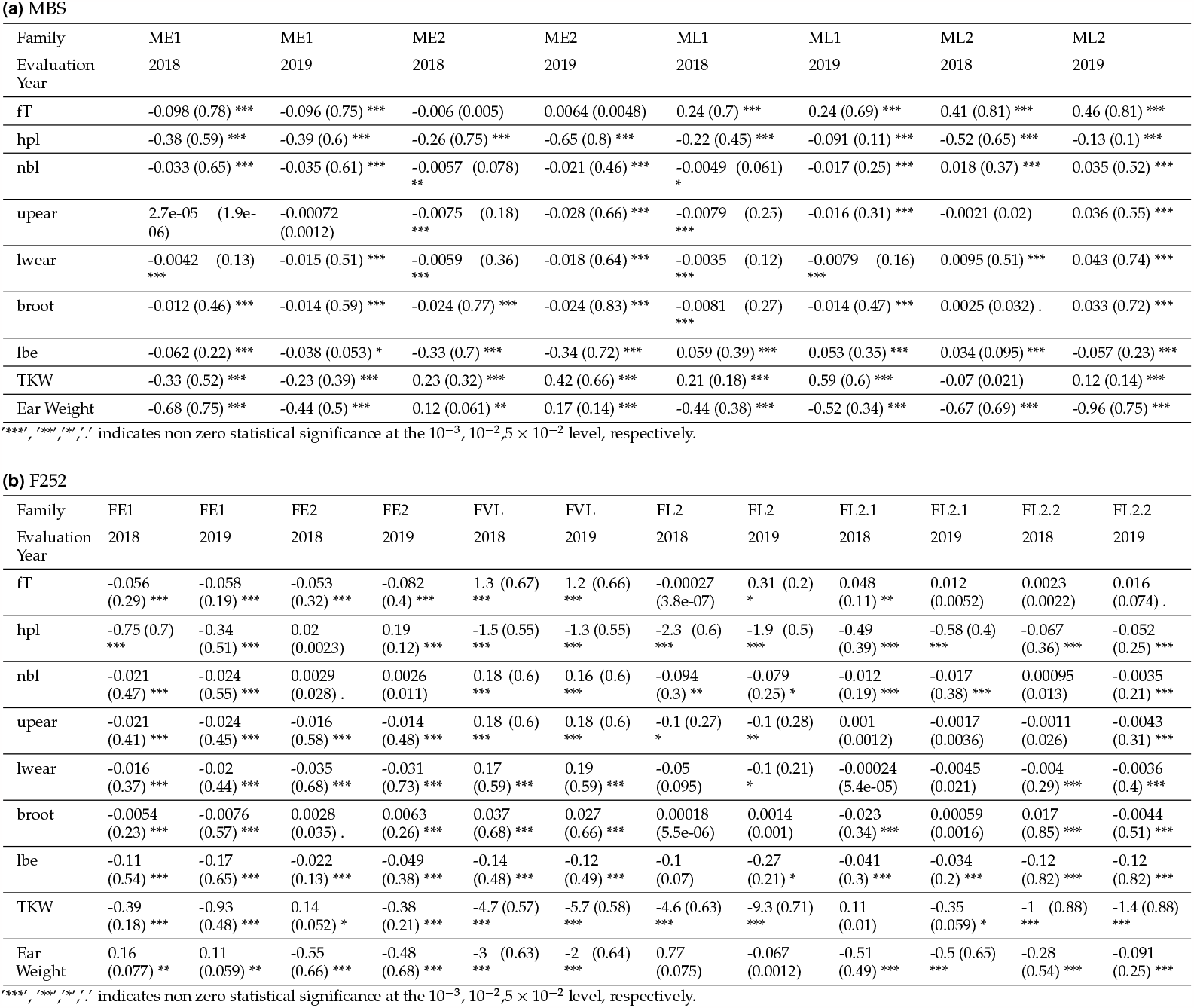
Average selection response estimated through linear regression of genetic values over generations for DSECG (2018 and 2109) in MBS (a), and F252 (b) genetic backgrounds. Slope, (R2) and significance of the regression are indicated for all measured traits.

**Table S6.** Variance (standard error) decomposition in DSECG for each traits for MBS genetic background (a), and F252 (b).

|  | FT | HT | #L | PHY | UE | LE | BR | LL | TKW | EW |
| --- | --- | --- | --- | --- | --- | --- | --- | --- | --- | --- |
| $\sigma_{02018}^2$ | 1.03 (0.465) * | 0 (0.684) | 8.56E-02<br>(3.65E-02) ** | 4.93E-04<br>(4.48E-04) | 1.64E-02<br>(1.00E-02) . | 1.24E-02<br>(7.51E-03) . | 5.08E-03<br>(8.07E-03) | 0.192 (0.239) | 16.3 (9.61) * | 0 (0.672) |
| $\sigma_{02018,2019}^2$ | 0.543 (0.283) * | 2.24 (1.41) . | 0.112 (4.63E-02) ** | -1.02E-03<br>(3.59E-04) | 3.25E-02<br>(1.75E-02) * | 2.33E-02<br>(1.14E-02) * | 4.76E-03<br>(5.76E-03) | 0.256 (0.249) | 6.2 (3.99) . | -0.188 (0.602) |
| $\sigma_{02019}^2$ | 0.206 (0.176) | 7.2 (3.44) * | 0.147 (6.27E-02) ** | 2.10E-04<br>(5.23E-04) | 5.56E-02<br>(3.14E-02) * | 3.42E-02<br>(1.87E-02) * | 0 (4.66E-03) | 0.187 (0.27) | 0 (1.86) | 0 (0.782) |
| $\sigma_{m2018}^2$ | 0.112 (1.42E-02) *** | 0.454 (9.58E-02) *** | 2.66E-03<br>(5.17E-04) *** | 6.43E-05<br>(3.70E-05) * | 1.48E-03<br>(3.69E-04) *** | 1.32E-03<br>(3.61E-04) *** | 2.35E-03<br>(5.98E-04) *** | 0.132 (2.07E-02) *** | 1.1 (0.384) ** | 0.87 (9.44E-02) *** |
| $\sigma_{m2018,2019}^2$ | 0.122 (1.36E-02) *** | 0.369 (7.00E-02) *** | 3.09E-03<br>(5.60E-04) *** | 1.55E-04<br>(4.44E-05) *** | 1.59E-03<br>(4.03E-04) *** | 1.54E-03<br>(3.69E-04) *** | 1.77E-03<br>(4.13E-04) *** | 0.114 (1.67E-02) *** | 0.595 (0.225) ** | 0.898 (7.33E-02) *** |
| $\sigma_{m2019}^2$ | 0.14 (1.62E-02) *** | 0.499 (9.82E-02) *** | 4.70E-03<br>(9.08E-04) *** | 8.34E-05<br>(7.59E-05) | 3.80E-03<br>(7.90E-04) *** | 4.11E-03<br>(7.00E-04) *** | 2.88E-03<br>(5.40E-04) *** | 0.134 (1.98E-02) *** | 1.16 (0.27) *** | 1.24 (0.113) *** |
| $\sigma_{2018}^2$ | 0.272 (7.61E-02) *** | 8.5 (1.7) *** | 4.25E-02<br>(4.46E-03) *** | 1.01E-02<br>(1.18E-03) *** | 5.57E-02<br>(5.51E-03) *** | 3.12E-02<br>(7.65E-03) *** | 3.32E-02<br>(1.03E-02) *** | 1.5 (0.246) *** | 53.7 (6.72) *** | 0 (3.74) |
| $\sigma_{2019}^2$ | 0.795 (0.109) *** | 6.22 (1.58) *** | 8.25E-02<br>(8.07E-03) *** | 5.24E-02<br>(4.11E-03) *** | 8.93E-02<br>(8.55E-03) *** | 6.12E-02<br>(9.74E-03) *** | 0 (7.16E-03) | 1.46 (0.237) *** | 42.5 (5.59) *** | 0 (3.78) |
| Intercept (2018) | 78.2 (0.691) *** | 1.41E+02<br>(1.15) *** | 18.3 (0.169) *** | 3.25 (3.63E-02) *** | 12.7 (8.46E-02) *** | 11.6 (9.85E-02) *** | 7.7 (7.52E-02) *** | 69.9 (0.518) *** | 1.98E+02<br>(2.73) *** | 62.3 (1.11) *** |
| 2019 Effect | 7.64 (0.494) *** | -25.8 (1.75) | -0.652 (8.86E-02) | -6.72E-02<br>(5.25E-02) | -0.123 (7.64E-02) | -0.28 (0.102) | 1.07 (6.31E-02) *** | -5.65 (0.446) | 17.7 (1.88) *** | 3.15 (1.3) ** |
| $h_{m2018}^2 = \frac{\sigma_{m2018}^2}{\sigma_{2018}^2}$ | 0.411 (0.133) ** | 5.34E-02<br>(1.79E-02) ** | 6.27E-02<br>(1.51E-02) *** | 6.39E-03<br>(4.00E-03) . | 2.65E-02<br>(7.80E-03) *** | 4.22E-02<br>(1.79E-02) ** | 7.08E-02<br>(3.29E-02) * | 8.78E-02<br>(2.19E-02) *** | 2.05E-02<br>(8.47E-03) ** | NA (NA) |
| $h_{m2019}^2 = \frac{\sigma_{m2019}^2}{\sigma_{2019}^2}$ | 0.176 (3.31E-02) *** | 8.02E-02<br>(2.98E-02) ** | 5.69E-02<br>(1.33E-02) *** | 1.59E-03<br>(1.48E-03) | 4.26E-02<br>(1.06E-02) *** | 6.71E-02<br>(1.78E-02) *** | NA (NA) | 9.21E-02<br>(2.23E-02) *** | 2.73E-02<br>(8.21E-03) *** | NA (NA) |

\*\*\*, \*\*, \*, . indicates non zero statistical significance at the $10^{-3}$ , $10^{-2}$ , $5 \times 10^{-2}$ level, respectively.

**(b) F252**
|  | FT | HT | #L | PHY | UE | LE | BR | LL | TKW | EW |
| --- | --- | --- | --- | --- | --- | --- | --- | --- | --- | --- |
| $\sigma_{02018}^2$ | 12.9 (3.93) *** | 69 (23.4) ** | 0.345 (0.115) ** | 8.52E-04<br>(1.00E-03) | 0.376 (0.119) *** | 0.293 (9.87E-02) ** | 1.40E-02<br>(1.21E-02) | 0.723 (0.364) * | 2.75E+02<br>(98.2) ** | 2.56E+02<br>(70.5) *** |
| $\sigma_{02018,2019}^2$ | 11.5 (3.81) ** | 59.4 (22.6) ** | 0.311 (0.102) ** | 1.87E-03<br>(1.72E-03) | 0.375 (0.117) *** | 0.349 (0.112) *** | 2.62E-03<br>(5.52E-03) | 1.64 (0.637) ** | 3.56E+02<br>(1.22E+02) ** | 1.72E+02<br>(47.6) *** |
| $\sigma_{02019}^2$ | 11.7 (4.1) ** | 78.2 (28.6) ** | 0.283 (9.59E-02) ** | 4.30E-03<br>(4.03E-03) | 0.377 (0.119) *** | 0.421 (0.138) ** | 1.88E-03<br>(3.13E-03) | 3.19 (1.22) ** | 4.62E+02<br>(1.65E+02) ** | 1.14E+02<br>(33.3) *** |
| $\sigma_{m2018}^2$ | 8.74E-02<br>(1.62E-02) *** | 0.512 (0.146) *** | 3.43E-03<br>(6.72E-04) *** | 1.02E-04<br>(6.36E-05) . | 2.87E-03<br>(6.36E-04) *** | 2.96E-03<br>(7.42E-04) *** | 1.13E-03<br>(5.06E-04) * | 8.32E-02<br>(1.72E-02) *** | 3.34 (0.797) *** | 0.781 (0.211) *** |
| $\sigma_{m2018,2019}^2$ | 0.118 (1.99E-02) *** | 0.453 (0.128) *** | 2.46E-03<br>(5.41E-04) *** | 8.93E-05<br>(7.20E-05) | 2.42E-03<br>(5.10E-04) *** | 1.69E-03<br>(5.79E-04) ** | 1.07E-03<br>(2.66E-04) *** | 5.83E-02<br>(1.26E-02) *** | 2.65 (0.821) *** | 0.385 (0.169) * |
| $\sigma_{m2019}^2$ | 0.144 (3.09E-02) *** | 0.61 (0.197) *** | 2.60E-03<br>(6.74E-04) *** | 1.17E-04<br>(1.64E-04) | 2.48E-03<br>(6.16E-04) *** | 2.35E-03<br>(8.17E-04) ** | 6.24E-04<br>(1.80E-04) *** | 6.94E-02<br>(1.69E-02) *** | 5.26 (1.49) *** | 0.741 (0.242) ** |
| $\sigma_{2018}^2$ | 0.781 (8.83E-02) *** | 14.5 (1.67) *** | 4.51E-02<br>(4.96E-03) *** | 1.55E-02<br>(1.58E-03) *** | 5.46E-02<br>(5.87E-03) *** | 7.87E-02<br>(8.52E-03) *** | 0.172 (1.51E-02) *** | 1.43 (0.308) *** | 80.1 (8.7) *** | 19 (2.1) *** |
| $\sigma_{2019}^2$ | 2.99 (0.303) *** | 22.9 (2.45) *** | 7.06E-02<br>(7.24E-03) *** | 5.69E-02<br>(5.42E-03) *** | 5.69E-02<br>(5.90E-03) *** | 0.105 (1.09E-02) *** | 3.68E-02<br>(3.70E-03) *** | 1.06 (0.285) *** | 1.71E+02<br>(17.6) *** | 34 (3.5) *** |
| Intercept (2018) | 63.1 (1.98) *** | 1.45E+02<br>(4.77) *** | 16 (0.325) *** | 2.98 (3.33E-02) *** | 11.5 (0.337) *** | 10.7 (0.301) *** | 6.65 (0.108) *** | 66.1 (0.636) *** | 2.33E+02<br>(9.24) *** | 67.6 (8.74) *** |
| 2019 Effect | 11.7 (0.84) *** | -46.5 (3.69) | -1.67 (8.99E-02) | 0.882 (4.78E-02) *** | -0.749 (7.46E-02) | -1.17 (0.112) | 0.327 (0.124) ** | -11 (0.703) | 16.9 (3.98) *** | -20.7 (3.29) |
| $h_{m2018}^2 = \frac{\sigma_{m2018}^2}{\sigma_{2018}^2}$ | 0.112 (2.60E-02) *** | 3.54E-02<br>(1.21E-02) ** | 7.60E-02<br>(1.85E-02) *** | 6.61E-03<br>(4.45E-03) . | 5.26E-02<br>(1.41E-02) *** | 3.76E-02<br>(1.13E-02) *** | 6.58E-03<br>(3.15E-03) * | 5.83E-02<br>(2.02E-02) ** | 4.17E-02<br>(1.21E-02) *** | 4.11E-02<br>(1.31E-02) *** |
| $h_{m2019}^2 = \frac{\sigma_{m2019}^2}{\sigma_{2019}^2}$ | 4.80E-02<br>(1.21E-02) *** | 2.67E-02<br>(9.84E-03) ** | 3.68E-02<br>(1.10E-02) *** | 2.05E-03<br>(2.97E-03) | 4.35E-02<br>(1.26E-02) *** | 2.24E-02<br>(8.74E-03) ** | 1.70E-02<br>(5.55E-03) ** | 6.52E-02<br>(2.74E-02) ** | 3.08E-02<br>(1.01E-02) ** | 2.18E-02<br>(8.07E-03) ** |

\*\*\*, \*\*, \*, . indicates non zero statistical significance at the $10^{-3}$ , $10^{-2}$ , $5 \times 10^{-2}$ level, respectively.

**Figure S21.**
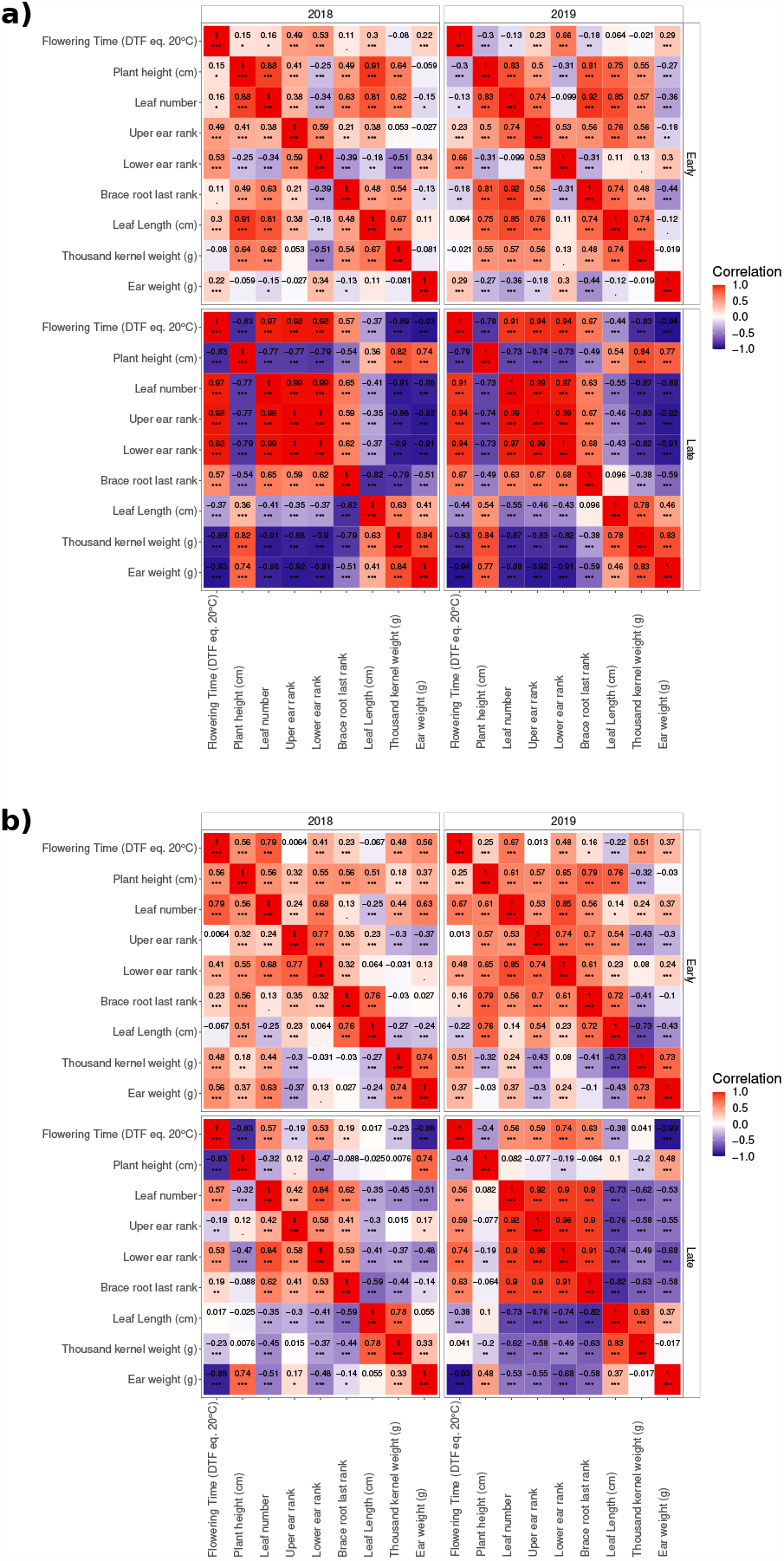
**Correlation matrix of the predicted environment specific total breeding values for all measured pairs of traits** for a) F252 and b) MBS. Pearson’s correlation coefficients were computed by environment and by population. Color corresponds the intensity of the correlations. ‘***’, ‘**’,’*’,’.’ indicates statistical significance at the 10^−3^, 10^−2^,5 *×* 10^−2^ level, respectively.

**Figure S22.**
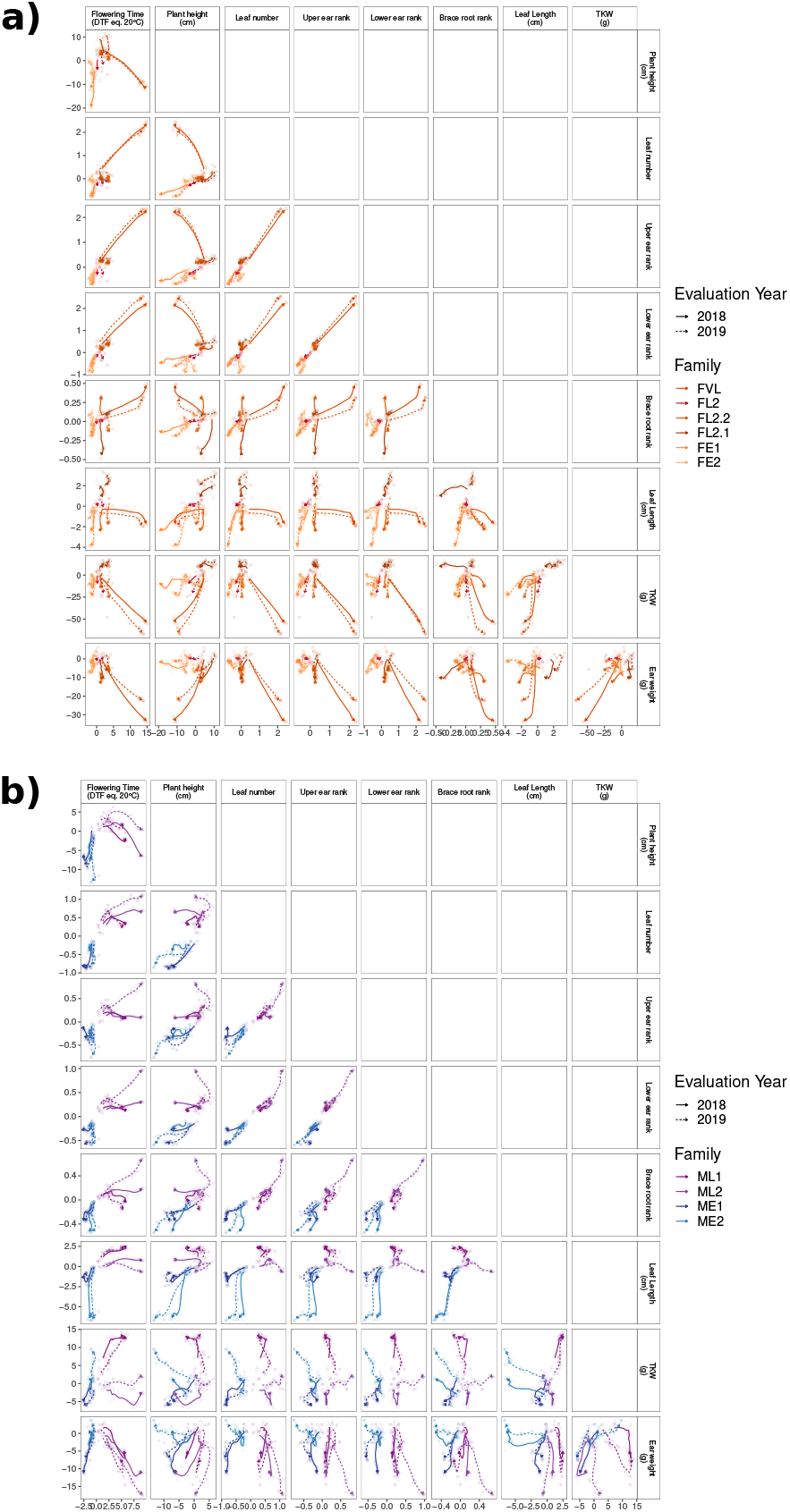
Evolution through generations of the environment specific breeding values for all pair of measured traits. for a) F252 and b) MBS. Colored lines indicate the evolution through time of the rolling mean breeding value per generation, family and evaluation year. Solid lines correspond to 2018 while dotted lines refer to 2019. Each transparent point corresponds to the average breeding value of a generation per family. Dots correspond to 2018 specific breeding values, while triangles correspond to 2019.

**Figure S23.**
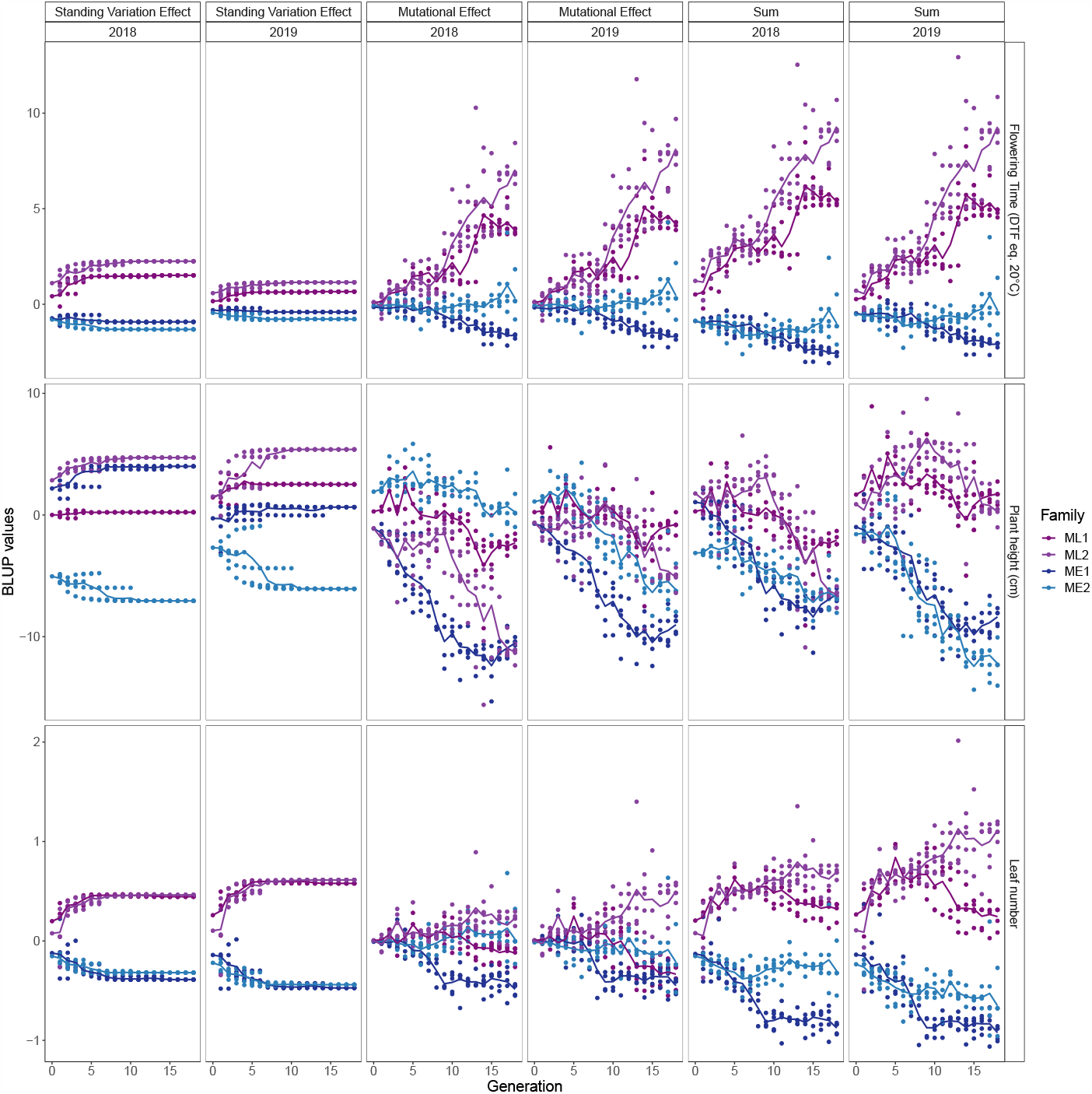
Evolution through generations of the BLUP values for each trait in the two DSECG environments (2018 and 2019) in MBS. Colored lines indicate the evolution of the mean value per generation and family through time. Each dot corresponds to a progenitor.

**Figure S23.**
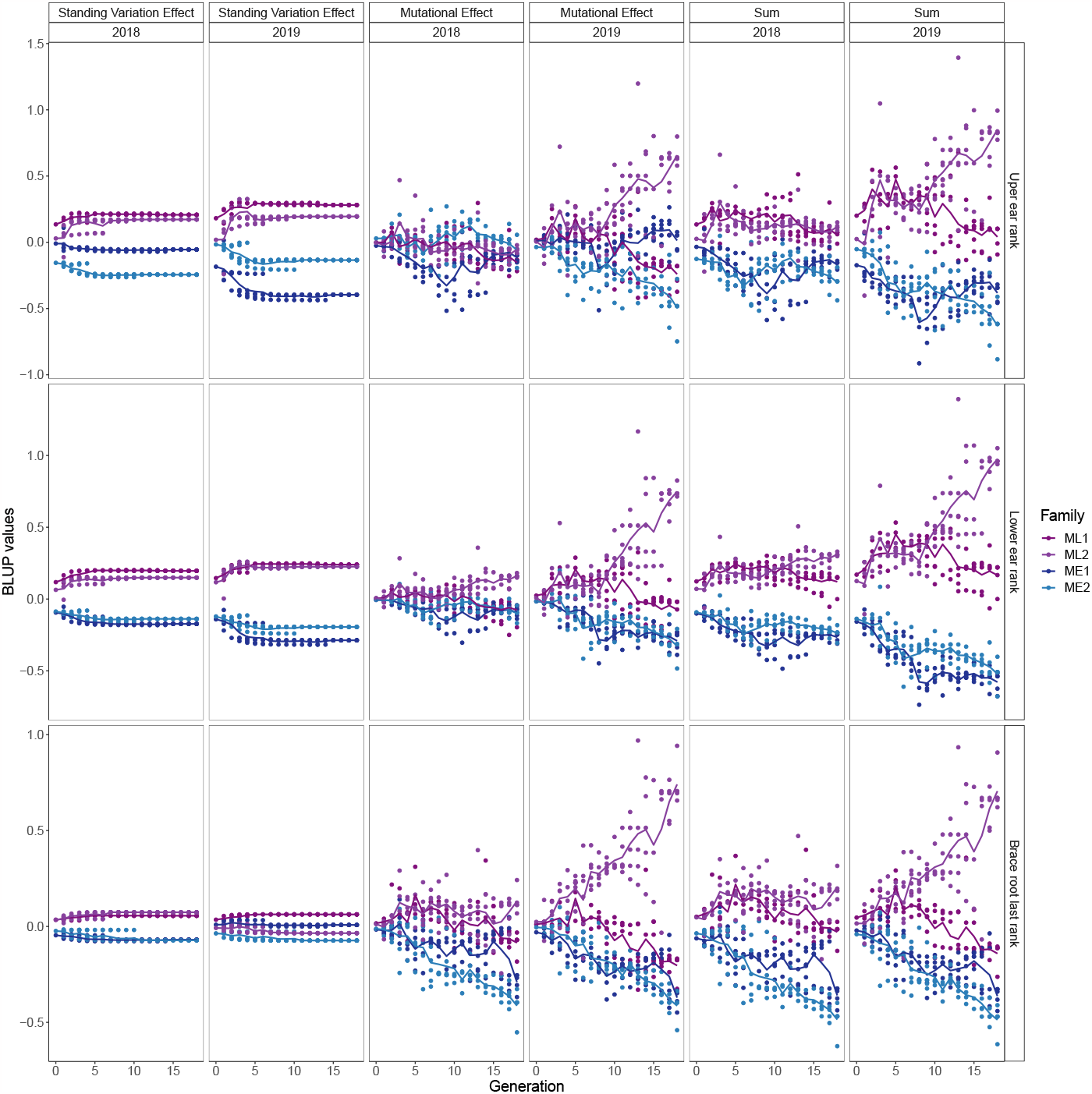
Evolution through generations of the BLUP values for each trait in the two DSECG environments (2018 and 2019) in MBS. Colored lines indicate the evolution of the mean value per generation and family through time. Each dot corresponds to a progenitor.

**Figure S23.**
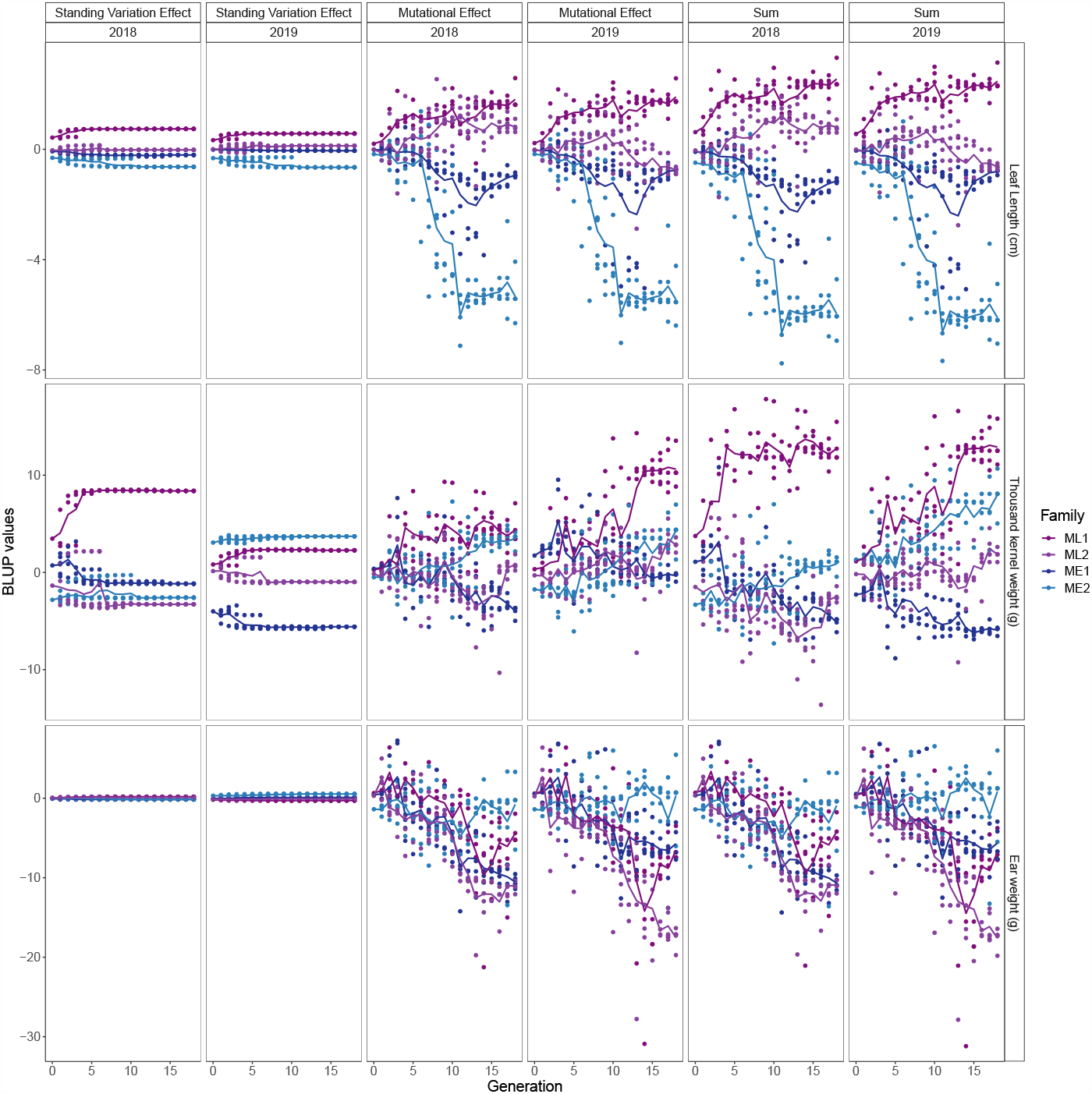
Evolution through generations of the BLUP values for each trait in the two DSECG environments (2018 and 2019) in MBS. Colored lines indicate the evolution of the mean value per generation and family through time. Each dot corresponds to a progenitor.

**Figure S24.**
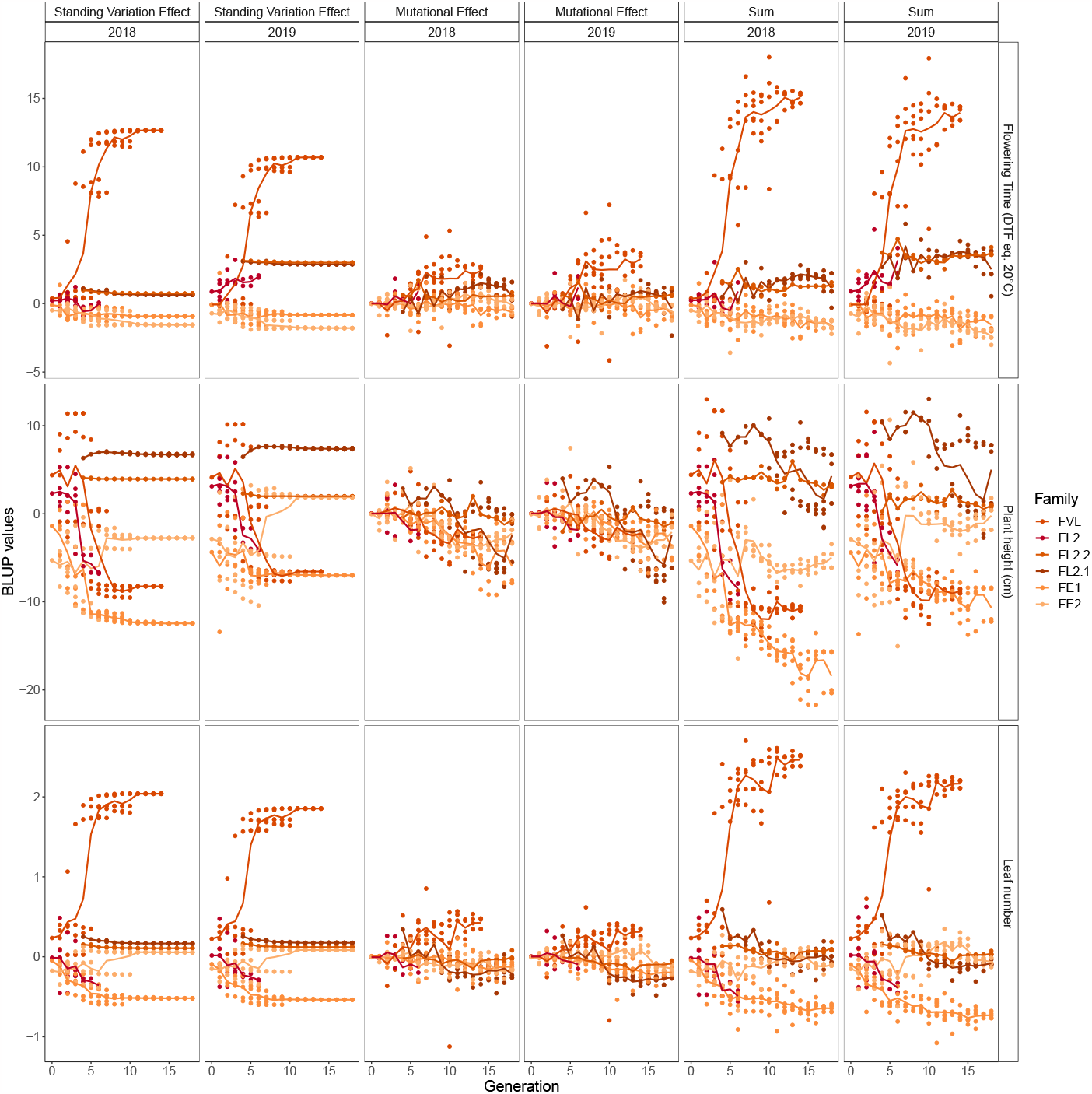
Evolution through generations of the BLUP values for each trait in the two DSECG environments (2018 and 2019) in F252. Colored lines indicate the evolution of the mean value per generation and family through time. Each dot corresponds to a progenitor.

**Figure S24.**
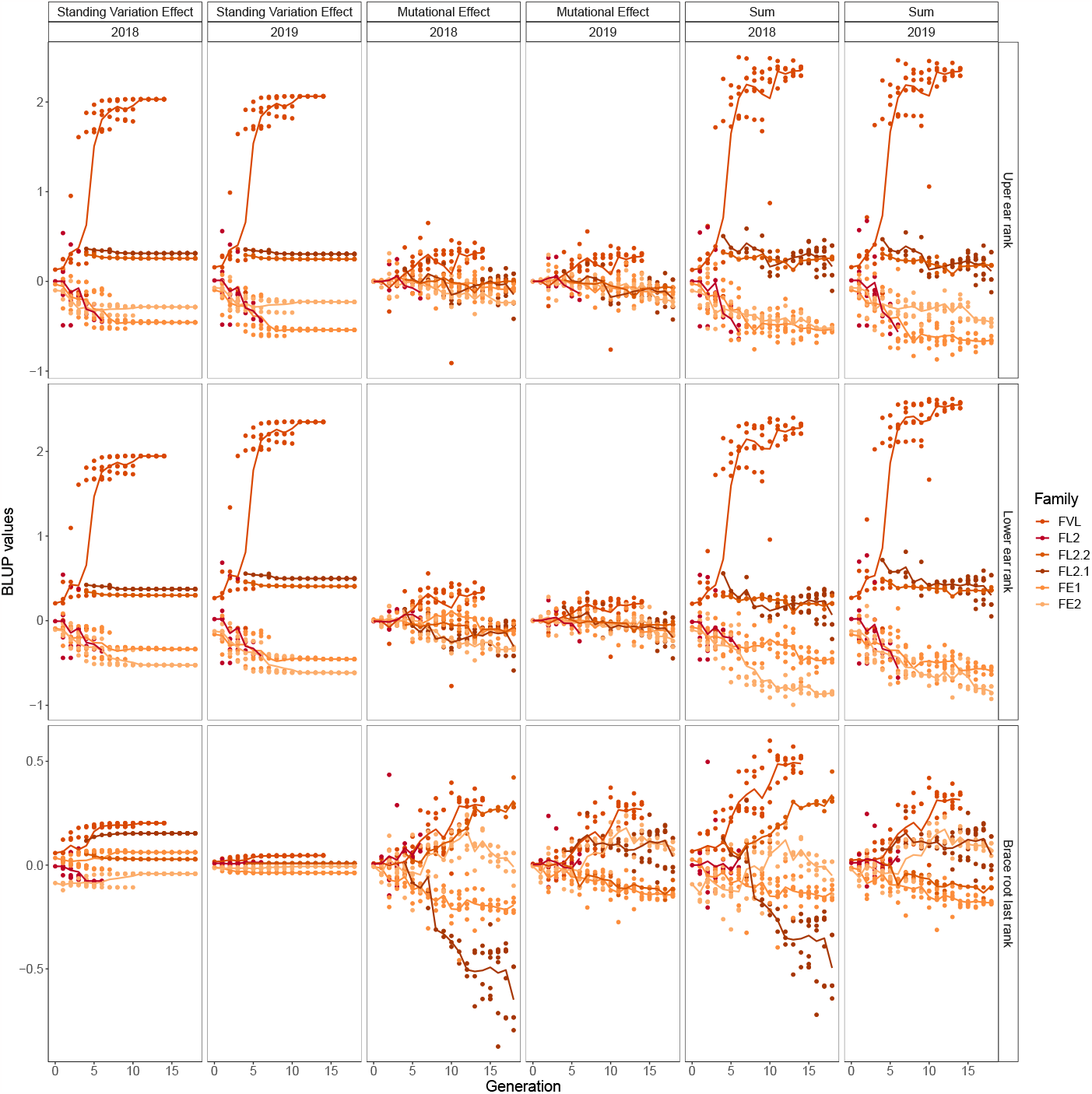
Evolution through generations of the BLUP values for each trait in the two DSECG environments (2018 and 2019) in F252. Colored lines indicate the evolution of the mean value per generation and family through time. Each dot corresponds to a progenitor.

**Figure S24.**
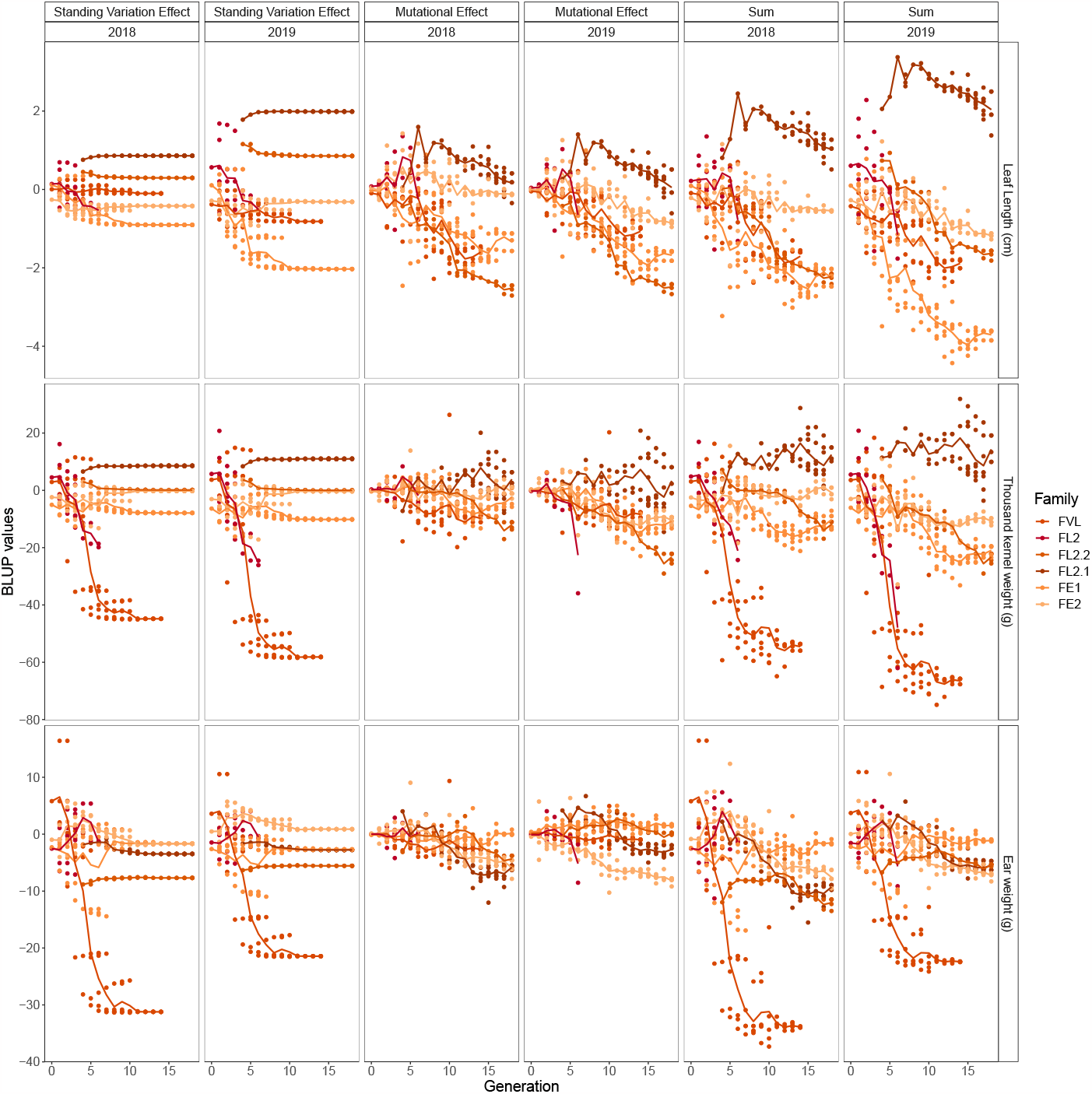
Evolution through generations of the BLUP values for each trait in the two DSECG environments (2018 and 2019) in F252. Colored lines indicate the evolution of the mean value per generation and family through time. Each dot corresponds to a progenitor.

**Figure S25.**
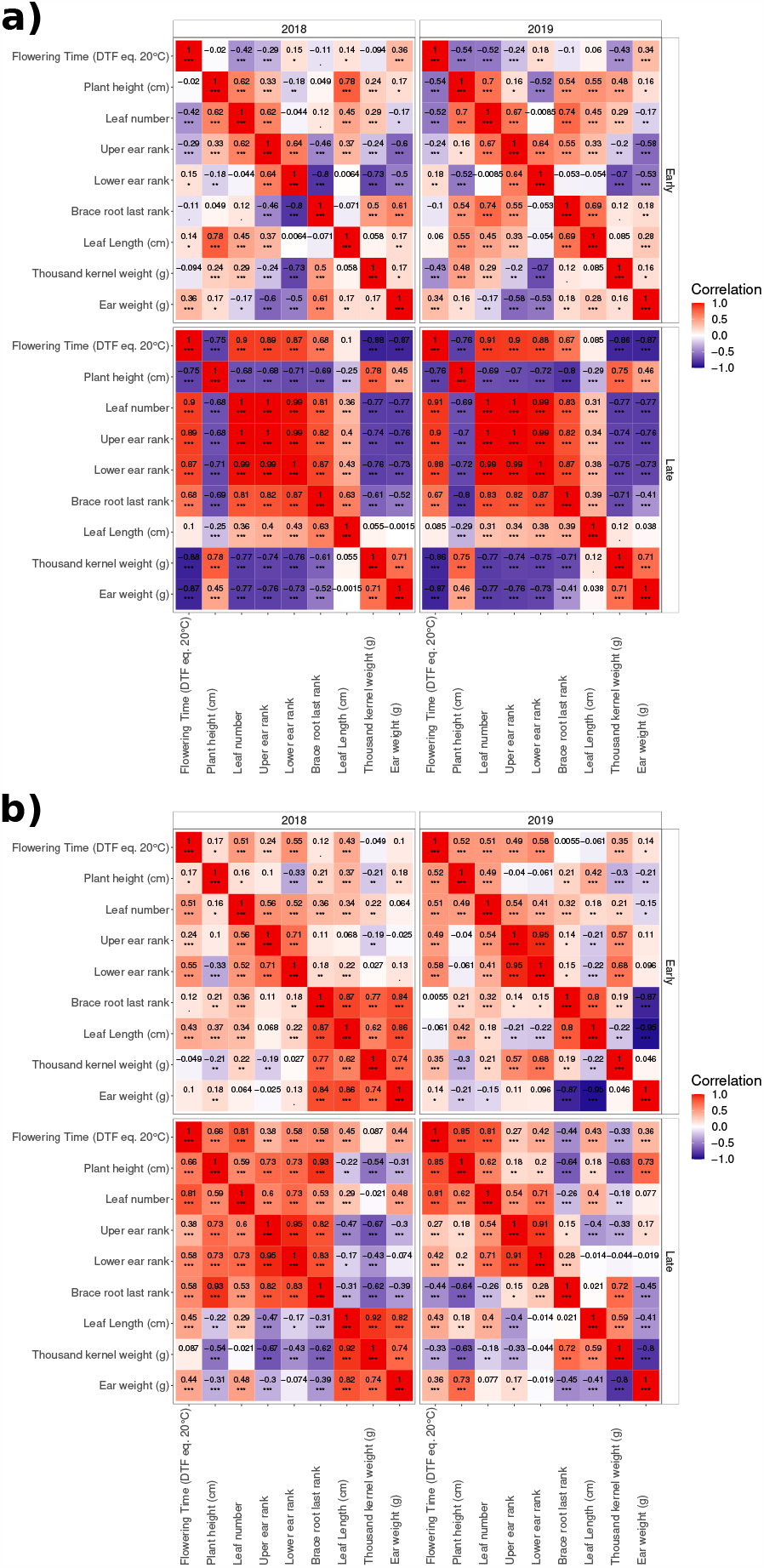
**Correlation matrix of the predicted environment specific standing variation contribution for all measured pairs of traits** for a) F252 and b) MBS. Pearson’s correlation coefficients were computed by environment and by population. Color corresponds the intensity of the correlations. ‘***’, ‘**’,’*’,’.’ indicates statistical significance at the 10^−3^, 10^−2^,5 *×* 10^−2^ level, respectively.

**Figure S26.**
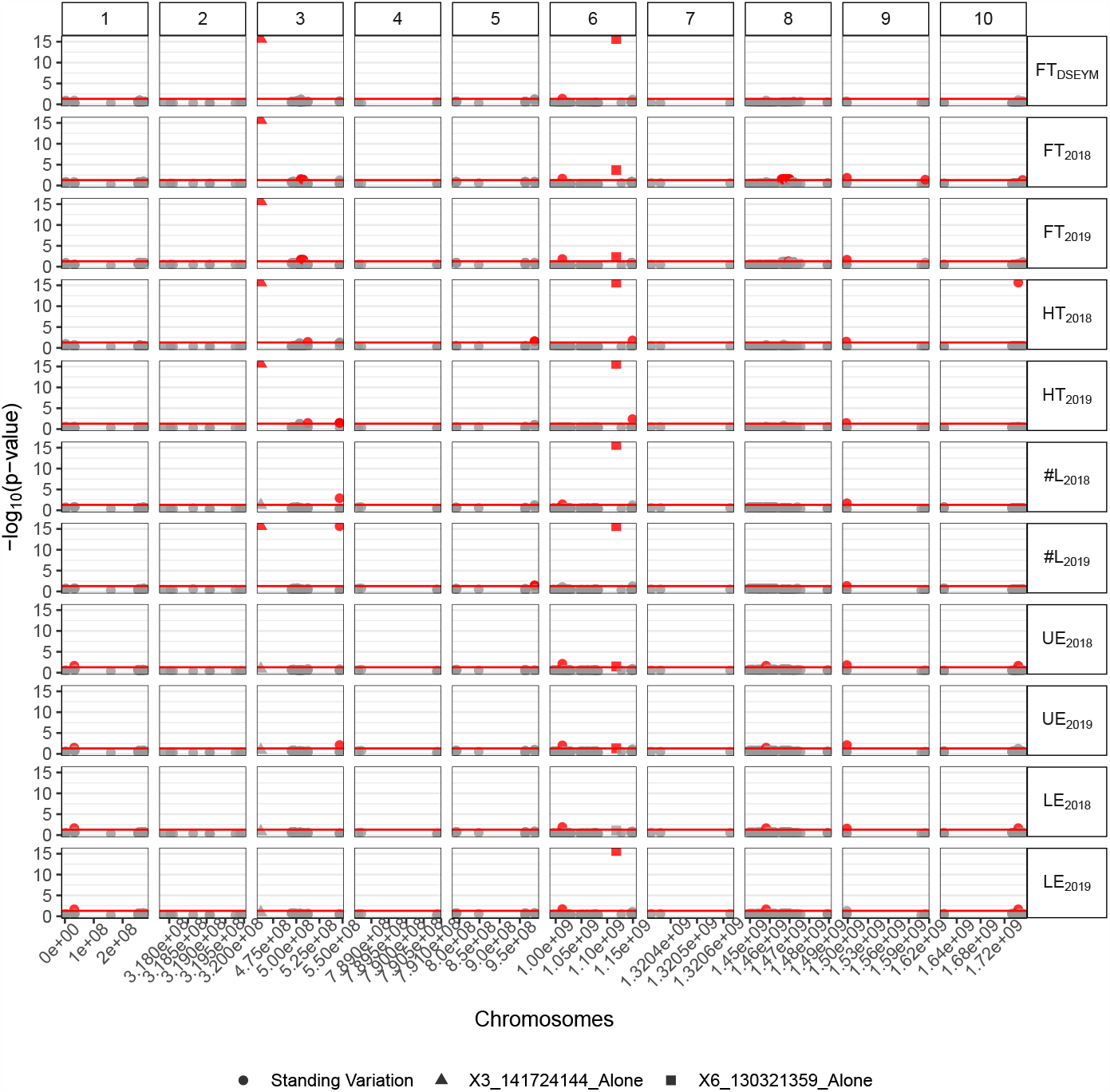
In F252, Manhattan plot for each trait representing the significance of the association of each SNPs (−*log*_10_(*p* − *value*), *FDR* < 5%). Each row of panels corresponds to one trait. Each column of panels corresponds to a chromosome. X-axis indicates the position of SNPs along the chromosomes in bp. Red point (resp. gray point) corresponds to significant (resp. non-significant) association to the trait for the corresponding SNP. Standing variation are represented by a dot, while the two detected mutations are represented respectively by a triangle or a square.

**Figure S26.**
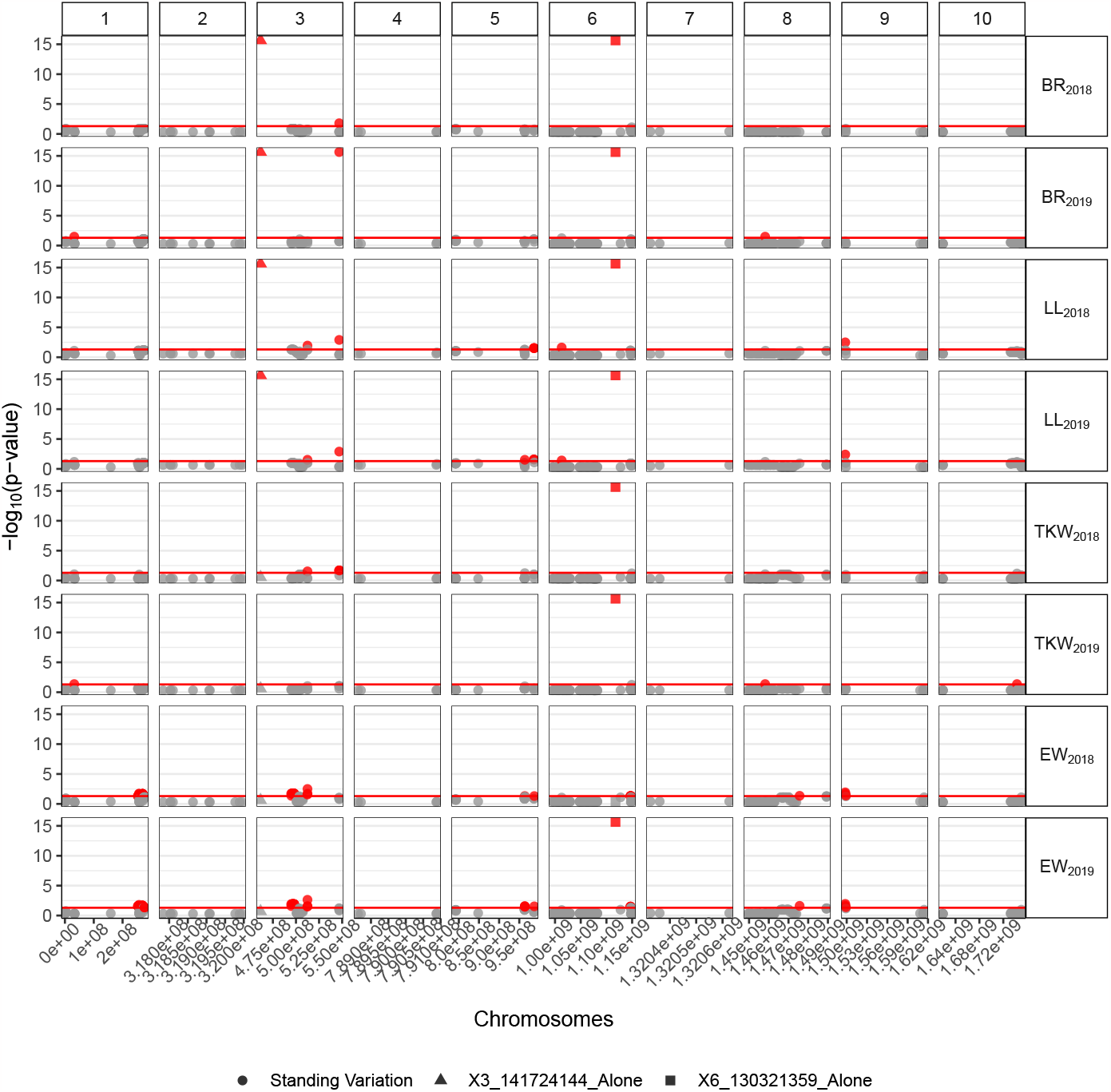
In F252, Manhattan plot for each trait representing the significance of the association of each SNPs (−*log*_10_(*p* − *value*), *FDR* < 5%). Each row of panels corresponds to one trait. Each column of panels corresponds to a chromosome. X-axis indicates the position of SNPs along the chromosomes in bp. Red point (resp. gray point) corresponds to significant (resp. non-significant) association to the trait for the corresponding SNP. Standing variation are represented by a dot, while the two detected mutations are represented respectively by a triangle or a square.

**Figure S27.**
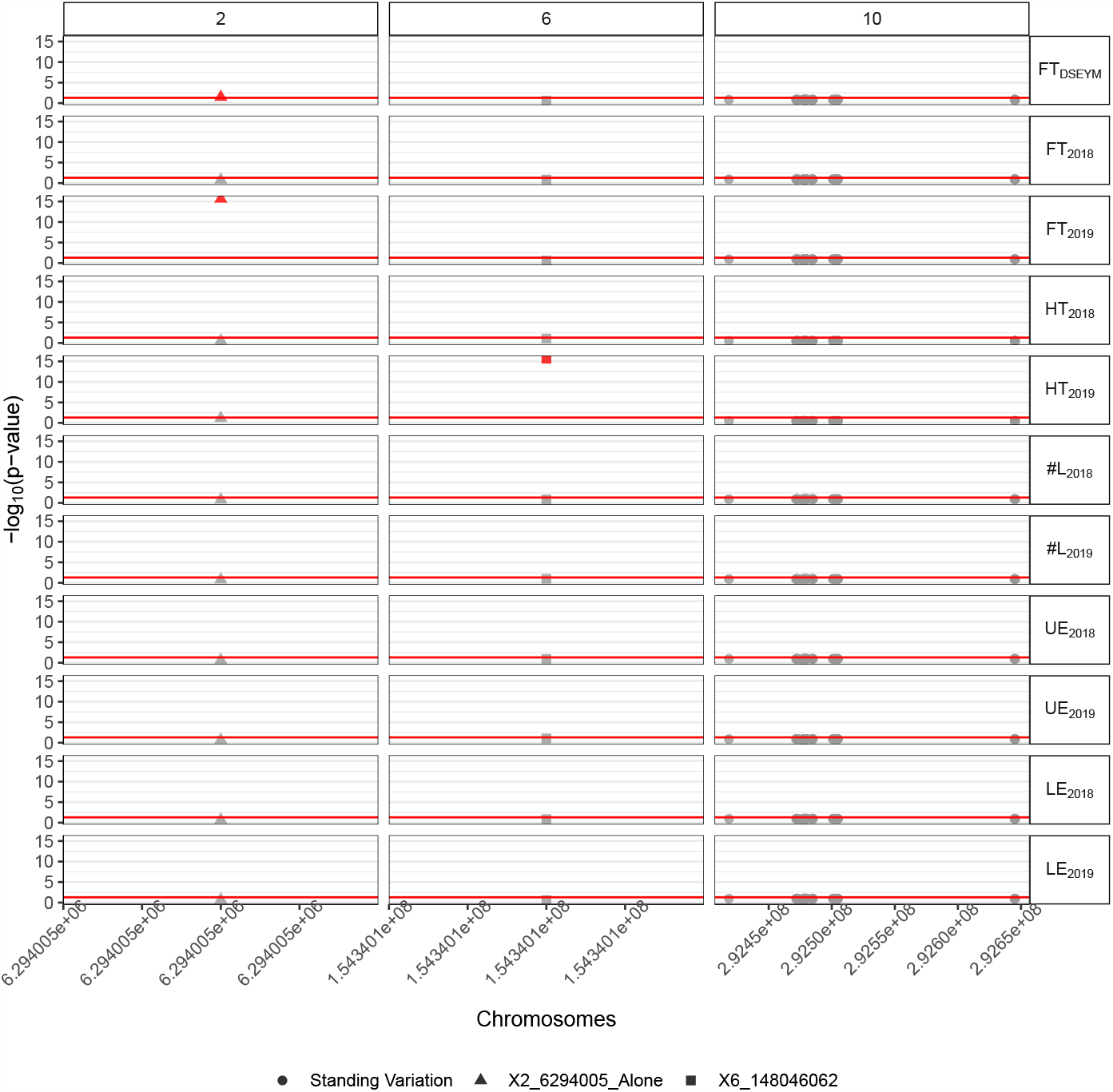
In MBS, Manhattan plot for each trait representing the significance of the association of each SNPs (−*log*_10_(*p* − *value*), *FDR* < 5%). Each row of panels corresponds to one trait. Each column of panels corresponds to a chromosome. X-axis indicates the position of SNPs along the chromosomes in bp. Red point (resp. gray point) corresponds to significant (resp. non-significant) association to the trait for the corresponding SNP. Standing variation are represented by a dot, while the two detected mutations are represented respectively by a triangle or a square.

**Figure S27.**
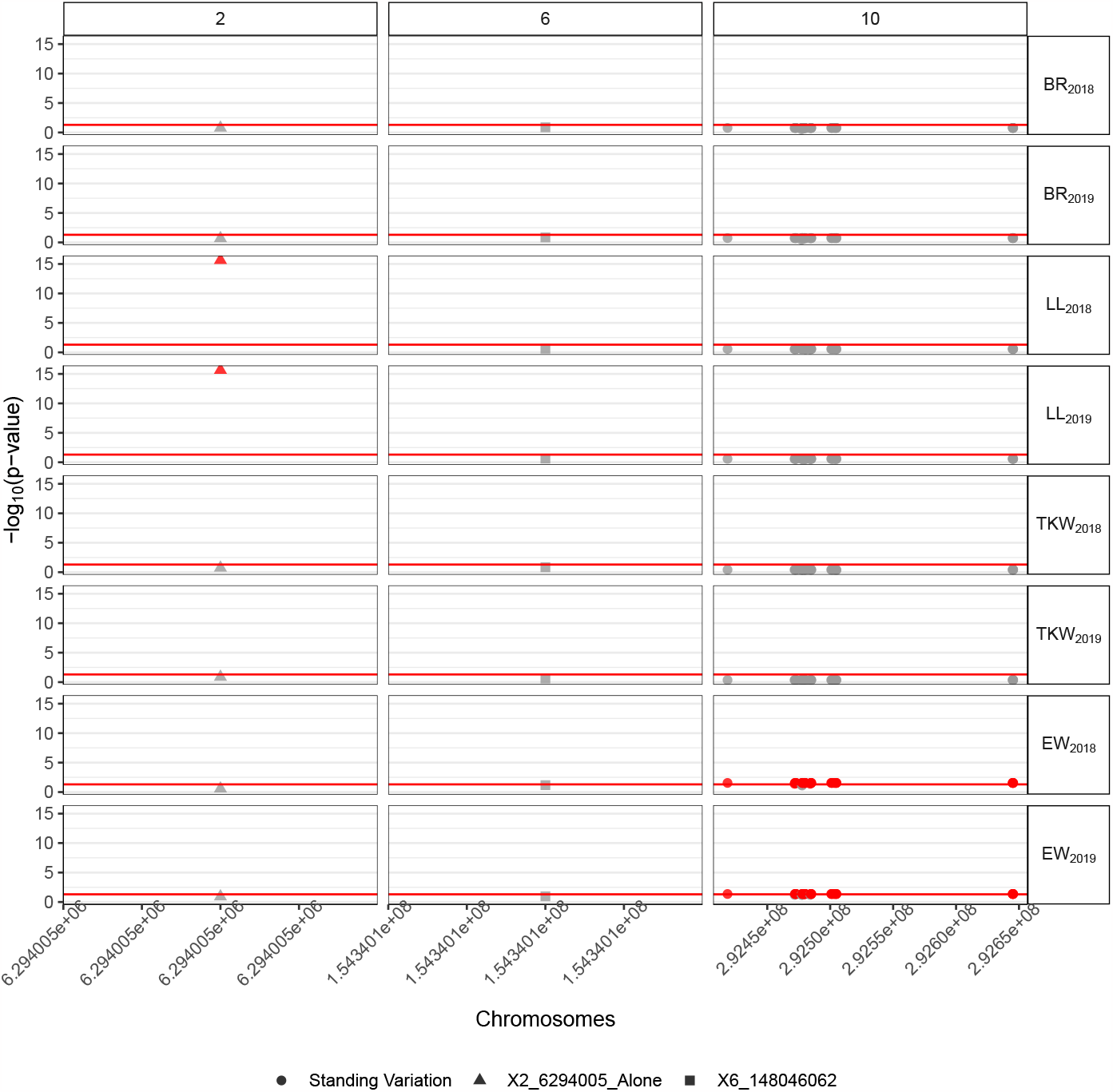
In MBS, Manhattan plot for each trait representing the significance of the association of each SNPs (−*log*_10_(*p* − *value*), *FDR* < 5%). Each row of panels corresponds to one trait. Each column of panels corresponds to a chromosome. X-axis indicates the position of SNPs along the chromosomes in bp. Red point (resp. gray point) corresponds to significant (resp. non-significant) association to the trait for the corresponding SNP. Standing variation are represented by a dot, while the two detected mutations are represented respectively by a triangle or a square.

**Table S7.** Significant trait association (allele effect) in 2018, 2019 DSECG as well as DSEYM in MBS. The first columns provide information on the chromosome, position, gene id and knwon description.

| Inbred | CHROM | POS | Gene id | Gene function | 2018 | 2019 | DSEYM |
| --- | --- | --- | --- | --- | --- | --- | --- |
| MBS | 2 | 6,294,005 | Zm00001d002125 | nuclear RNA polymerase D2/E2 | LL(1.06) | FT(0.73); LL(1.14) | FT(1.43) |
| MBS | 6 | 148,046,062 | Zm00001d038104 | Mannosylglycoprotein endo-beta-mannosidase | . | HT(1.05) | . |
| MBS | 10 | 138,078,378 | Zm00001d026084 | IQ-domain 23 | EW(0.6) | EW(-0.65) | . |
| MBS | 10 | 138,131,658 | Zm00001d026088 | Outer cell layer2 | EW(0.62) | . | . |
| MBS | 10 | 138,131,894 | Zm00001d026088 | Outer cell layer2 | EW(0.6) | EW(-0.65) | . |
| MBS | 10 | 138,132,103 | Zm00001d026088 | Outer cell layer2 | EW(0.6) | EW(-0.65) | . |
| MBS | 10 | 138,132,566 | Zm00001d026088 | Outer cell layer2 | EW(0.6) | EW(-0.65) | . |
| MBS | 10 | 138,132,599 | Zm00001d026088 | Outer cell layer2 | EW(0.6) | EW(-0.66) | . |
| MBS | 10 | 138,132,898 | Zm00001d026088 | Outer cell layer2 | EW(0.6) | EW(-0.65) | . |
| MBS | 10 | 138,137,201 | Zm00001d026088 | Outer cell layer2 | EW(0.61) | . | . |
| MBS | 10 | 138,137,451 | Zm00001d026088 | Outer cell layer2 | EW(0.6) | EW(-0.66) | . |
| MBS | 10 | 138,137,960 | Zm00001d026088 | Outer cell layer2 | EW(0.59) | EW(-0.69) | . |
| MBS | 10 | 138,138,044 | Zm00001d026088 | Outer cell layer2 | EW(0.6) | EW(-0.65) | . |
| MBS | 10 | 138,138,046 | Zm00001d026088 | Outer cell layer2 | EW(0.6) | EW(-0.65) | . |
| MBS | 10 | 138,139,518 | Zm00001d026089 | Replication factor C sub-unit 2 | EW(0.6) | EW(-0.65) | . |
| MBS | 10 | 138,139,571 | Zm00001d026089 | Replication factor C sub-unit 2 | EW(0.6) | EW(-0.65) | . |
| MBS | 10 | 138,139,579 | Zm00001d026089 | Replication factor C sub-unit 2 | EW(0.6) | EW(-0.67) | . |
| MBS | 10 | 138,139,659 | Zm00001d026089 | Replication factor C sub-unit 2 | EW(0.6) | EW(-0.66) | . |
| MBS | 10 | 138,140,091 | Zm00001d026089 | Replication factor C sub-unit 2 | EW(0.6) | EW(-0.65) | . |
| MBS | 10 | 138,140,299 | Zm00001d026089 | Replication factor C sub-unit 2 | EW(0.58) | EW(-0.65) | . |
| MBS | 10 | 138,144,078 | Zm00001d026089 | Replication factor C sub-unit 2 | EW(0.62) | . | . |
| MBS | 10 | 138,144,542 | Zm00001d026089 | Replication factor C sub-unit 2 | EW(0.6) | EW(-0.65) | . |
| MBS | 10 | 138,144,756 | Zm00001d026089 | Replication factor C sub-unit 2 | EW(0.6) | EW(-0.65) | . |
| MBS | 10 | 138,144,795 | Zm00001d026089 | Replication factor C sub-unit 2 | EW(0.6) | EW(-0.65) | . |
| MBS | 10 | 138,144,976 | Zm00001d026089 | Replication factor C sub-unit 2 | EW(0.59) | EW(-0.65) | . |
| MBS | 10 | 138,160,754 | Zm00001d026091 | E3 ubiquitin-protein ligase | EW(0.6) | EW(-0.65) | . |
| MBS | 10 | 138,161,139 | Zm00001d026091 | E3 ubiquitin-protein ligase | EW(0.6) | EW(-0.65) | . |
| MBS | 10 | 138,161,310 | Zm00001d026091 | E3 ubiquitin-protein ligase | EW(0.6) | EW(-0.66) | . |
| MBS | 10 | 138,162,346 | Zm00001d026091 | E3 ubiquitin-protein ligase | EW(0.6) | EW(-0.65) | . |
| MBS | 10 | 138,163,421 | Zm00001d026091 | E3 ubiquitin-protein ligase | EW(0.6) | EW(-0.65) | . |
| MBS | 10 | 138,163,611 | Zm00001d026091 | E3 ubiquitin-protein ligase | EW(0.6) | EW(-0.65) | . |
| MBS | 10 | 138,164,662 | Zm00001d026091 | E3 ubiquitin-protein ligase | EW(0.6) | EW(-0.65) | . |
| MBS | 10 | 138,164,821 | Zm00001d026091 | E3 ubiquitin-protein ligase | EW(0.59) | EW(-0.67) | . |
| MBS | 10 | 138,164,834 | Zm00001d026091 | E3 ubiquitin-protein ligase | EW(0.6) | EW(-0.65) | . |
| MBS | 10 | 138,304,922 | Zm00001d026094 | Heat stress transcription factor B-2b | EW(0.6) | EW(-0.65) | . |
| MBS | 10 | 138,304,960 | Zm00001d026094 | Heat stress transcription factor B-2b | EW(0.6) | EW(-0.65) | . |
| MBS | 10 | 138,305,007 | Zm00001d026094 | Heat stress transcription factor B-2b | EW(0.6) | EW(-0.65) | . |
| MBS | 10 | 138,305,241 | Zm00001d026094 | Heat stress transcription factor B-2b | EW(0.6) | EW(-0.65) | . |
| MBS | 10 | 138,305,316 | Zm00001d026094 | Heat stress transcription factor B-2b | EW(0.6) | EW(-0.65) | . |

**Table S8.**
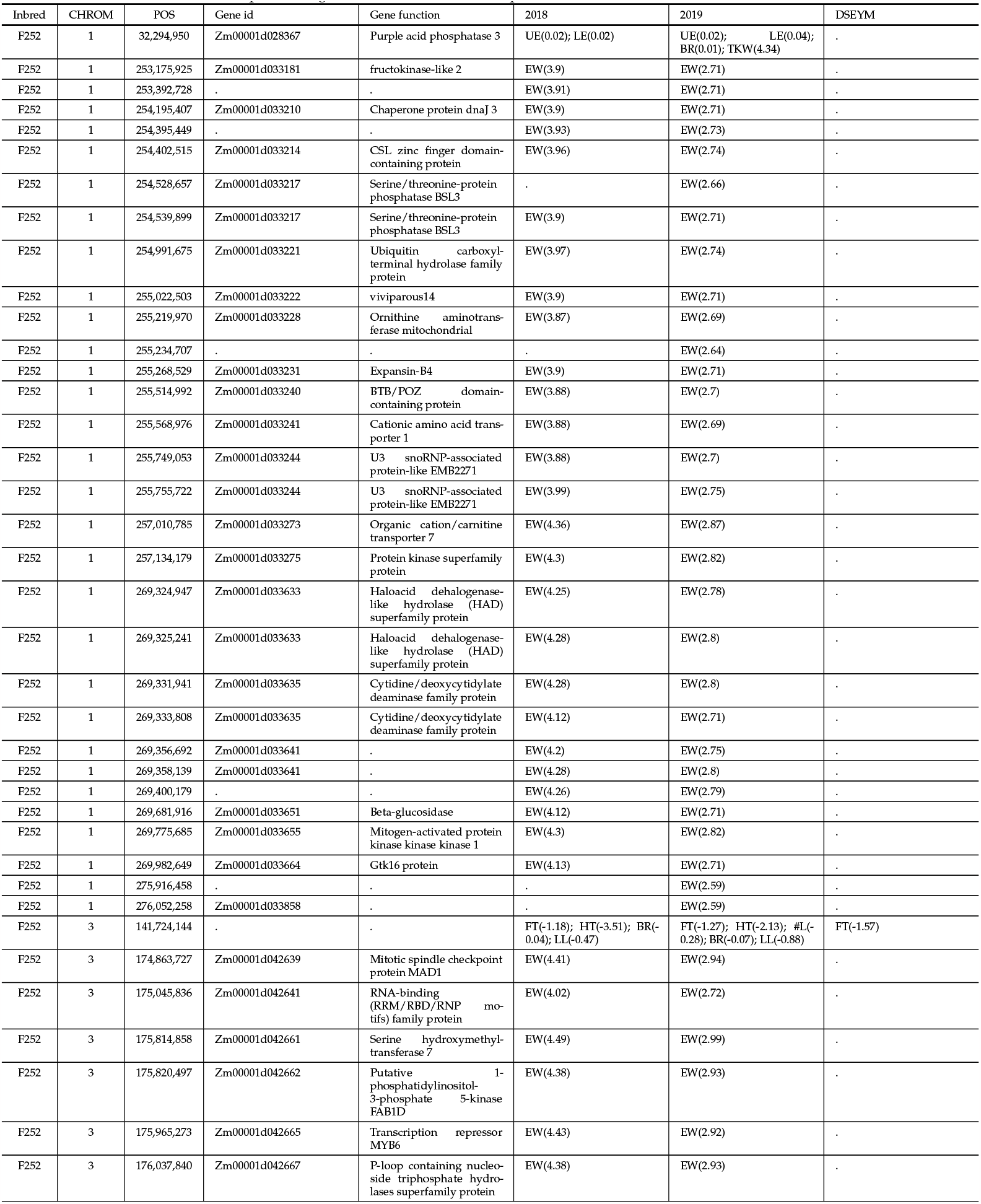
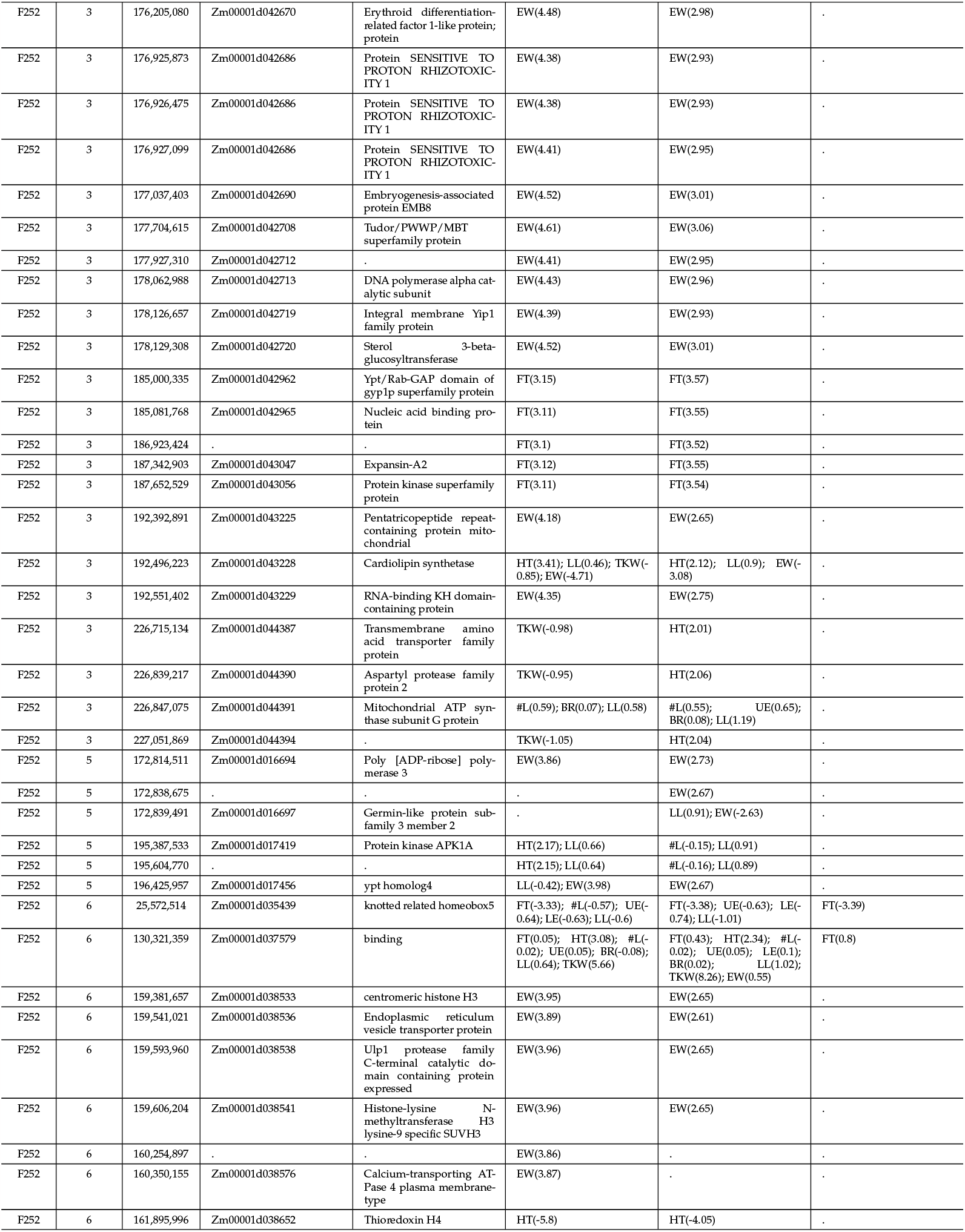
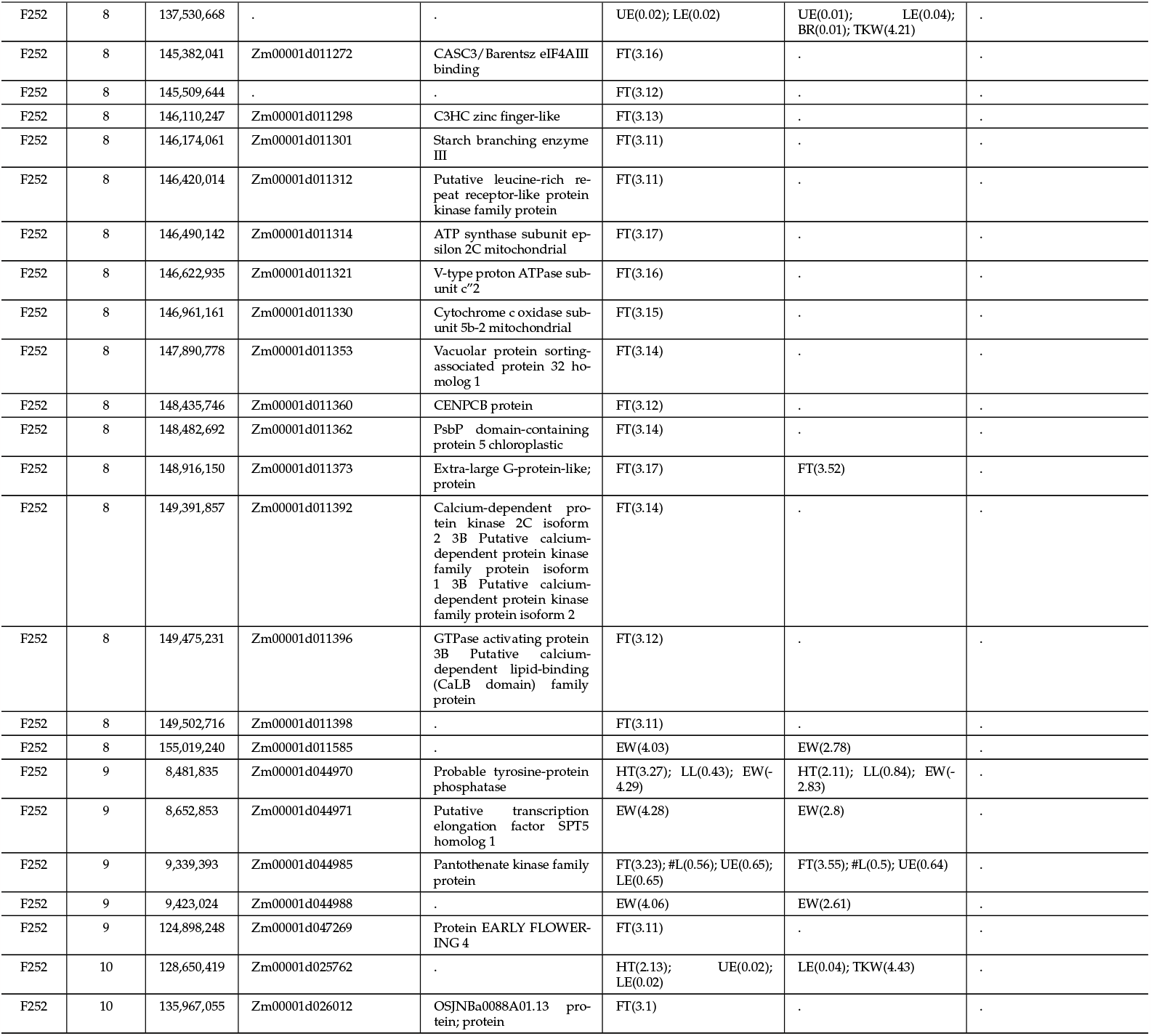
Significant trait association (allele effect) in 2018, 2019 DSECG as well as DSEYM in F252. The first columns provide information on the chromosome, position, gene id and knwon description.

